# Spatiotemporal dynamics of non-ecological speciation in rubyspot damselflies (*Hetaerina* spp.)

**DOI:** 10.1101/2024.12.07.627243

**Authors:** C. Patterson, A. Brennan, H. Cowling, A. González-Rodríguez, G.F. Grether, L. Mendoza Cuenca, M. Springer, Y. M. Vega-Sanchez, J. Drury

**Affiliations:** Department of Biosciences, Durham University, Stockton Road, Durham, United Kingdom DH1 3LE; Instituto de Investigaciones en Ecosistemas y Sustentabilidad, Universidad Nacional Autónoma de México, Morelia, México; Department of Ecology & Evolutionary Biology, University of California, Los Angeles, USA; Facultad de Biología, Universidad Michoacana de San Nicolás de Hidalgo, Morelia, México; Escuela de Biología and Museo de Zoología, Centro de Investigación en Biodiversidad y Ecología Tropical, Universidad de Costa Rica, San José, Costa Rica

**Keywords:** Odonata, Zygoptera, ddRAD, speciation, population genomics

## Abstract

Non-ecological speciation is a common mode of speciation which occurs when allopatric lineages diverge in the absence of pronounced ecological differences. Yet, relative to other speciation mechanisms, non-ecological speciation remains understudied. Numerous damselfly clades are characterized as non-adaptive radiations (the result of several rounds of non-ecological speciation), but there are few damselfly lineages for which we have a detailed understanding of the spatiotemporal dynamics of divergence. Recent phylogeographic analyses demonstrate that American rubyspot damselflies (*Hetaerina americana* sensu lato) actually comprise at least two cryptic lineages that coexist sympatrically across most of Mexico. To broaden our understanding of the dynamics of diversification to other rubyspot lineages, we investigated the phylogeographic history of smoky rubyspot damselflies (*Hetaerina titia*) using genomic data collected across Central and North America. Unexpectedly, we found evidence of reproductive isolation between the highly genetically differentiated Pacific and Atlantic lineages of *H. titia* in a narrow secondary contact zone on the Isthmus of Tehuantepec, Mexico. We then fit models of historical demography to both *H. americana* sensu lato and *H. titia* to place these comparisons in a temporal context. Our findings indicate that Pacific and Atlantic lineages of *H. titia* split more recently than the broadly sympatric lineages within *H. americana* sensu lato, supporting key assumptions of the non-ecological speciation model and demonstrating that these two pairs of sister lineages are at different stages of the speciation cycle.

## Resumen

La especiación no-ecológica es un modo común de especiación que ocurre cuando los linajes alopátricos divergen sin diferencias ecológicas pronunciadas. Sin embargo, comparado con otros mecanismos de especiación, la especiación no-ecológica sigue siendo poco estudiada. Varios clados de caballitos del diablo se han caracterizado como radiaciones no-adaptativas (es decir, el resultado de varios ciclos de especiación no-ecológica), pero hay pocos ejemplos de caballitos del diablo para los cuales tenemos un conocimiento detallado de la dinámica espaciotemporal de la divergencia. Análisis filogeográficos recientes demuestran que el caballito escarlata común (*Hetaerina americana* sensu lato) en realidad consiste en al menos dos linajes crípticos que coexisten simpátricamente a lo largo de México. Para ampliar nuestra comprensión de la dinámica de la diversificación a otros linajes de los caballitos escarlatas, investigamos patrones filogeográficos del caballito escarlata de alas ahumadas (*Hetaerina titia*) utilizando datos genómicos de individuos de América Central y del Norte. Inesperadamente, encontramos evidencia de que las poblaciones altamente diferenciadas genéticamente del Pacífico y el Atlántico de *H. titia* exhiben aislamiento reproductivo en un punto de contacto secundario en el Istmo de Tehuantepec en México. Además, modelamos la demografía histórica tanto de *H. americana* sensu lato como de *H. titia* para compararlas en un contexto temporal. Descubrimos que las linajes del Pacífico y el Atlántico de *H. titia* se dividieron más recientemente que los linajes hermanos ampliamente simpátricos de *H. americana* sensu lato, lo cual apoya las suposiciones claves del modelo de especiación no-ecológica y demuestra que estos dos pares de linajes hermanos se encuentran en diferentes etapas del ciclo de especiación no-ecológica.

## Introduction

Speciation—the process by which a split in one lineage leads to two or more reproductively isolated lineages—is a key process contributing to the accumulation of biodiversity on Earth. Much research into the speciation process examines the role of natural selection in driving divergence between lineages via ecological speciation, where species divergence and reproductive isolation are underpinned by adaptation to different ecological niches (Nosil, 2012; Rundle & Nosil, 2005). When sustained over several bursts of speciation, ecological speciation leads to adaptive radiations such as the iconic Galápagos finches, Lake Victoria cichlids, or Greater Antillean anoles, and has been the central focus of evolutionary biologists interested in explaining the origin and accumulation of biodiversity (Schluter, 2000; Simpson, 1944).

However, many speciation events lead to species without discernible ecological differentiation between daughter lineages. An alternative model of speciation is non-ecological speciation (Czekanski-Moir & Rundell, 2019; Gittenberger, 1991), where divergence between species is not primarily driven by natural selection but rather by the accumulation of reproductive isolation after a period of allopatric separation of daughter lineages. Such isolation can result from genomic incompatibles that arise over time from genetic drift (Dion-Côté & Barbash, 2017; Ravinet et al., 2017; Westram et al., 2022) or through divergence in reproductive traits (Arnegard et al., 2010; McEachin et al., 2022; Mendelson et al., 2014; Mendelson & Safran, 2021; Okamoto & Grether, 2013).

Despite the intense research focus on adaptive radiations, most clades have not diversified via adaptive radiation (Czekanski-Moir & Rundell, 2019; Rundell & Price, 2009). When sustained through several bouts of speciation, non-ecological speciation can lead to a radiation characterized by minimal ecological differentiation between clade members, referred to as a non-adaptive radiation (Czekanski-Moir & Rundell, 2019; Gittenberger, 1991; Rundell & Price, 2009). A recent analysis of insular radiations of birds (including several textbook examples) demonstrates that the majority of such radiations are non-adaptive (Illera et al., 2024). Indeed, examples of non-adaptive radiations are abundant (Czekanski-Moir & Rundell, 2019) and likely to increase in frequency as genomics leads to the discovery of new cryptic species (Eme et al., 2018; Struck et al., 2018).

In addition to being common in nature, non-adaptive radiations offer compellingly simplified models for studying the diversification process.. For biodiversity to accumulate in a given region, species must be able to both cooccur (e.g., via dispersal into a common area) and coexist (i.e., experience population growth) in one another’s presence (Weir and Price 2011, Tobias et al. 2020). Non-adaptive radiations provide useful case studies for characterising the circumstances under which sister lineages attain range overlap in the absence of ecological differentiation.

Damselflies (Odonata, suborder Zygoptera) provide several iconic examples of non-adaptive radiations (Wellenreuther & Sánchez-Guillén, 2016). According to the widely accepted conceptual model for diversification in damselflies, diversity accumulates via non-ecological speciation as species come into secondary sympatry after sufficient time has passed in allopatry for divergent lineages to become reproductively isolated via the evolution of species-specific genital morphology (e.g., male claspers and the [pro]thoracic plates of females which come into physical contact with male claspers during mating) (Paulson 1974, Wellenreuther & Sánchez-Guillén 2016). Consistent with this model, sympatric assemblages of congeners often exhibit little ecological differentiation (e.g., *Calopteryx* spp. [(Svensson et al., 2018)], *Ischnura* spp. [(Sánchez-Guillén et al., 2005)], *Enallagma* spp. [(McPeek & Brown, 2000)]). Species do, however, possess reproductive characters that are highly divergent from those of other congeners (e.g., *Calopteryx* spp [Svensson et al., 2010, 2014], *Ischnura* spp. [Sánchez-Guillén, Córdoba-Aguilar, et al., 2014; Sánchez-Guillén et al., 2005], *Enallagma* spp. [McPeek et al., 2009, 2011]). Yet, while these observations support the hypothesis that these damselfly genera are non-adaptive radiations, no study to date has reconstructed the temporal dynamics of reproductive isolation and secondary contact in damselflies.

Here, we investigate whether divergence time predicts the outcome of secondary contact within a subset of damselfly species within the genus *Hetaerina. Hetaerina* damselflies have a crown age estimate of 36.2 million years ago (mya) (Standring et al., 2022) with most species living in sympatry with one or more congeners. There are currently 39 recognised *Hetaerina* species (Garrison, 1990; Standring et al., 2022), but the recent discovery that *Hetaerina americana* sensu lato consists of at least two highly diverged and sympatric cryptic species (now named *H. americana* and *Hetaerina calverti;* Vega-Sánchez et al., 2020, 2024) suggests the number may be higher. The morphology of male claspers is the only way to identify some adult *Hetaerina* species in the field (Vega-Sánchez et al., 2020, 2024). All *Hetaerina* species are lotic habitat (stream, river) specialists and closely resemble one another in morphology, diet, and reproductive behaviour, despite the wide diversity of forms and behaviours present in Odonata (Corbet, 1999). Although *Hetaerina* spp. show moderate levels of climatic and microhabitat differentiation (Grether et al., 2024; McEachin et al., 2022), the ecological and phenotypic similarities between species are more remarkable than the differences considering their ancient divergence. Consequently, *Hetaerina* damselflies likely represent another example of a non-adaptively radiating damselfly clade.

We investigate two geographically widespread lineages of *Hetaerina* from across North and Central America: *H. americana* sensu lato (i.e., the *H. americana* and *H. calverti* species complex) and *H. titia*. *H. calverti* is found in sympatry with both the Northern and Southern lineages of *H. americana* (Vega-Sánchez et al., 2024). *H. titia* exhibits the largest latitudinal range of any *Hetaerina* species, extending from Canada in the north to Panama in the south (Grether et al., 2024; Paulson, 2020). Phylogenies of *H. titia* constructed using mitochondrial and nuclear genes suggest divergence between populations that reside in Pacific and Atlantic drainages (Drury, Anderson, et al., 2019; Drury & Grether, 2014). Together, these taxa offer a window into the process of non-ecological speciation.

Here, we use genome-wide markers from specimens across Central and North America to reconstruct the population-level relationships between distinct lineages within the species currently recognised as *H. americana, H. calverti,* and *H. titia*. We then estimate the divergence times between these lineages to characterise the timescale of isolation and secondary sympatry in a non-ecological radiation.

## Methods

### Sampling and sequencing

Whole organism samples of smoky rubyspot (*Hetaerina titia*) and American rubyspot (*H. americana* sensu lato) damselflies were collected between 2006 and 2021 from across Central and North America, submerged in ≥ 95% ethanol or RNALater (Invitrogen), and stored at ≤ −20°C. For DNA extraction approximately 2 mm^3^ of wing muscle tissue was removed from the thorax and processed using DNeasy Blood and Tissue Kits (Qiagen) following standard manufacture protocols. To generate genome-wide sequence data, we followed double digest restriction enzyme associated DNA (ddRAD) protocols (DaCosta & Sorenson, 2014; Franchini et al., 2017; Peterson et al., 2012). We used the restriction enzymes PstI and EcoR and generated multiplexed libraries by ligating adapters containing a region of four random nucleotides for PCR clone removal. After paired-end 150 bp sequencing on a NovaSeq 6000 (Illumina), we demultiplexed and filtered clones (see Supplementary Material for further details on library prep and the full bioinformatics pipelines is outlined in Supplementary Figure 1 and Supplementary Table 1). In total, we obtained sequence data for 205 individuals of *H. titia*, and 58 individuals of *H. americana* sensu lato from across Central and North America.

### SNP Calling

Individual sequences were mapped to a *H. americana* reference genome (Grether *et al*., 2023) and to a *H. titia* reference genome (Patterson *et al*., 2023) using the Burrow-Wheeler aligner (bwa) *mem* alignment algorithm (Li & Durbin, 2009). Genotype calling was done using *bcftools v1.13* (Danecek et al., 2021; Li, 2011) using the *mpileup* and *call* commands.

The probability of any two samples having the same restriction site at a particular locus decreases with phylogenetic distance. As such, multiple SNP libraries were constructed including varying combinations of species for use in different analyses (Table 1, Supplementary Table 2). Firstly, two SNP libraries were produced that contained all *Hetaerina* samples (*H. americana* sensu lato and *H. titia*) and were mapped to either the draft genome of *H. americana* (Grether et al., 2023) or *H. titia* (Patterson *et al*., 2023). Additionally, four different SNP libraries were constructed, again using both draft genomes, for all *H. americana* sensu lato samples and, separately, all *H. titia* samples. Finally, we also conducted a de novo (reference-free) SNP assembly, to determine if there was any ascertainment bias using draft genomes that were more closely related to either species or population within our sample set. To reduce computation time, we limited the de novo SNP library to 3 of the highest coverage samples from each identified lineage (18 samples in total) from the reference mapped libraries. The de novo pipeline was constructed using ipyrad (Eaton & Overcast, 2020) as outlined in the Supplementary Methods. As recent introgression (< 2 generations) violates assumptions of the phylogenetic and demographic analysis, we created additional sets of SNP libraries excluding the samples from a drainage where preliminary results suggested recent introgression between Atlantic and Pacific population clusters of *H. titia*. The full bioinformatics pipeline is outlined in the Supplementary Methods, and all scripts are available on GitHub (https://github.com/ChristophePatterson/Phylogeography-Hetaerina).

**Table 1.** Overview of all analyses and SNP/loci libraries used for each. Each library consists of different combinations of samples from different species and reads were aligned to the draft genome of *H. americana* (HetAmer1.0, Grether et al., 2023), *H. titia* (HetTit1.0, Patterson et al., 2023) or mapped *de novo*. Each analysis and library are marked as to whether the results are presented in the main text (M) or in the supplementary (S). *Hetaerina americana* sensu lato consists of three distinct lineages including the recently described *H. calverti* (Vega-Sánchez et al., 2024). A breakdown of the number of samples and the SNP/loci number for each analysis is presented in Supplementary Table 2.

|  |  | Population<br>structure | Phylogenies |  | Demographic / divergence<br>times |  |  | Introgress<br>ion |
| --- | --- | --- | --- | --- | --- | --- | --- | --- |
| Species included | Read alignment | sNMF &<br>PCA | RAxM<br>L | SVD<br>Quartets | delimit<br>R | SNAP<br>P | G-Phocs | Introgress |
| *H. americana* sensu lato & *H. titia* | HetAmer1.0 | - | S | S | - | M | - | - |
| *H. americana* sensu lato & *H. titia* | HetTit1.0 | - | S | S | - | M | - | - |
| *H. americana* sensu lato & *H. titia* | *de novo* | - | - | - | - | M | - | - |
| *H. americana* sensu lato | HetAmer1.0 | M | M | S | S | - | S | - |
| *H. americana* sensu lato | HetTit1.0 | S | S | S | S | - | S | - |
| *H. americana* sensu lato | *de novo* | - | - | - | - | - | M | - |
| *H. titia* | HetAmer1.0 | S | S | S | S | - | S | - |
| *H. titia* | HetTit1.0 | M | M | S | S | - | S | M |
| *H. titia* | *de novo* | - | - | - | - | - | M | - |

The resulting vcf files were imported into R using the package vcfR (Knaus & Grünwald, 2017). Further SNP and sample filtering (Supplementary Methods) and conversion of vcf into compatible formats for each analysis software was done using the R packages ape (Paradis & Schliep, 2019), adegenet (Jombart, 2008), and poppr (Kamvar et al., 2014). The total number of samples, species included, number of SNPs/loci, and read alignment methodology used in each analysis are presented in Supplementary Table 2.

### Species delimitation and population structure

To characterise the population structure of *H. americana* sensu lato and *H. titia*, we used the R package LEA (Frichot & François, 2015) to conduct principle component analysis (PCA) and non-negative matrix factorization algorithms (sNMF) for least-squares estimates of ancestry proportions for each sample (Frichot et al., 2014). We restricted the SNPs to those that were biallelic and removed samples that had more than 20% missing data. To maintain equal level of ploidy we removed SNPs mapped to the X chromosome, as *Hetaerina* has an XX/XO sex determination system (Patterson et al., 2023). In sNMF, we tested for a range of ancestral populations (K = 1 to 10) and plotted the mean cross-entropy values for 100 repetitions. We used hierfstat (Goudet, 2005) to calculate F_st_ between each identified cluster.

### Phylogenetic inference

We reconstructed phylogenetic trees for each reference mapped SNP library (Table 1) using RAxML/8.2.12 (Stamatakis, 2014). As RAxML requires homozygous SNPs, we filtered the vcfs to include only homozygous-called sites, then excluded sites that were invariant across individuals after removing samples with greater than 20% missing data. Phylogenies were reconstructed under a general time reversible model (GTR), a gamma distribution of rate heterogeneity, and a Lewis ascertainment correction due to the exclusion of invariant sites (-m = ASC_GTRGAMMA) (Devitt et al., 2019; Lozier et al., 2016).

We also reconstructed phylogenies in SVDquartets (Chifman & Kubatko, 2014) in PAUP* (Wilgenbusch & Swofford, 2003). Heterozygous sites, which are compatible with SVDquartet analysis, were retained (Supplementary Table 2). We calculated the SVD score of 100,000 unrooted 4-‘taxa’ trees (quartets) and to infer the optimal phylogenetic relationship between the samples for each quartet, we used the Quartet FM method (Reaz et al., 2014). We then constructed a consensus tree by repeating the process 100 times to produce bootstrap support values for each tree node determined by the percentage of times the node was part of the consensus topology of the tree.

### Testing for migration between lineages

To test the assumption of the non-ecological speciation hypothesis that little to no migration occurs between diverged lineages, we used the R package delimitR (Smith & Carstens, 2020). delimitR uses site frequency spectrums (SFS) built from a SNP data set to predict the most likely demographic history for several potential populations or species. It then uses fastsimcoal2 (v2.6) (Excoffier et al., 2013, 2021) to simulate SFS for each specified demographic scenario under a range of priors and builds a random forest classifier to estimate the most likely demographic scenario for the observed data. For population clusters of *H. titia* and for *H. americana* sensu lato, we simulated each valid combination of several demographic scenarios with and without migration between populations (Supplementary Figure 2). We simulated each scenario using broad, uniform priors (Supplementary Methods).

Empirical SFS were calculated using the package easySFS (https://github.com/isaacovercast/easySFS) which builds off the dadi.Spectrum class from the software ∂a∂i (Gutenkunst et al., 2009). To take into account missing SNPs, which are inherent to ddRAD data, we projected down the SFS to maximise the number of segregating sites following Gutenkunst et al. (2009).

### Divergence time estimation

To place divergence among *Hetaerina* lineages within the broader context of the speciation cycle, we estimated the divergence times of population clusters using two approaches. Firstly, we ran the Bayesian coalescent analysis SNAPP implemented within the programme Beast v2.7.5 (Bouckaert et al., 2019). Due to computational constraints, we restricted the analysis to four individuals per cluster identified by sNMF (24 individuals in total) with the highest SNP coverage from each distinct ancestral clustering identified by sNMF. We then removed SNPs that were either no longer polymorphic between the selected samples, genotyped in less than one individual from each population, or mapped to X chromosome. We used previous estimated divergence times from Standring et al. (2022) as priors by secondary calibration for divergence time between *H. titia* and *H. americana* senso lato (mean = 33.08 million years ago (mya), standard deviation = 5.53 mya) and for the divergence of *H. americana* and *H. calverti* (mean = 3.76 mya, standard deviation = 1.87 mya). We used a starting tree that had the same relationships identified in RAxML and SVDquartets for each of the clusters and ran MCMC for 1,000,000 generations, sampling every 500 iterations. A SNAPP configuration file was created using a custom R script and the ruby script from https://github.com/mmatschiner/snapp_prep. We assessed the convergence using tracer and calculated the maximum clade credibility tree, with a 10% burn in removal, using TreeAnnotator v2.7.5 (Bouckaert et al., 2019).

For an alternative estimate of divergence times not based on a secondary calibration, we fit models of historical demography using G-PhoCS (Gronau et al., 2011) which uses a Bayesian coalescent approach. We present parameter estimates for the demographic models that were best supported by delimitR. We ran G-PhoCS using loci mapped using heterospecific draft genome, conspecific draft genome, and loci mapped *de novo* for each species. We converted mutation rate-scaled parameter estimates of G-PhoCS into the number of diploid individuals and the number of years using 2.8e-9 mutations per base pair per generation (Keightley et al., 2014). We converted generations to years using an estimated generation time of one year. We present results from G-PhoCS using the loci mapped de novo in the main text as these libraries minimise ascertainment bias (see Supplementary Methods for further detail).

### Investigating a potential secondary contact zone

Preliminary analysis identified an individual of *H. titia* with admixed ancestry from a site on the Isthmus of Tehuantepec, in Mexico. To determine the number of generations since the putative hybridisation event and see if any other individuals had admixed ancestry, we ran a hybridisation analysis using the R package introgress (Gompert & Buerkle, 2010). We subset our data to samples from sites in and around the Isthmus of Tehuantepec. We calculated the allele frequency for each SNP for both Pacific and Atlantic populations, excluding samples from the drainage where the putative hybrid was identified. We then subset our dataset to 914 autosomal SNPs and 19 sex linked SNPs that had an allele frequency difference greater than 0.8 between the Pacific and Atlantic, in line with DeRaad *et al*. (2022). We then assigned each allele to a “parental” Pacific or Atlantic genotype and calculated both the percentage of Pacific and Atlantic alleles carried by each sample (the hybridisation index), and the average autosomal heterozygosity across all highly divergent SNPs (the multi-allele heterozygosity) for each sample.

## Results

Nearly all analyses, using all different combinations of libraries produced comparable results. For brevity, we summarise the results in the main text and present the result for each individual library in the Supplementary Material. For an overview of which SNP/loci libraries were used in each analysis, see Table 1.

We retained sequence data for 259 to 263 samples of *H. americana* sensu lato and *H. titia* which had between 519 to 609 SNPs with adequate genotyping across all samples (Supplementary Tables 2-3). For SNPs libraries that only included *H. americana* sensu lato, we retained sequencing for 58 samples with 1,816 to 5,259 SNPs. For SNPs libraries which only included *H. titia*, we retained sequencing for between 205 to 207 samples with 1,122 to 3,819 SNPs depending on read alignment methodology. Across all SNP libraries, we attained an average coverage of between 50-53x and a median missing genotype rate of around 1.6% to 1.9%.

### Hetaerina titia population structure

sNMF admixture analyses and principal component analyses both identified three distinct clusters in *H. titia* (Figure 1, 2, Supplementary Fig. 3): (1) a Caribbean and Southern Gulf of Mexico cluster, (2) a Northern Gulf of Mexico and Atlantic cluster, and (3) a Pacific Coast cluster (Figure 1-2, Supplementary Figures 3-7). Hereafter, we refer to these three clusters as the Southern Atlantic *H. titia* cluster, the Northern Atlantic *H. titia* cluster, and the Pacific *H. titia* cluster, respectively. The pairwise F_st_ values between the three groups indicate high levels of differentiation. Using the 3,819 SNPs mapped to *H. titia* draft genome, the F_st_ was 0.818 between the Pacific and Northern Atlantic, 0.730 between the Pacific and Southern Atlantic, and 0.521 between the Northern and Southern Atlantic. We identified one sample with extensive admixture between the Pacific and Southern Atlantic clusters from site CUAJ01 in Cuajinicuil, Oaxaca (16°47’24.00′N, 95°0’36.00′W) on the Gulf slope of Isthmus of Tehuantepec (Mexico).

**Figure 1.**
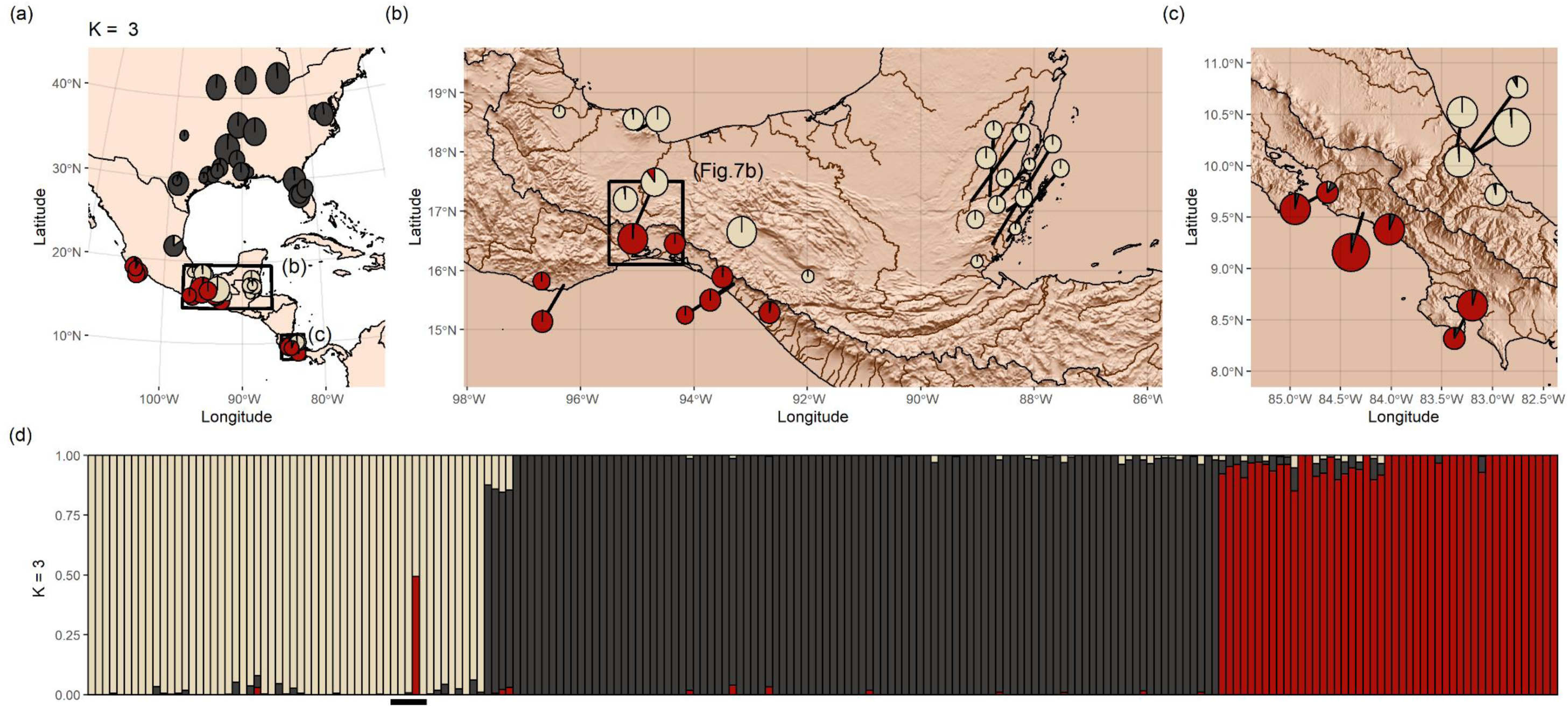
Ancestry estimates for 205 *Hetaerina titia* with a dataset of 3,819 unlinked biallelic autosomal SNPs. SNPs were generated by mapping ddRAD reads to the draft genome of *H. titia*. LEA was run for 20 repetitions and an alpha value of 100. (a) The mean estimate of ancestry proportion for all samples within each sample site of *H. titia* across Central and North America, (b) Isthmus of Tehuantepec and Belize, and (c) Costa Rica. Within panels a, b, and c, the area of each pie chart is proportional to the number of samples from each site and then coloured by the mean proportion of estimated ancestry across all samples from each site. (d) Estimate of ancestry for each individual. Samples are ordered by drainage, then country, and then latitude. Rivers and drainage basins from Hydrosheds. Topography data from the R package elevatr. The black boxes shown in panel (a) are the bounding areas for panels (b) and (c). The black box in panel (b) is the bounding box for Figure 7b and the five samples from the site with an identified hybrid individual are underlined in panel (d).

**Figure 2.**
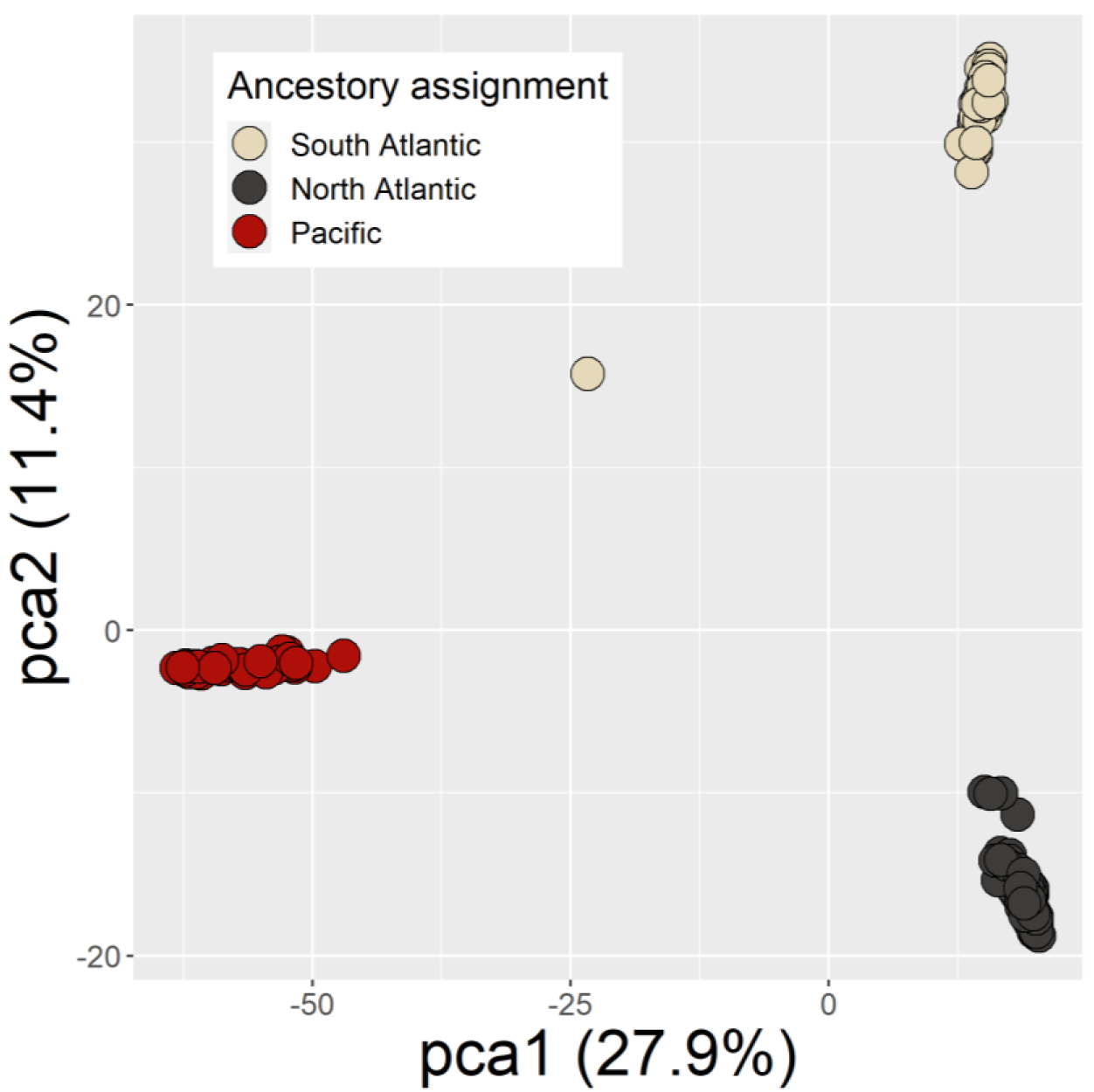
Principal component analysis of 205 *Hetaerina titia* with a dataset of 3,819 unlinked biallelic autosomal SNPs that were generated by mapping ddRAD reads to the draft genome of *H. titia*. Percentages indicate how much variation is explained by each component and colour indicates the highest assigned ancestry population from sNMF for each individual. The single point directly between the main Pacific and Atlantic cluster is the putative F_1_ hybrid.

### Hetaerina americana sensu lato population structure

Consistent with previous work conducted on a different set of specimens with different restriction enzymes (Vega-Sánchez et al., 2020, 2024), analyses of *H. americana* sensu lato also grouped samples into three distinct clusters (Supplementary Figure 8-13). *H. calverti* forms one cluster, and *H. americana* is split into two distinct clusters—a Northern population in the continental United States, and a Southern population, found on both the Gulf and Pacific slopes of Mexico. We refer to these lineages as Northern *H. americana* and Southern *H. americana* going forward. Using SNPs mapped to the draft genome of *H. americana,* pairwise F_st_ values, between the identified groups were 0.833 (*H. calverti* vs Northern *H. americana*), 0.791 (*H. calverti* vs Southern *H. americana*), and 0.699 (Northern vs Southern *H. americana*).

### Phylogenetic inference of Hetaerina

In agreement with population structure analyses, populations of *H. titia* which reside in drainages that flow into the Atlantic, including the Gulf of Mexico and the Caribbean, are more closely related to each other than populations that reside in drainages that flow into the Pacific. The Atlantic lineage is split into two groups (1) samples that originated from the continental United States and the most Northern sample site in Mexico, and (2) the remaining samples from Mexico, Belize, and Costa Rica (Figure 3). Within the Pacific *H. titia* lineage, there are three distinct groups, one group from Costa Rica and two separate Central and Southern lineages in Mexico (Figure 3, Supplementary Figure 14).

**Figure 3.**
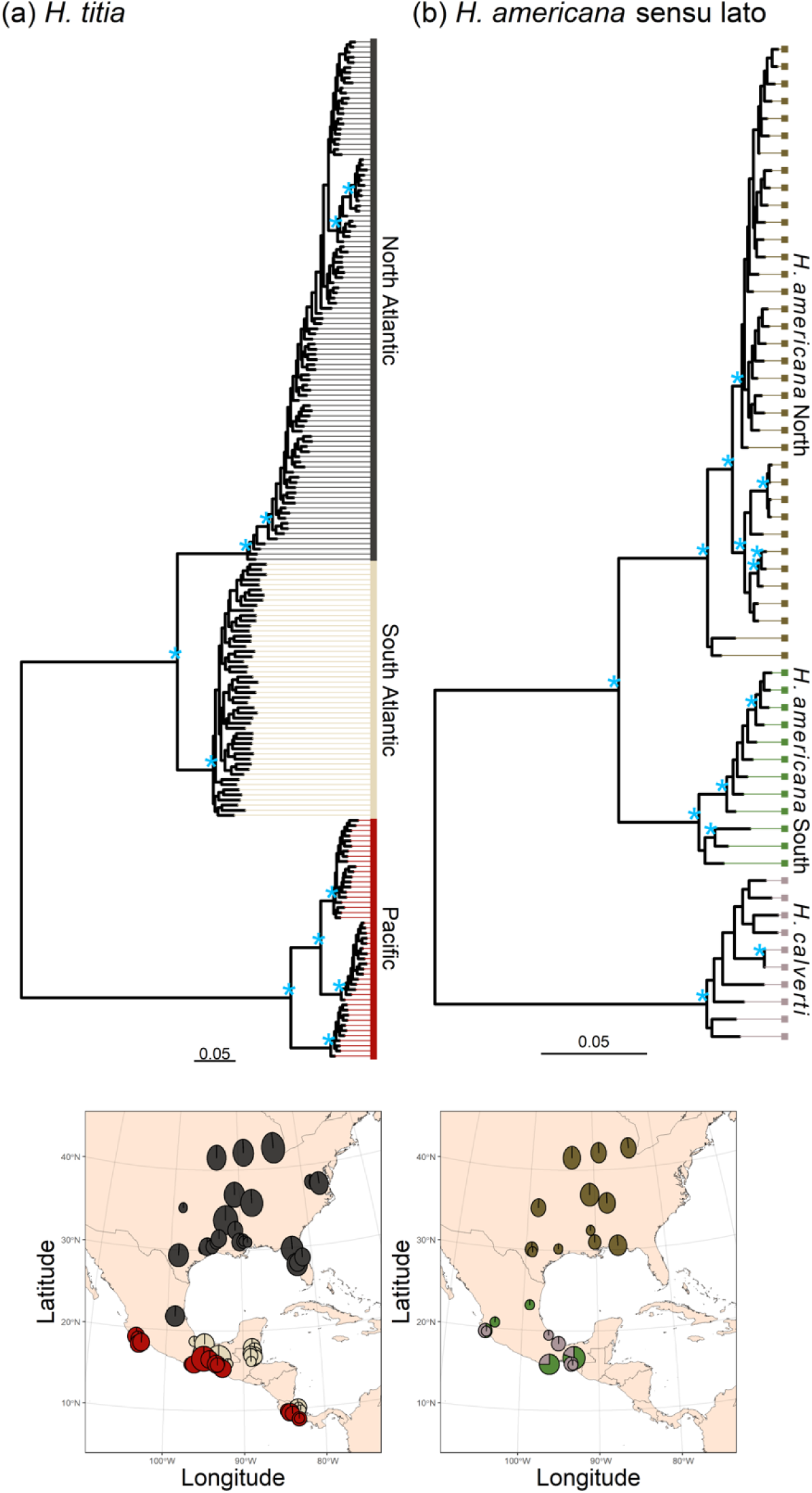
The maximum likelihood tree for (a) *Hetaerina titia* and (b) *Hetaerina americana* sensu lato. Calculated using RAxML with 3,020 SNPs for *H. titia* and 3,949 SNPs for *H. americana* and *H. calverti* and mapped onto the genome of *H. titia.* Scale bar indicates the mean number of substitutions per SNP site. Due to exclusion of invariant sites and differences in the total number of SNPs used in each analysis, scale bars should not be used to compare phylogenetic distances within *H. titia* to distance within *H. americana* sento lato. The nodes marked with a blue star “*” indicate a bootstrap support value (out of 100) of greater than 95%. The tree tips are coloured according to the species and the max sNMF ancestry assignment (K=3). The geographical location of each sample is shown in the bottom two maps. Each pie chart shows the number of samples assigned to each ancestry cluster from each sample site, split between *H. titia* and *H. americana* sensu lato (*H. americana*/*calverti*).

Within the *H. americana* sensu lato lineage there is a distinct split between populations in continental United States and populations in Mexico. Unlike *H. titia* lineages, neither *H. americana* sensu lato lineage is restricted to either Pacific or Atlantic drainages— the *H. americana* south and *H. calverti* lineages ranges broadly overlap and are commonly found coexisting sympatrically (Supplementary Figure 10-11).

Key inferences from SVDquartet analyses were qualitatively similar to those derived from RAxML (Supplementary Figure 16-17).

### Tests for migration between lineages

The best-supported demographic scenarios for *H. titia* suggest that Pacific and Atlantic lineages are completely isolated, with no evidence of ancient or contemporary migration between them. The best fit demographic scenarios did contain ancestral—but not contemporary—gene flow between the Northern and Southern Atlantic lineages (Model 13 in Supplementary Figure 2, receiving 85.3% of support). There was some support for the demographic scenarios with no migration, neither contemporary nor ancestral, between any lineage (Model 5 in Supplementary Figure 2, receiving 14.3% of support). The out of the bag error rate varied among *H. titia* demographic scenarios but was low for models 5 and 13 (20% and 10%). Furthermore, incorrect classifications of models 5 and 13 were limited to the alternative of these two scenarios. No other demographic scenarios for *H. titia* received more than 2% support (Supplementary Table 4).

Similarly, for *H. americana* and *H. calverti*, models suggest no ancient or contemporary migration between *H. americana* and *H. calverti*. The most favoured model had three separate lineages with no migration (Model 5 in Supplementary Figure 3, receiving 72.6% of support), followed by a model with three lineages with isolation with ancient migration between the Northern and Southern lineages of *H. americana* (Model 13 in Supplementary Figure 3, receiving 13.3% of support). The out of the bag error rate for *H. americana* and *H. calverti* demographic scenarios varied but were again low for models 5 and 13 (26% and 11%, respectively) and incorrect classifications were limited to the alternative of these two scenarios. No other demographic scenarios received any support.

### Divergence times

SNAPP analysis using the SNPs mapped to the *H. americana* genome, the *H. titia* genome, and mapped de novo, converged on the same tree and estimates of divergence times between each species and sub-population overlapped (Figure 4, Supplementary Figures 18-20). Based on the de novo SNP data, the divergence time between *H. titia* and *H. americana* sensu lato was estimated to be 24.5 mya (95% highest posterior density [HPD] 15.67-33.59 mya). The divergence between *H. calverti* and *H. americana* was estimated to be 6.83 mya (HPD 4.23-9.17 mya). SNAPP analysis also identified relatively distant dates for the divergence between the sub-populations within *H. titia* and *H. americana*. Populations of *H. titia* that reside in Atlantic drainages were estimated to have diverged from populations in the Pacific 3.74 mya (HPD 2.18-5.59 mya). The two lineages of *H. titia* that reside within Atlantic drainages separated at an estimated 1.11 mya (HPD 0.52-1.75 mya). The two identified lineages of *H. americana* diverged 3.25 mya (HPD 1.79-4.90 mya). Across the posterior distribution of trees of the de novo SNAPP run, the split between Pacific and Atlantic *H. titia* was younger than the split between *H. americana* and *H. calverti* (mean 3.09 million years, HPD +1.48 to +4.86 million years). In the SNAPP analysis using the SNPs mapped to the draft genome of *H. americana*, in 99.8% of posterior distribution trees, the split between Pacific and Atlantic *H. titia* was younger than the split between *H. americana* and *H. calverti* (mean +2.38 million years HPD +0.90 to +3.95 million years). Using the SNPs mapped to the draft genome of *H. titia*, in 75.3% of posterior distribution trees, the split between Pacific and Atlantic *H. titia* was younger than the split between *H. americana* and *H. calverti* (mean +0.5 million HPD −1.17 to +1.92 million years).

**Figure 4.**
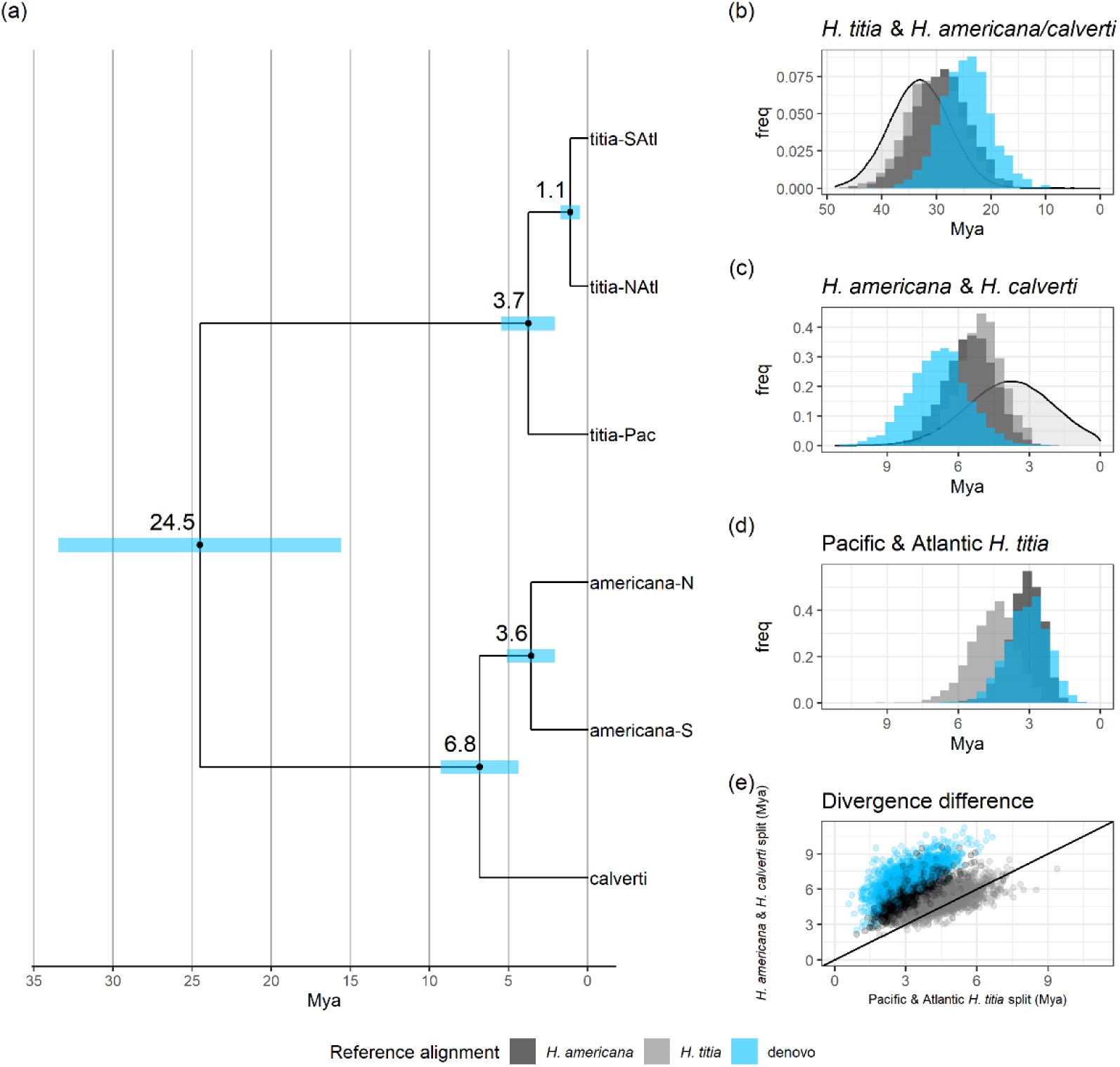
(a) Estimates of divergence dates (million years ago - mya) between populations of Pacific *Hetaerina titia* (titia-Pac), Atlantic *H. titia* (titia-NAtl and titia-SAtl), *Hetaerina americana* (americana-N and americana-S), and *Hetaerina calverti* (calverti) calculated using SNAPP analysis in Beast. Node labels indicate the mean estimated divergence date with 95% highest posterior density in blue. All branches had a posterior distribution of 1. Tree plotted in R using the packages *treeio* and *ggtree.* Input data was 552 autosomal SNPs called using a de novo method of SNP calling (ipyrad). (b-c) The prior and posterior distribution (where applicable) of divergence times between the major lineages. The three different histograms denote the posterior distribution of the divergence times using three different SNP datasets, those mapped the draft genome of *H. americana* (dark grey), mapped the draft genome of *H. titia* (grey), and *de novo* SNP calling. The prior distribution, where applicable, is denoted by the grey density distribution (e) Comparison between the divergence times of *H. americana* and *H. calverti* and Atlantic and Pacific *H. titia*. Each point is the divergence times from a tree in the posterior distribution, the black line indicates values where the divergence time is between the lineages are equal.

Divergence times estimated by G-PhoCS were generally more recent than those estimated by SNAPP (Figure 5, Supplementary Figures 21-22), but in all cases the divergence between *H. americana* and *H. calverti* was estimated as approximately twice as old as the split between Pacific and Atlantic clusters of *H. titia*.

**Figure 5.**
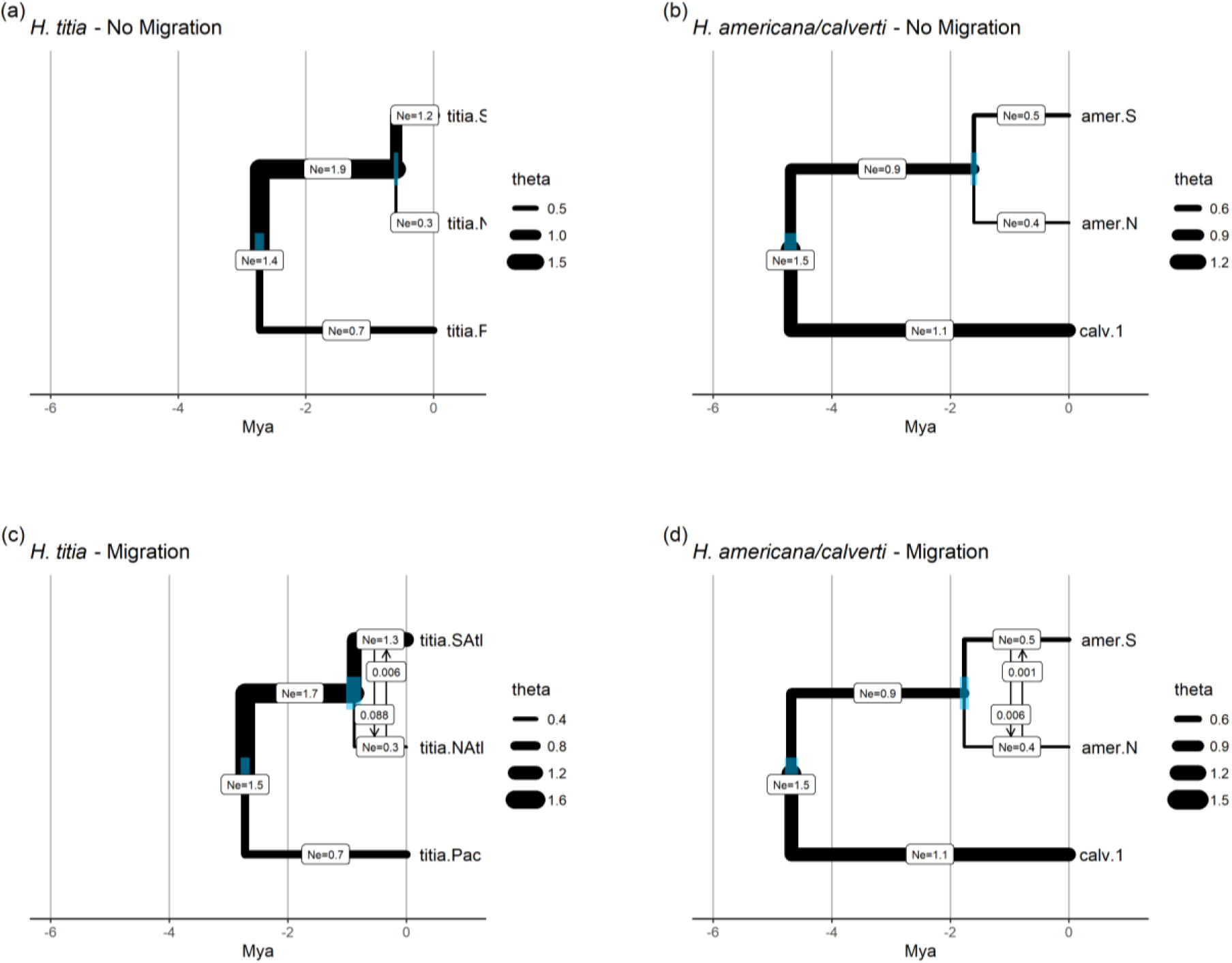
The estimated divergence times (Mya = million years ago) and effective population size (theta - Ne in millions of individuals) from G-PhoCS analysis of *H. titia* and *H. americana.* Migration rate is the number of individuals per generation with vertical arrows indicating direction of migration (from and to). All models ran for 1,000,000 iterations with 10% burn in. Blue bars show 95% highest posterior density for each divergence date (a) Model estimates for *H.titia* with no migration bands. The estimated divergence time for Atlantic and Pacific *H.titia* was 2.72 mya (2.65-2.80 mya HPD) and divergence time between Northern and Southern Atlantic clusters was estimated as 0.59 mya (0.55-0.62 mya HPD). (b) Model estimates for *H. americana* and *H. calverti* with no migration bands. The divergence time for *H. calverti* and *H. americana* was estimated to be 4.7 mya (3.47-4.68 mya HPD). The Northern and Southern *H. americana* clusters diverged 1.60 mya (1.55-1.66 mya). (c) Model estimates for *H. titia* demography with migration bands between Northern and Southern Atlantic *H. titia.* The divergence time between Northern and Southern Atlantic *H. titia* was 0.87 mya (0.76-1.01 mya HPD) and the split between Atlantic and Pacific *H. titia* was 2.71 mya (2.64-2.79 HPD). (d) Model estimates for *H. americana* and *H. calverti* with migration bands between North and Southern *H. americana*. The divergence time for *H. calverti* and *H. americana* was estimated to be 4.68 mya (4.57-4.79 mya HPD). The Northern and Southern *H. americana* clusters diverged 1.75 mya (1.68-1.83 mya). G-PhoCS runs presented here are conducted on the RAD loci mapped de novo using ipyrad..

The estimated effective population sizes for each lineage of *H. titia* were consistent between models and runs. For *H. titia*, the Southern Atlantic *H. titia* lineage had the largest effective population size, around 1.0 million individuals (0.96-1.11 HPD), and the Northern Atlantic lineage had the smallest, around 0.28 million individuals (0.26-0.310 HPD). The Pacific lineage’s effective population size was estimated to be around 0.39 million individuals (0.37-0.42 HPD). *H. calverti* was estimated to have a much greater effective population size than either lineage of *H. americana*; 1.1 million individuals (1.08-1.16 HPD) compared to 0.67 million (0.65-0.70 HPD) for the Southern *H. americana* lineage and 0.41 million (0.39-0.43 HPD) for the Northern *H. americana* lineage..

Where migration was included in the demographic models, the estimated migration rate between populations was low. For all migration bands, the percentage of individuals within each population per generation that were estimated to have originated by migration was between 0.01 and 0.08 individuals per generation). For both *H. titia* and *H. americana*, migration from southern populations to northern populations was estimated to occur more often than migration from northern to southern populations (Figure 5).

### An F1 hybrid at a zone of secondary contact

Calculations of hybrid index and heterozygosity indicated that sample CUAJa02 from site CUAJ01 (an Atlantic drainage near the continental divide) is an F_1_ hybrid between Pacific and Atlantic lineages (Figure 6). Sample CUAJa02 had an autosome heterozygosity of 93.3% and a hybrid index of 0.50, close to the theoretical level of an F_1_ hybrid (100% and 0.5 respectively) and markedly above the heterozygosity of an F_2_ hybrid (50%). A second-generation backcross would produce a hybrid index of 0.25 or 0.75, depending on the proportion of Pacific vs Atlantic parentage. The X chromosome of sample CUAJa02 was nearly entirely homozygous for Pacific alleles. As *Hetaerina* exhibit an XO sex determination system, its parents were likely a female from the Pacific lineage and a male from the Atlantic lineage. The single sex-linked heterozygous site in the hybrid individual had markedly higher read depth than the other SNPs on the X chromosome, suggesting the SNP was autosomal and incorrectly mapped to the X chromosome (Supplementary Figure 24). The rate of heterozygosity across the highly divergent sites was close to zero for all other samples. All other samples either had nearly entirely Pacific or Atlantic genotypes.

**Figure 6.**
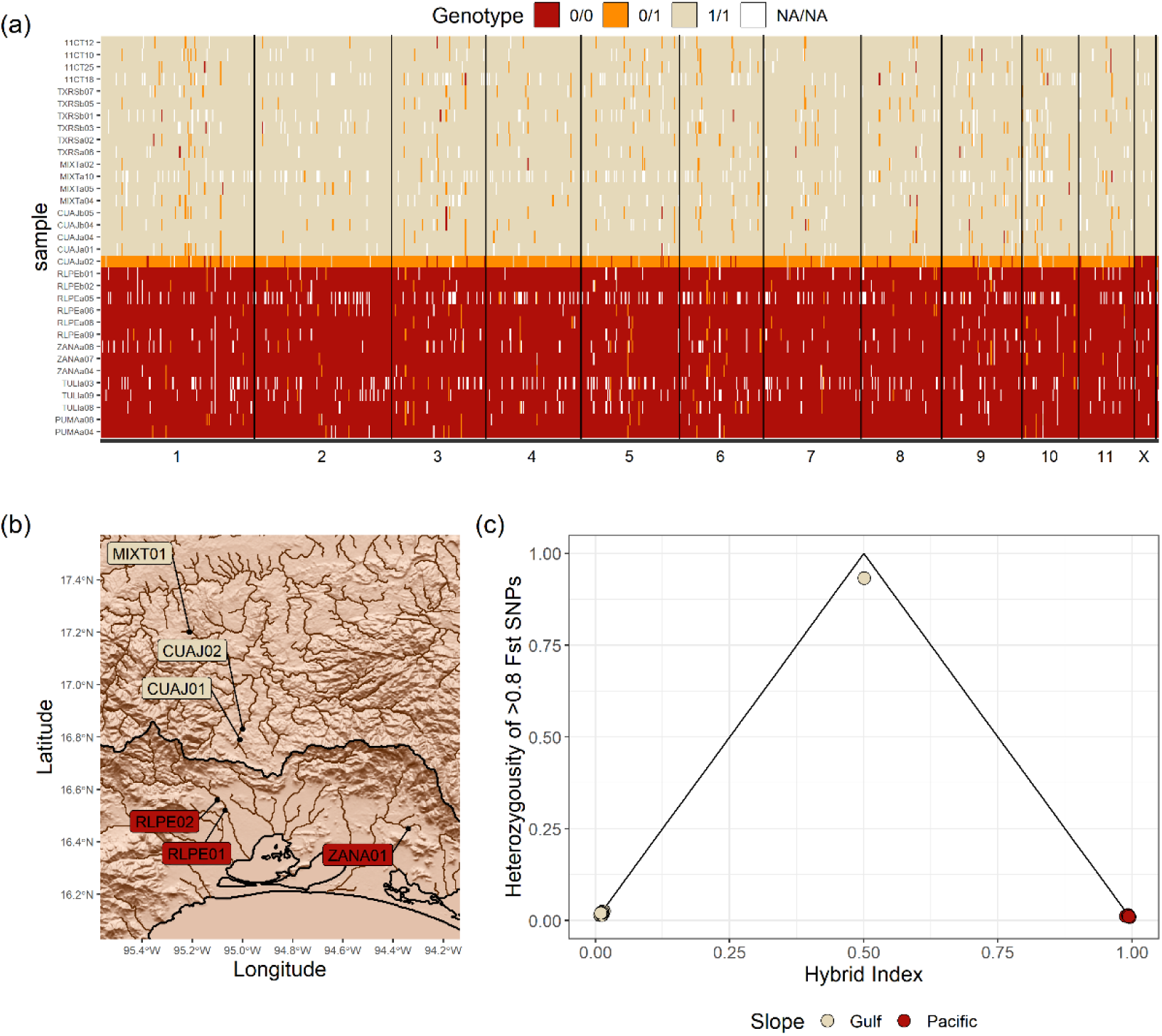
Hybrid zone between Pacific and Southern Atlantic *Hetaerina titia* in the Isthmus of Tehuantepec. (a) Genotypes for 914 autosomal SNPs and 19 sex-linked SNPs that had a greater than 0.8 allele frequency difference between Pacific and Atlantic individuals (calculations excluded samples from CUAJ01/02). Each sample is positioned along the y-axis with each SNP ordered by the position along each chromosome along the x axis. The F_1_ hybrid is sample CUAJa02. Each SNP is coloured by whether they were homozygous for the Pacific allele (0/0 –red), homozygous for the Atlantic allele (1/1 – beige), or heterozygous (0/1 – orange). (b) Sample locations around the Isthmus of Tehuantepec. The Atlantic and Pacific watershed boundary is shown in black (see Figure 1b for a map of wider region and Supplementary Figure 23 for a map of terrain height rather than a shaded relief). (c) A triangle plot showing the hybrid index, measuring the percentage of “parental” genotype, and the heterozygosity of each sample. A theoretical F_1_ hybrid would be placed at the top corner of the triangle. SNPs on the X chromosome were excluded when calculating the hybrid index and heterozygosity.

## Discussion

We reconstructed the spatiotemporal dynamics of divergence in multiple lineages of rubyspot damselflies. As predicted by the non-adaptive radiation model commonly invoked for damselflies (Wellenreuther & Sánchez-Guillén, 2016), we found clear evidence that divergence time between lineages is positively related to levels of reproductive isolation and spatial overlap. Specifically, the older species pair (*H. americana* and *H. calverti*, estimated to have diverged 6.8 mya in our SNAPP analysis) are those whose ranges broadly overlap and exhibit no evidence of introgression; the younger lineages (Pacific *H. titia* and Atlantic *H. titia*, estimated to have diverged 3.7 mya) are found largely in allopatry, with evidence of limited hybridization suggesting strong post-zygotic isolation at a narrow point of secondary contact. In addition, our divergence time estimates reveal deep splits between sister lineages, in agreement with theory (Czekanski-Moir & Rundell, 2019; Rundell & Price, 2009) and other studies that have estimated slow diversification rates in non-adaptive radiations— salamanders (Kozak et al., 2005), killifish (Lambert et al., 2019), blindsnakes (Tiatragul et al., 2023, 2024), and snails (Fehér et al., 2013; Koch et al., 2020). Similarly, models of historical demography estimate extremely little to no migration between rubyspot lineages, consistent with theoretical requirements for non-ecological speciation (Nosil & Flaxman, 2011). Overall, these findings demonstrate that these rubyspot damselfly lineages are at different stages of the non-ecological speciation cycle. These observations also reinforce the converse notion that niche divergence accelerates speciation, highlighting the usefulness of non-ecologically speciating taxa as simplified models for studying the evolution of reproductive isolation.

Although our results support the non-ecological speciation model for damselfly diversification, analysing patterns in additional diverging lineages would help to disentangle the mechanisms leading to reproductive isolation, such as mutation-order processes (Mendelson et al., 2014) and/or reproductive character displacement acting on traits that mediate isolation (e.g., genital morphology and mate recognition; (Pfennig & Pfennig, 2012). Moreover, geographical isolating barriers are not uniform in space and time, which is likely to influence the time it takes for daughter lineages to attain secondary contact and overlap in sympatry.

Our analyses shed light on a number of biogeographic factors that have influenced dynamics in *Hetaerina*. For instance, the three population clusters in *H. titia* identified in our analyses are separated by pronounced barriers to dispersal—the Continental Divide (separating the Pacific and Atlantic clusters) and the Trans-Mexican Volcanic Belt, separating the Northern and Southern Atlantic clusters. These have emerged as important phylogeographic barriers in other studies (Edwards et al., 2022; Mastretta-Yanes et al., 2015). Samples from San Luis Potosí, just north of the Trans-Mexican Volcanic Belt, had a majority Northern *H. titia* ancestry but with a potential small proportion of ancestry from Southern *H. titia.* Further sampling is required in the zone between Northern and Southern Atlantic lineages of *H. titia*, which occurs near a similar divide between Northern and Southern lineages of *H. americana* (Vega-Sánchez et al. 2024). The timing of the split between Pacific and Atlantic lineages of *H. titia* overlaps with the timing of the formation of the Isthmus of Panama; given its phylogenetic affinity with species found in South America (Standring et al., 2022), therefore, one hypothesis is that this split arose from northward dispersal. In other words, the last common ancestor of Pacific and Atlantic lineages of *H. titia* could have occurred in southern Central or northern South America before going locally extinct in that region. Our discovery of an F_1_ hybrid between Pacific and Southern Atlantic *H. titia* on the Isthmus of Tehuantepec demonstrates that the Pacific and Atlantic clusters have come into secondary contact in this region. The site with a hybrid individual is only ∼27 km from the nearest Pacific site where we have found *H. titia*. Here the barrier to dispersal across the Continental Divide is reduced: the elevation of the Isthmus of Tehuantepec is around 200 meters (having dropped from a higher elevation during the Late Miocene and Early Pliocene [Barrier et al., 1998]). In comparison, the mountains east and west of the region extend to over 2000-meters in elevation with limited suitable riparian habitat. Finally, our demographic models estimated the lowest effective population sizes in the population clusters furthest north—a result consistent with demographic declines as a result of glaciation (Hewitt, 2000).

Despite the presence of the hybrid individual, we did not detect any further admixture within these lineages that would suggest a history of introgression, suggesting that post-zygotic isolation may be complete, even if pre-zygotic isolation is not. In combination with the deep divergence time estimated for these lineages, it is likely that *H. titia* sensu lato represents a species complex containing multiple cryptic, reproductively isolated lineages. Further study is required to test whether there is divergence in mating preferences or reproductive traits (e.g., wing colour, genitalia morphology) between the two lineages and/or low hybrid fitness related to reproductive traits. Male mate recognition in rubyspot damselflies is based largely on female wing colour (Drury, Anderson, et al., 2015, 2019; Drury, Okamoto, et al., 2015). Pacific and Atlantic *H. titia* exhibit marked differences in seasonal melanisation, which could allow discrimination between Pacific and Atlantic *H. titia*, but only during the peak-breeding season when newly emerged Atlantic *H. titia* exhibit high levels of wing melanisation (Drury, Anderson, et al., 2015; Drury, Barnes, et al., 2019). Anecdotally, the F_1_ individual male had wing pigmentation intermediate between Pacific and Atlantic lineages (with more red pigment than typical Atlantic individuals and more dark pigment than typical Pacific individuals). We have also attained preliminary whole genome resequencing for an individual with a fully Pacific genotype which was collected from the same Atlantic site and on the same date as the F_1_ hybrid. Characterising mate recognition in Pacific and Atlantic *H. titia* within the site of secondary contact could further our understanding of the evolution of pre-zygotic mating barriers.

We find it unlikely that our sampling coincided with the first contact between Pacific and Atlantic *H. titia* in ∼3.7 million years. What, therefore, has prevented Pacific *H. titia* from becoming more widely sympatric with Atlantic *H. titia*? Sympatry can be prevented by the production of low-fitness hybrids which can cause population decline and local extinction (i.e., sexual exclusion [Irwin & Schluter, 2022; Kuno, 1992; Mikkelsen & Irwin, 2021]). For instance, an increase in hybrid zones, due to climate driven ranges shifts, has been identified as a conservation concern for an endangered species of damselfly (Sánchez-Guillén, Muñoz, et al., 2014). Within *H. titia,* there also may be differences in fitness between Pacific and Atlantic lineages. The high level of melanisation seen in the Atlantic lineages of *H. titia* is beneficial in reducing interspecific behavioural interference (Anderson & Grether, 2011; Drury, Anderson, et al., 2015). Therefore, Atlantic *H. titia* may have an advantage over Pacific *H. titia* within river drainages that contain other species of *Hetaerina,* such as *H. occisa* and *H. americana*, which are found within the river drainage of the hybrid site (personal observation). Interspecific behavioural interference can itself influence the range dynamics of populations (Patterson & Drury, 2023). Consequently, mating and territorial interactions between Pacific and Atlantic *H. titia*, as well as behavioural interference between Pacific *H. titia* and other *Hetaerina* spp. in Atlantic drainages, may be restricting the dispersal of the Pacific *H. titia*.

## Conclusion

We estimated divergence times for multiple lineages in a non-adaptive radiation.

Divergence times correlate well with the stage of the non-ecological speciation cycle of each lineage pair, with the most distantly related lineages found in sympatry and the most closely related being in allopatry. We identified a site where there is contemporary but limited hybridisation between two highly differentiated lineages of the same (currently recognized) species. Collectively, this research provides insight into multiple stages of the non-ecological speciation cycle and paves the way for future work on diversification dynamics in non-adaptive radiations.

## Data Availability

All code is available on GitHub: https://github.com/ChristophePatterson/Phylogeography-Hetaerina and includes SNP libraries in vcf format. Raw demultiplexed sequence reads will be made available on NCBI upon publication.

## Acknowledgements

This work was supported by a NERC Environmental Omics Facility Grant (NEOF1274) to JPD, funding from Durham University (including a Durham Doctoral Studentship to CWP), and NSF DEB-NERC-2040883 to JPD and GFG. Specimen collection was conducted under permit SGPA/ DGVS/04421/21 issued by the Mexican Secretaría de Medio Ambiente y Recursos Naturales (SEMARNAT) to LMC; permit 08112112 issued by the Florida Department of Environmental Protection to JPD; license FD/WL/7/21(04) issued by the Forest Department in Belize to JPD & GFG; permit No. R-SINAC-SE-DT-PI-003-2021 issued by the Costa Rican Ministry of Environment (Minae) and genetic access permission No. 377 issued by the Comisión Institucional de Biodiversidad, Universidad de Costa Rica. CWPs was provided training and support for laboratory work by Gavin Horsburgh at the Natural Environment Research Council Omics Facility (NEOF) at the University of Sheffield. NEOF also provided CWP with training and support with bioinformatics. We thank Andreanna Welch, Lesley Lancaster, Erandi Bonillas-Monge, and Dan Nesbit for helpful feedback on an earlier draft of the manuscript.

