## Supplementary Material for "Spatiotemporal dynamics of non-ecological speciation in rubyspot damselflies (*Hetaerina* spp.)"

**Supporting Information for Spatiotemporal dyanmics of non-ecological speciation in rubyspot damselflies (*Hetaerina* spp.)**

Patterson, C., Brennan, A., Cowling, H., González-Rodríguez, A., Grether, G.F., Mendoza Cuenca, L. Springer, M., Vega-Sanchez, Y. M., and Drury, J.

Supplementary Methods
Supplementary Tables 1-4
Supplementary Figures 1-23

### Supplementary Methods

Library preparation was conducted twice creating two separate ddRAD libraries. Library One was constructed at the NERC environmental omics facility (NEOF), at the University of Sheffield, using the ddRAD protocol from DaCosta and Sorenson (2014). Library Two was conducted at Durham University following the methodology from Franchini *et al.* (2017). Samples extracted for ddRAD library One were initially resuspended using 200µl of buffer AE, and then were desiccated and resuspended in 50µl of buffer AE. Samples extracted for ddRAD library Two were resuspended in 50µl of buffer AE. DNA degradation was visualised via gel electrophoresis and the concentration calculated on a Qubit 2.0 Fluorometer broad range (BR). DNA concentration ranged from 0.47 ng/µl to 26.8 ng/µl with a mean of 4.44 ng/µl. For each sample, this totalled a range 5361ng to 95.4ng of DNA with an average of 888.32ng.

*Library One*

Double digest restriction enzyme association DNA (ddRAD) was created for 190 *Hetaerina* samples (183 *H. titia*, 4 *H. occisa*, and 3 *H. americana* senso lato) using two, six-base restriction enzymes, PstI and EcoRI. Library preparation was conducted using custom Illumina adapters; 92 barcoded P1 adapters and two P2 adapters with 192 unique barcode combinations. P2 adapters contained 8 bases of random nucleotides for the detection of PCR duplication (DaCosta & Sorenson, 2014).

Final DNA concentration for each of the 16 sample pools was calculated by fluorometer and ranged from 84.4ng/µl to 2.99ng/µl, with an average of 30.1ng/µl. 40ng of 16 samples were pooled into separate libraries and then quantified by qPCR. Separate libraries were then pooled into a single library for sequencing and sent for paired end 150 bp sequencing on Illumina NovaSeq 6000.

*Bioinformatics for Library One*

Sequencing was conducted on a NovaSeq 6000 (Illumina) within a single lane. Raw sequence files were demultiplex into one of the two P2 adapters (BC1 and BC2) and processed separately. Initially $212.1 \times 10^6$ BC1 paired read files and $189.1 \times 10^6$ BC2 paired read files were identified. Trimming was conducted using *trimomatic 0.39* to remove low quality bases using a sliding window of 4 and an average base quality of 20. Reads shorter than 15 bases were removed. After filtering $169.9 \times 10^6$ (80.1%) BC1 paired sequences and $153.6 \times 10^6$ (81.2%) BC2 paired sequences remained.

Sequences were processed with STACKS v2.60 (Catchen et al., 2013). clone_filter identified 15.12% of BC1 reads to be PCR clones and were removed, retaining $144.2 \times 10^6$ paired reads. pcr_filter identified 19.66% of BC2 reads to be PCR clones and removed retaining $123.4 \times 10^6$ paired reads.

Reads were demultiplexed using process_radtags using the 8 base unique barcodes on the P1 adapters. 268,996,468 (93.3%) of BC1 reads were retained and assigned to a specific sample, 6.4% of reads did not contain an identifiable barcode and 0.3% could not identify the RAD cut site. 229,259,030 (92.9%) of BC2 reads were retained and assigned to a specific sample, 6.8% did not contain an identifiable barcode and 0.4% could not identify a RAD cut site. All remaining sequence were processed through *cutadapt* to remove both the remaining adapter regions. For all samples the total number of identified paired sequences was 496,388,354. The mean number of reads per sample was 1,292,678 (sd = 824,724) and ranged from 256 to 4,174,260.

*Library Two*

Double digest restriction enzyme association DNA (ddRAD) was created for 192 *Hetaerina* samples (112 *H. titia*, 80 *H. americana* senco lato) again using restriction enzymes PstI and EcoRI. Library preparation was done using custom Illumina adapters; three inner i5 barcoded adapters and four inner i7 adapters plus four outer i5 primers and four outer i7 primers creating 192 unique barcode combinations. Both inner i5 and i7 adapters contained a sequence of 4 bases of random nucleotides for the detection of PCR duplication. This protocol follows Franchini et al. (2017) but the adapters were modified to ligate to PstI and EcoRI restriction sites.

Final size selection was conducted using a pippin prep 2% agarose gel cassette with DF Marker L. 30µl of cleaned pooled PCR product was loaded onto a single lane and size selected for 250-400bp. 40µl of size selected digested, ligated, and indexed DNA was retrieved from the pippin prep elusion module, DNA concentration was quantified using the qubit high sensitivity and size range was quantified using a Tapestation HS d1000 cassette. This yield 40µl of quaddRAD library at 1.08ng/µl (43.2ng in total) which was sent for paired end 150 bp sequencing on Illumina NovaSeq 6000.

*Bioinformatics for Library Two*

Sequencing was conducted on a NovaSeq 6000 (Illumina) within a single lane. Raw sequence files were demultiplex into one of 16 combinations of outer barcodes. Each pool contained on average 16.1x$10^6$ paired reads (range 7.3 x$10^6$ to 16.1 x$10^6$) with 294.1x$10^6$ paired reads in total. Trimming was conducted using *trimomatic 0.39* to remove low quality bases using a sliding window of 4 and an average base quality of 20. Reads shorter than 15 bases were removed. After filtering, 289.5x$10^6$ (98.2%) sequences remained. Sequences were processed with *STACKS v2.60* (Catchen et al., 2013). *clone_filter* identified 17.4% of reads to be PCR clones and were removed, retaining 239.3x$10^6$ paired reads.

Sequences were demultiplexed using *process_radtags* which generated 192 individual samples files. In total 215.5x$10^6$ paired reads (90.1%) were assigned to a specific sample ranging from 671 to 13.3x$10^6$ (mean = 1.12x$10^6$) reads per sample.

*SNP and Loci calling*

For all SNP libraries*, samtools v1.13* was used to align all demultiplex sequence data and stored as a compressed sequence alignment file (.bam). After aligning the sequence reads to the draft genomes, we excluded samples that had a mean depth of less than 5.

Genotype calling was done using *bcftools v1.13* (Danecek et al., 2021; Li, 2011) using the *mpileup* and *call* commands, with a max depth of 10,000 and a prior expected substitution rate of 1e-6. As our samples were collected over a large geographical scale and included multiple species, we did not use Hardy-Weinberg equilibrium calculations for variant calling. After genotyping, variant sites were restricted to SNPs. SNP sites were filtered to remove all SNP calls with a read depth of less than 10, an average read depth across samples greater than 200 per sample, and a quality score less than 20. Further filtering removed SNPs that had a minor allele frequency less than 0.05% (<2 samples), SNPs that were found in less than 80% of samples, and to avoid linkage we randomly selected one SNP from a window of 1000 base pairs using the *prune+* command in *bcftools* (Danecek et al., 2021). See break down in Supplementary Table 1)

*De novo SNP calling using ipyrad*

There are large phylogenetic distances (> 6 million years) between and within the species used in our study (Standring et al., 2022; Vega-Sánchez et al., 2024). Mapping reads to draft genomes with high phylogenetic distance to the populations and species within a study can lead to ascertainment bias in SNP calling (Bohling, 2020). As such we used ipyrad (Eaton & Overcast, 2020) to assembly RAD loci de novo. To reduce computation time, we used a subset of the top three highest coverage samples from each lineage identified using the SNP libraries constructed using draft genomes (18 samples in total across six lineages). The assignment of samples to each lineage did not change across any SNP library. We ran ipyrad following default parameters of the manual. *Denovo* mapping of the sequenced read identified 42,109 RAD loci, with each sample having an average of 14,693 RAD loci sequenced (range 10,319 to 17,554). SNPs were filtered to one random SNP per RAD loci and then removed loci that were missing from more than 20% of samples. For de novo SNP libraries that contained both *H. americana* sensu lato and *H. titia* retained a data set of 518 SNPs. The SNP library that contained only *H. americana* sensu lato retained 2,652 SNPs. The SNP library that contained only *H. titia* retained 1,733.

*Re-sequencing of highly heterozygous samples*

Pre-liminary analysis, using the r package SambaR (de Jong et al., 2021), identified five samples as having high levels of heterozygosity - one sample from Library One and four samples from Library Two. The high heterozygosity samples from Library Two were processed within the same pool and had the same i5 inner barcode and are likely due to a fault in inner barcode ligation. The likely cause of the high heterozygosity for the sample from Library One could not be determined. To determine whether laboratory error was the cause for the high heterozygosity, we re-extracted DNA (from tissue) for the five identified samples and re-sequenced them in line with the methodology described in Library Two. We attained, $4.7x10^6$ paired reads of sequence data which was trimmed to $4.6\ x10^6$ paired reads, removed clones retaining $4.2\ x10^6$ paired reads, and demultiplexed which retained $4.1\ x10^6$ paired reads using the methodology described in Library Two. All four samples from Library Two did not show high levels of heterozygosity after re-sequencing. The sample from Library One retained high levels of heterozygosity.

*Migration analyses in delimitR*

To decrease computation time while running *delimitR* (Smith & Carstens, 2020), we wrote a custom R function called *fastsimcoalsim_sbatch*. Built off the default *delimitR* function *fastsimcoalsim*, *fastsimcoalsim_sbatch* allows each set of fastsimcoal simulations (for each demographic scenario) to be run in parallel using the SLURM HPC management system.

We simulated each scenario using broad, uniform priors. Effective populations size (N*e*) was set from 100,000 to 2,000,000 haploid individuals (Ne/2), divergence time for the first population split (Tdiv1) was 500,000 to 12,000,000 generations, and divergence for the second population split (Tdiv2) was 500,000 to 8,000,000 generations. We set Tdiv1 to be always greater Tdiv2. Migration rates were set from 0.000005 to 0.0005 as a proportion of N*e* per generation.

*Historical demography analyses in G-PhoCS*

In comparison to previous analysis which used SNP datasets, G-PhoCS uses sequences of whole loci. G-PhoCS uses MCMCcoal to estimate divergence times and effective population sizes from multilocus sequence data and allows users to specify migration bands between lineages.

For the SNP mapped to the draft genomes, we used a custom R script to generate consensus sequences for each whole ddRAD loci for each sample using the bcftools consensus command. We identified the position of ddRAD loci using the position of the SNPs identified during the previous calling pipeline. We generated consensus sequences, for each sample, 500 base pairs either side of each identified SNPs. We trimmed loci to remove bases that were outside of the sequenced region then removed loci that had a sequence length of less than 100bp and had greater than 20% missing bases. For the de novo SNP library we generated the RAD loci sequence file using the gphocs output option within ipyrad.

Due to computational constraints, we restricted G-PhoCS analysis to the four samples with highest loci coverage from each ancestral cluster identified by sNMF and include no more than 2000 randomly selected ddRAD loci, similar to Prates *et al.*, (2018). We used the tree topology generated from RAxML, SVDquartet, and SNAPP and ran G-PhoCS with and without migration for all possible migration bands within the tree (with the exception of migration from either of the recently diverged lineages to the older lineage, which is an unresolved issue within G-PhoCS: https://github.com/gphocs-dev/G-PhoCS/issues/25). We ran G-PhoCS with a range of uninformative priors following Prates *et al* (2018). The effective population size (theta - $\theta$) and the divergence times (tau - $\tau$) priors were set using gamma distributions of $\alpha$=1 and $\beta$=20, and $\alpha$=1 and $\beta$=200, respectively. The migration rate prior was set to a gamma distribution with $\alpha$=1 and $\beta$=2e-7. To avoid ascertainment bias in loci mapped to a more closely related draft genome we compared divergence times between lineages of *H. titia* and *H. americana* using loci mapped to the relative heterospecific draft genome and compared that with the de novo loci.

Each run of G-PhoCS was conducted for 1,000,000 iterations with a 10% burn-in and sampled every 100 iterations. We optimised the initial acceptance rate to be 30-40% by running 100 interactions of automatic fine-tuning. Convergence was confirmed by visualising trace plots in R. The R package *coda* was used to calculate the effective sample size (ESS) and the 95% Highest Posterior Density (HPD) from the trace file. We converted mutation rate-scaled parameter estimates of G-PhoCS into the number of diploid individuals and the number of years using 2.8e-9 mutations per base pair per generation (Keightley et al., 2014). We converted generations to years using an estimated generation time of one year.

Supplementary Table 1: The total number of variants identified in calling and the number of variants remaining after the following filtering steps. Removing indels (All single base calls). Removing all base calls that had read depth of less than 10 and greater than 200, and a quality score less than 20 (DP10-200, Q>20). Removing all SNPs not genotyped for 80% of samples, removing SNPs with a minimum allele frequency less than 0.05, and randomly removing all but one SNP within 1000 base window.

| *Species* | Draft genome | Number of samples | All variant calls | All single base calls | DP10-200, Q>20 | 80% Samples | MAF>0.05 | Random 1-1000 |
|---|---|---|---|---|---|---|---|---|
| *all* | *H. americana* | 364 | 134810308 | 134707986 | 2233411 | 80344 | 34506 | 1156 |
| *all* | *H. titia* | 363 | 161441042 | 161305711 | 2871466 | 81701 | 36376 | 1199 |
| *H. americana/ calverti* | *H. americana* | 75 | 73615352 | 73549147 | 4842870 | 1042635 | 151033 | 6997 |
| *H. americana/ calverti* | *H. titia* | 75 | 44189192 | 44164151 | 2258691 | 495629 | 80935 | 4103 |
| *H. titia* | *H. americana* | 286 | 81630947 | 81586704 | 1709783 | 260765 | 73471 | 3674 |
| *H. titia* | *H. titia* | 285 | 136427605 | 136309637 | 2997868 | 494012 | 121745 | 6369 |

Supplementary Table 2: Overview of all analysis presented within the paper and which SNP/loci libraries were used. Each library consists of different combinations of samples from different species and reads were aligned to the draft genome of *Hetaerina americana* (Grether et al., 2023), *Hetaerina titia* (Patterson et al., 2023) or mapped de novo. Each analysis and library are marked with whether the results are presented in the main text (M) or in the supplementary (S). Table includes the number of samples (N) and the total number of SNPs/loci that were included in analysis.

| | | | **Population structure** | | | **Phylogentics** | | | | | | **Demographic/ divergence times** | | | | | | | | | **Introgression** | | |
|---|---|---|---|---|---|---|---|---|---|---|---|---|---|---|---|---|---|---|---|---|---|---|---|
| | | | **sNMF & PCA** | | | **RAxML** | | | **SVDQuartets** | | | **delimitR** | | | **SNAPP** | | | **G-Phocs** | | | **Introgress** | | |
| **Library Number** | **Included species** | **Read alignment** | **SNPs** | **N** | **M/S** | **SNPs** | **N** | **M/S** | **SNPs** | **N** | **M/S** | **2N** | **Down Sample (2N)** | **M/S** | **SNPs** | **N** | **M/S** | **Loci** | **N** | **M/S** | **SNPs (sex-linked)** | **N** | **M/S** |
| #1 | *H. americana* sensu lato & *H. titia* | HetAmer1.0 | - | - | - | 524 | 258 | S | 566 | 258 | S | - | - | - | 571 | 98 | **M** | - | - | - | - | - | - |
| #2 | *H. americana* sensu lato & *H. titia* | HetTit1.0 | - | - | - | 560 | 254 | S | 609 | 254 | S | - | - | - | 603 | 95 | **M** | - | - | - | - | - | - |
| #3 | *H. americana* sensu lato & *H. titia* | *de novo* | - | - | - | - | - | - | - | - | - | - | - | - | 519 | 18 | **M** | - | - | - | - | - | - |
| #4 | *H. americana* sensu lato | HetAmer1.0 | 5259 | 58 | **M** | 3949 | 58 | **M** | 5259 | 58 | S | 22,20,72 | 10,12,18 | S | - | - | - | 2000 | 9 | S | - | - | - |
| #5 | *H. americana* sensu lato | HetTit1.0 | 1816 | 58 | S | 1034 | 58 | S | 1816 | 58 | S | 22,20,72 | 10,11,18 | S | - | - | - | 1814 | 9 | S | - | - | - |
| #6 | *H. americana* sensu lato | *de novo* | - | - | - | - | - | - | - | - | - | - | - | - | - | - | - | 2652 | 9 | **M** | - | - | - |
| #7 | *H. titia* | HetAmer1.0 | 1122 | 207 | S | 846 | 202 | S | 1122 | 202 | S | 100,204,92 | 20,25,20 | S | - | - | - | 1122 | 9 | S | - | - | - |
| #8 | *H. titia* | HetTit1.0 | 3819 | 205 | **M** | 3020 | 199 | **M** | 3819 | 199 | S | 100,204,92 | 26,50,20 | S | - | - | - | 2000 | 9 | S | 914 (19) | 33 | **M** |
| #9 | *H. titia* | *de novo* | - | - | - | - | - | - | - | - | - | - | - | - | - | - | - | 1733 | 9 | **M** | - | - | - |

Supplementary Table 3: Geographical location of all samples for which adequate sequence data was attained. Species refers to the taxonomic classification assigned to sample upon collection, either *H. titia* or *H. americana* sensu lato. Lineage assignment refers to the lineage that the sample was assigned to after genotyping. Either *H. titia* Pacific (titia-Pac), *H. titia* North Atlantic (titia-NAtl), *H. titia* South Atlantic (titia-SAtl), *H. calverti* (calverti), *H. americana* South (americana-S) or *H. americana* North (americana-N).

| Sample name | Sex | Species | Site ID | Lat | Long | Country | Province | Ocean drainage | Lineage assignment |
|---|---|---|---|---|---|---|---|---|---|
| 10OT23 | | titia | 10OT | 18.68 | -96.38 | Mexico | Veracruz | Gulf | titia-SAtl |
| 11CT10 | M | titia | 11CT | 18.37 | -95 | Mexico | Veracruz | Gulf | titia-SAtl |
| 11CT12 | | titia | 11CT | 18.37 | -95 | Mexico | Veracruz | Gulf | titia-SAtl |
| 11CT18 | M | titia | CT | 18.37 | -95 | Mexico | Veracruz | Gulf | titia-SAtl |
| 11CT25 | F | titia | CT | 18.37 | -95 | Mexico | Veracruz | Gulf | titia-SAtl |
| ACSFa01 | M | titia | ACSF01 | 29.83 | -82.65 | United States | Florida | Gulf | titia-NAtl |
| ACSFa02 | M | titia | ACSF01 | 29.83 | -82.65 | United States | Florida | Gulf | titia-NAtl |
| ACSFa03 | M | titia | ACSF01 | 29.83 | -82.65 | United States | Florida | Gulf | titia-NAtl |
| ACSFc01 | M | titia | ACSF03 | 29.83 | -82.68 | United States | Florida | Gulf | titia-NAtl |
| ACSFc02 | M | titia | ACSF03 | 29.83 | -82.68 | United States | Florida | Gulf | titia-NAtl |
| ACSFc03 | M | titia | ACSF03 | 29.83 | -82.68 | United States | Florida | Gulf | titia-NAtl |
| ACTPa02 | M | titia | ACTP01 | 15.29 | -92.67 | Mexico | Chiapas | Pacific | titia-Pac |
| ACTPa03 | M | titia | ACTP01 | 15.29 | -92.67 | Mexico | Chiapas | Pacific | titia-Pac |
| ACTPa08 | M | titia | ACTP01 | 15.29 | -92.67 | Mexico | Chiapas | Pacific | titia-Pac |
| AR0101 | M | titia | AR01 | 33.9 | -92.23 | United States | Arkansas | Gulf | titia-NAtl |
| AR0102 | M | titia | AR01 | 33.9 | -92.23 | United States | Arkansas | Gulf | titia-NAtl |
| AR0103 | M | titia | AR01 | 33.9 | -92.23 | United States | Arkansas | Gulf | titia-NAtl |
| AR0104 | M | titia | AR01 | 33.9 | -92.23 | United States | Arkansas | Gulf | titia-NAtl |
| AR0207 | M | titia | AR02 | 33.7 | -92.03 | United States | Arkansas | Gulf | titia-NAtl |
| AR0208 | M | titia | AR02 | 33.7 | -92.03 | United States | Arkansas | Gulf | titia-NAtl |
| AR0209 | M | titia | AR02 | 33.7 | -92.03 | United States | Arkansas | Gulf | titia-NAtl |
| AR0210 | M | titia | AR02 | 33.7 | -92.03 | United States | Arkansas | Gulf | titia-NAtl |
| ARa02 | M | titia | AR | 18.95 | -103.95 | Mexico | Colima | Pacific | titia-Pac |

| Sample name | Sex | Species | Site ID | Lat | Long | Country | Province | Ocean drainage | Lineage assignment |
|---|---|---|---|---|---|---|---|---|---|
| ARa04 | F | titia | AR | 18.95 | -103.95 | Mexico | Colima | Pacific | titia-Pac |
| ARa05 | M | titia | AR | 18.95 | -103.95 | Mexico | Colima | Pacific | titia-Pac |
| BALB01 | M | titia | BALB | 10.05 | -83.33 | Costa Rica | Limón | Caribbean | titia-SAtl |
| BALB02 | M | titia | BALB | 10.05 | -83.33 | Costa Rica | Limón | Caribbean | titia-SAtl |
| BALB06 | M | titia | BALB | 10.05 | -83.33 | Costa Rica | Limón | Caribbean | titia-SAtl |
| BAMB03 | M | titia | BAMB | 10.06 | -83.37 | Costa Rica | Limón | Caribbean | titia-SAtl |
| BAMB06 | M | titia | BAMB | 10.06 | -83.37 | Costa Rica | Limón | Caribbean | titia-SAtl |
| BZ1509 | M | titia | BZ15 | 17.01 | -88.48 | Belize | | Caribbean | titia-SAtl |
| BZ1513 | M | titia | BZ15 | 17.01 | -88.48 | Belize | | Caribbean | titia-SAtl |
| BZ2105 | M | titia | BZ21 | 17.17 | -89.11 | Belize | | Caribbean | titia-SAtl |
| BZ2106 | M | titia | BZ21 | 17.17 | -89.11 | Belize | | Caribbean | titia-SAtl |
| BZ2303 | M | titia | BZ23 | 17.15 | -88.71 | Belize | | Caribbean | titia-SAtl |
| BZ2601 | M | titia | BZ26 | 17.26 | -88.79 | Belize | | Caribbean | titia-SAtl |
| BZ2604 | M | titia | BZ26 | 17.26 | -88.79 | Belize | | Caribbean | titia-SAtl |
| BZ3004 | M | titia | BZ30 | 17.56 | -88.52 | Belize | | Caribbean | titia-SAtl |
| BZ3005 | M | titia | BZ30 | 17.56 | -88.52 | Belize | | Caribbean | titia-SAtl |
| BZ3204 | M | titia | BZ32 | 17.9 | -88.85 | Belize | | Caribbean | titia-SAtl |
| BZ3205 | M | titia | BZ32 | 17.9 | -88.85 | Belize | | Caribbean | titia-SAtl |
| BZ3209 | F | titia | BZ32 | 17.9 | -88.85 | Belize | | Caribbean | titia-SAtl |
| BZ4001 | M | titia | BZ40 | 17.11 | -88.66 | Belize | | Caribbean | titia-SAtl |
| BZ4002 | M | titia | BZ40 | 17.11 | -88.66 | Belize | | Caribbean | titia-SAtl |
| BZ4602 | M | titia | BZ46 | 16.86 | -89.04 | Belize | | Caribbean | titia-SAtl |
| BZ4603 | M | titia | BZ46 | 16.86 | -89.04 | Belize | | Caribbean | titia-SAtl |
| BZ4801 | M | titia | BZ48 | 16.67 | -88.45 | Belize | | Caribbean | titia-SAtl |
| BZ4802 | M | titia | BZ48 | 16.67 | -88.45 | Belize | | Caribbean | titia-SAtl |
| BZ4905 | M | titia | BZ49 | 16.51 | -88.48 | Belize | | Caribbean | titia-SAtl |
| BZ5001 | M | titia | BZ50 | 16.45 | -88.74 | Belize | | Caribbean | titia-SAtl |

| Sample name | Sex | Species | Site ID | Lat | Long | Country | Province | Ocean drainage | Lineage assignment |
|---|---|---|---|---|---|---|---|---|---|
| BZ5005 | M | titia | BZ50 | 16.45 | -88.74 | Belize | | Caribbean | titia-SAtl |
| BZ5206 | M | titia | BZ52 | 16.15 | -89.01 | Belize | | Caribbean | titia-SAtl |
| CA0103 | M | titia | CA01_CR | 9.76 | -84.62 | Costa Rica | Puntarenas | Pacific | titia-Pac |
| CA0104 | M | titia | CA01_CR | 9.76 | -84.62 | Costa Rica | Puntarenas | Pacific | titia-Pac |
| CUAJa01 | M | titia | CUAJ01 | 16.79 | -95.01 | Mexico | Oaxaca | Gulf | titia-SAtl |
| CUAJa02 | M | titia | CUAJ01 | 16.79 | -95.01 | Mexico | Oaxaca | Gulf | F1 (titia-SAtl and titia-Pac) |
| CUAJa04 | M | titia | CUAJ01 | 16.79 | -95.01 | Mexico | Oaxaca | Gulf | titia-SAtl |
| CUAJb04 | M | titia | CUAJ02 | 16.83 | -95 | Mexico | Oaxaca | Gulf | titia-SAtl |
| CUAJb05 | M | titia | CUAJ02 | 16.83 | -95 | Mexico | Oaxaca | Gulf | titia-SAtl |
| CVa02 | M | titia | CV | 29.33 | -98.87 | United States | Texas | Gulf | titia-NAtl |
| CVa03 | M | titia | CV | 29.33 | -98.87 | United States | Texas | Gulf | titia-NAtl |
| CVa04 | M | titia | CV | 29.33 | -98.87 | United States | Texas | Gulf | titia-NAtl |
| CVa08 | M | titia | CV | 29.33 | -98.87 | United States | Texas | Gulf | titia-NAtl |
| CVa13 | M | titia | CV | 29.33 | -98.87 | United States | Texas | Gulf | titia-NAtl |
| CYa04 | M | titia | CY | 21.75 | -98.96 | Mexico | San Luis Potosí | Gulf | titia-NAtl |
| CYa07 | F | titia | CY | 21.75 | -98.96 | Mexico | San Luis Potosí | Gulf | titia-NAtl |
| CYa10 | F | titia | CY | 21.75 | -98.96 | Mexico | San Luis Potosí | Gulf | titia-NAtl |
| CYa11 | F | titia | CY | 21.75 | -98.96 | Mexico | San Luis Potosí | Gulf | titia-NAtl |
| ESRBa10 | M | titia | ESRB1 | 9.72 | -82.97 | Costa Rica | Limón | Caribbean | titia-SAtl |
| GA0203 | M | titia | GA02 | 8.7 | -83.19 | Costa Rica | Puntarenas | Pacific | titia-Pac |
| GCJRa01 | M | titia | GCJR01 | 37.67 | -77.89 | United States | Virginia | Atlantic | titia-NAtl |
| GCJRa02 | M | titia | GCJR01 | 37.67 | -77.89 | United States | Virginia | Atlantic | titia-NAtl |
| GCJRa05 | M | titia | GCJR01 | 37.67 | -77.89 | United States | Virginia | Atlantic | titia-NAtl |

| Sample name | Sex | Species | Site ID | Lat | Long | Country | Province | Ocean drainage | Lineage assignment |
|---|---|---|---|---|---|---|---|---|---|
| GOa02 | M | titia | GO01 | 8.64 | -83.2 | Costa Rica | Puntarenas | Pacific | titia-Pac |
| GOa07 | M | titia | GO01 | 8.64 | -83.2 | Costa Rica | Puntarenas | Pacific | titia-Pac |
| HCARa02 | M | titia | HCAR01 | 27.88 | -82.17 | United States | Florida | Gulf | titia-NAtl |
| HCARa03 | F | titia | HCAR01 | 27.88 | -82.17 | United States | Florida | Gulf | titia-NAtl |
| HCARa05 | M | titia | HCAR01 | 27.88 | -82.17 | United States | Florida | Gulf | titia-NAtl |
| HCARb01 | M | titia | HCAR02 | 27.87 | -82.14 | United States | Florida | Gulf | titia-NAtl |
| HCARb02 | F | titia | HCAR02 | 27.87 | -82.14 | United States | Florida | Gulf | titia-NAtl |
| HCICa01 | M | titia | HCIC01 | 28.09 | -82.09 | United States | Florida | Gulf | titia-NAtl |
| HCICa02 | M | titia | HCIC01 | 28.09 | -82.09 | United States | Florida | Gulf | titia-NAtl |
| HCICa03 | F | titia | HCIC01 | 28.09 | -82.09 | United States | Florida | Gulf | titia-NAtl |
| HtiCn13 | M | titia | HtiCn | 18.25 | -103.23 | Mexico | Michoacán | Pacific | titia-Pac |
| HtiCn2 | M | titia | HtiCn | 18.25 | -103.23 | Mexico | Michoacán | Pacific | titia-Pac |
| HtiCn7 | M | titia | HtiCn | 18.25 | -103.23 | Mexico | Michoacán | Pacific | titia-Pac |
| HtiNR1 | M | titia | HtiNR | 15.9 | -91.99 | Mexico | Chiapas | Gulf | titia-SAtl |
| HtiPo10 | M | titia | HtiPo | 18.04 | -103.54 | Mexico | Michoacán | Pacific | titia-Pac |
| HtiPo2 | M | titia | HtiPo | 18.04 | -103.54 | Mexico | Michoacán | Pacific | titia-Pac |
| HtiPo6 | M | titia | HtiPo | 18.04 | -103.54 | Mexico | Michoacán | Pacific | titia-Pac |
| HtiTi14 | M | titia | HtiTi | 18.67 | -103.69 | Mexico | Michoacán | Pacific | titia-Pac |
| HtiTi15 | M | titia | HtiTi | 18.67 | -103.69 | Mexico | Michoacán | Pacific | titia-Pac |
| HXRCb05 | F | titia | HXRC02 | 15.75 | -96.3 | Mexico | Oaxaca | Pacific | titia-Pac |
| HXRCb06 | F | titia | HXRC02 | 15.75 | -96.3 | Mexico | Oaxaca | Pacific | titia-Pac |
| HXRCb08 | M | titia | HXRC02 | 15.75 | -96.3 | Mexico | Oaxaca | Pacific | titia-Pac |
| IA0406 | M | titia | IA04 | 41.3 | -94.07 | United States | Iowa | Gulf | titia-NAtl |
| IA0408 | M | titia | IA04 | 41.3 | -94.07 | United States | Iowa | Gulf | titia-NAtl |
| IA0413 | M | titia | IA04 | 41.3 | -94.07 | United States | Iowa | Gulf | titia-NAtl |
| IA0416 | F | titia | IA04 | 41.3 | -94.07 | United States | Iowa | Gulf | titia-NAtl |
| IA0417 | M | titia | IA04 | 41.3 | -94.07 | United States | Iowa | Gulf | titia-NAtl |

| Sample name | Sex | Species | Site ID | Lat | Long | Country | Province | Ocean drainage | Lineage assignment |
|---|---|---|---|---|---|---|---|---|---|
| IA0418 | M | titia | IA04 | 41.3 | -94.07 | United States | Iowa | Gulf | titia-NAtl |
| IL0121 | M | titia | IL01 | 42.32 | -89.36 | United States | Illinois | Gulf | titia-NAtl |
| IL0123 | M | titia | IL01 | 42.32 | -89.36 | United States | Illinois | Gulf | titia-NAtl |
| IL0135 | M | titia | IL01 | 42.32 | -89.36 | United States | Illinois | Gulf | titia-NAtl |
| IL0140 | F | titia | IL01 | 42.32 | -89.36 | United States | Illinois | Gulf | titia-NAtl |
| Il0408 | M | titia | IL04 | 42.02 | -89.33 | United States | Illinois | Gulf | titia-NAtl |
| Il0409 | M | titia | IL04 | 42.02 | -89.33 | United States | Illinois | Gulf | titia-NAtl |
| IL0413 | M | titia | IL04 | 42.02 | -89.33 | United States | Illinois | Gulf | titia-NAtl |
| LCPCa02 | M | titia | LCPC1 | 31.14 | -89.64 | United States | Mississippi | Gulf | titia-NAtl |
| LCPCa03 | M | titia | LCPC1 | 31.14 | -89.64 | United States | Mississippi | Gulf | titia-NAtl |
| LCSBc01 | M | titia | LCSB3 | 31.16 | -89.61 | United States | Mississippi | Gulf | titia-NAtl |
| MACM02 | F | titia | MACM | 10.1 | -83.25 | Costa Rica | Limón | Caribbean | titia-SAtl |
| MC02.001 | M | titia | MC02 | 30.49 | -94.83 | United States | Texas | Gulf | titia-NAtl |
| MC02.002 | M | titia | MC02 | 30.49 | -94.83 | United States | Texas | Gulf | titia-NAtl |
| MC02.003 | M | titia | MC02 | 30.49 | -94.83 | United States | Texas | Gulf | titia-NAtl |
| MEBA013928 | F | titia | MEBA01 | 29.72 | -99.08 | United States | Texas | Gulf | titia-NAtl |
| MI0111 | M | titia | MI01 | 42.23 | -84.49 | United States | Michigan | Atlantic | titia-NAtl |
| MI0112 | M | titia | MI01 | 42.23 | -84.49 | United States | Michigan | Atlantic | titia-NAtl |
| MI0113 | M | titia | MI01 | 42.23 | -84.49 | United States | Michigan | Atlantic | titia-NAtl |
| MI0114 | F | titia | MI02 | 42.23 | -84.49 | United States | Michigan | Atlantic | titia-NAtl |
| MI0162 | F | titia | MI01 | 42.23 | -84.49 | United States | Michigan | Atlantic | titia-NAtl |
| MI0225 | F | titia | MI02 | 42.5 | -83.74 | United States | Michigan | Atlantic | titia-NAtl |
| MI0230 | F | titia | MI02 | 42.5 | -83.74 | United States | Michigan | Atlantic | titia-NAtl |
| MI0238 | M | titia | MI02 | 42.5 | -83.74 | United States | Michigan | Atlantic | titia-NAtl |
| MI0239 | M | titia | MI02 | 42.5 | -83.74 | United States | Michigan | Atlantic | titia-NAtl |
| MIXTa02 | M | titia | MIXT01 | 17.2 | -95.21 | Mexico | Oaxaca | Gulf | titia-SAtl |
| MIXTa04 | M | titia | MIXT01 | 17.2 | -95.21 | Mexico | Oaxaca | Gulf | titia-SAtl |

| Sample name | Sex | Species | Site ID | Lat | Long | Country | Province | Ocean drainage | Lineage assignment |
|---|---|---|---|---|---|---|---|---|---|
| MIXTa05 | M | titia | MIXT01 | 17.2 | -95.21 | Mexico | Oaxaca | Gulf | titia-SAtl |
| MIXTa10 | M | titia | MIXT01 | 17.2 | -95.21 | Mexico | Oaxaca | Gulf | titia-SAtl |
| MO0210 | M | titia | MO02 | 36.95 | -90.99 | United States | Missouri | Gulf | titia-NAtl |
| MO0211 | M | titia | MO02 | 36.95 | -90.99 | United States | Missouri | Gulf | titia-NAtl |
| MO0309 | M | titia | MO03 | 36.56 | -90.35 | United States | Missouri | Gulf | titia-NAtl |
| MO0311 | M | titia | MO03 | 36.56 | -90.35 | United States | Missouri | Gulf | titia-NAtl |
| MO0313 | M | titia | MO03 | 36.56 | -90.35 | United States | Missouri | Gulf | titia-NAtl |
| MO0351 | F | titia | MO03 | 36.56 | -90.35 | United States | Missouri | Gulf | titia-NAtl |
| MS0108 | M | titia | MS01 | 32.52 | -90.74 | United States | Mississippi | Gulf | titia-NAtl |
| MS0114 | M | titia | MS01 | 32.52 | -90.74 | United States | Mississippi | Gulf | titia-NAtl |
| MS0116 | M | titia | MS01 | 32.52 | -90.74 | United States | Mississippi | Gulf | titia-NAtl |
| NA0103 | M | titia | NA01 | 9.74 | -84.63 | Costa Rica | | Pacific | titia-Pac |
| NCRRb04 | M | titia | NCRR02 | 37.84 | -78.81 | United States | Virginia | Atlantic | titia-NAtl |
| NCRRb05 | M | titia | NCRR02 | 37.84 | -78.81 | United States | Virginia | Atlantic | titia-NAtl |
| NMSCa01 | M | titia | NMSC01 | 15.77 | -93.3 | Mexico | Chiapas | Pacific | titia-Pac |
| NMSCa02 | M | titia | NMSC01 | 15.77 | -93.3 | Mexico | Chiapas | Pacific | titia-Pac |
| NMSCa05 | M | titia | NMSC01 | 15.77 | -93.3 | Mexico | Chiapas | Pacific | titia-Pac |
| NPKBa05 | F | titia | NPKB1 | 31.52 | -93.14 | United States | Louisiana | Gulf | titia-NAtl |
| NPKBa06 | F | titia | NPKB1 | 31.52 | -93.14 | United States | Louisiana | Gulf | titia-NAtl |
| NPKBa10 | M | titia | NPKB1 | 31.52 | -93.14 | United States | Louisiana | Gulf | titia-NAtl |
| NR01.01 | M | titia | NR01 | 30.36 | -94.09 | United States | Texas | Gulf | titia-NAtl |
| NR02.06 | M | titia | NR02 | 30.68 | -94.09 | United States | Texas | Gulf | titia-NAtl |
| OK0101 | M | titia | OK01 | 35.24 | -98.55 | United States | Oklahoma | Gulf | titia-NAtl |
| PAa01 | M | titia | PA | 18.55 | -95.07 | Mexico | Veracruz | Gulf | titia-SAtl |
| PAa02 | M | titia | PA | 18.55 | -95.07 | Mexico | Veracruz | Gulf | titia-SAtl |
| PAa07 | F | titia | PA | 18.55 | -95.07 | Mexico | Veracruz | Gulf | titia-SAtl |
| PC01.024 | M | titia | PC01 | 31.08 | -93.49 | United States | Louisiana | Gulf | titia-NAtl |

| **Sample name** | **Sex** | **Species** | **Site ID** | **Lat** | **Long** | **Country** | **Province** | **Ocean drainage** | **Lineage assignment** |
|---|---|---|---|---|---|---|---|---|---|
| PCa01 | M | titia | PC01 | 31.08 | -93.49 | United States | Louisiana | Gulf | titia-NAtl |
| PCCC1 | M | titia | PCCC1 | 30.99 | -89.01 | United States | Mississippi | Gulf | titia-NAtl |
| PSa03 | M | titia | ps01 | 9.54 | -84.28 | Costa Rica | Puntarenas | Pacific | titia-Pac |
| PSa04 | M | titia | ps01 | 9.54 | -84.28 | Costa Rica | Puntarenas | Pacific | titia-Pac |
| PSa06 | B | titia | ps01 | 9.54 | -84.28 | Costa Rica | Puntarenas | Pacific | titia-Pac |
| PUMAa04 | F | titia | PUMA01 | 15.77 | -93.33 | Mexico | Chiapas | Pacific | titia-Pac |
| PUMAa08 | M | titia | PUMA01 | 15.77 | -93.33 | Mexico | Chiapas | Pacific | titia-Pac |
| RCJRa02 | M | titia | RCJR01 | 37.52 | -77.47 | United States | Virginia | Atlantic | titia-NAtl |
| RCJRa04 | M | titia | RCJR01 | 37.52 | -77.47 | United States | Virginia | Atlantic | titia-NAtl |
| RCJRa05 | M | titia | RCJR01 | 37.52 | -77.47 | United States | Virginia | Atlantic | titia-NAtl |
| RCJRb02 | M | titia | RCJR02 | 37.55 | -77.52 | United States | Virginia | Atlantic | titia-NAtl |
| RCJRb04 | M | titia | RCJR02 | 37.55 | -77.52 | United States | Virginia | Atlantic | titia-NAtl |
| RLPEa05 | M | titia | RLPE01 | 16.52 | -95.07 | Mexico | Oaxaca | Pacific | titia-Pac |
| RLPEa06 | M | titia | RLPE01 | 16.52 | -95.07 | Mexico | Oaxaca | Pacific | titia-Pac |
| RLPEa08 | M | titia | RLPE01 | 16.52 | -95.07 | Mexico | Oaxaca | Pacific | titia-Pac |
| RLPEa09 | M | titia | RLPE01 | 16.52 | -95.07 | Mexico | Oaxaca | Pacific | titia-Pac |
| RLPEb02 | M | titia | RLPE02 | 16.56 | -95.1 | Mexico | Oaxaca | Pacific | titia-Pac |
| SCSCa01 | F | titia | SCSC01 | 28.72 | -81.31 | United States | Florida | Atlantic | titia-NAtl |
| SCSCa02 | F | titia | SCSC01 | 28.72 | -81.31 | United States | Florida | Atlantic | titia-NAtl |
| SCSCa03 | F | titia | SCSC01 | 28.72 | -81.31 | United States | Florida | Atlantic | titia-NAtl |
| SJ01.21 | M | titia | SJ01_US | 30.21 | -95.4 | United States | Texas | Gulf | titia-NAtl |
| SMAA06 | M | titia | SMAA2 | 10.04 | -83.33 | Costa Rica | Limón | Caribbean | titia-SAtl |
| SMAA07 | M M | titia | SMAA2 | 10.04 | -83.33 | Costa Rica | Limón | Caribbean | titia-SAtl |
| ST0102 | M | titia | ST01 | 9.38 | -84.02 | Costa Rica | Puntarenas | Pacific | titia-Pac |
| ST0103 | M | titia | ST01 | 9.38 | -84.02 | Costa Rica | Puntarenas | Pacific | titia-Pac |
| STDMa04 | M | titia | STDM01 | 15.82 | -96.69 | Mexico | Oaxaca | Pacific | titia-Pac |
| STDMa06 | M | titia | STDM01 | 15.82 | -96.69 | Mexico | Oaxaca | Pacific | titia-Pac |

| Sample name | Sex | Species | Site ID | Lat | Long | Country | Province | Ocean drainage | Lineage assignment |
|---|---|---|---|---|---|---|---|---|---|
| TN0101 | M | titia | TN01 | 35.03 | -89.35 | United States | Tennessee | Gulf | titia-NAtl |
| TN0103 | M | titia | TN01 | 35.03 | -89.35 | United States | Tennessee | Gulf | titia-NAtl |
| TN0105 | M | titia | TN01 | 35.03 | -89.35 | United States | Tennessee | Gulf | titia-NAtl |
| TN0411 | M | titia | TN04 | 36.55 | -87.14 | United States | Tennessee | Gulf | titia-NAtl |
| TN0423 | M | titia | TN04 | 36.55 | -87.14 | United States | Tennessee | Gulf | titia-NAtl |
| TN0425 | M | titia | TN04 | 36.55 | -87.14 | United States | Tennessee | Gulf | titia-NAtl |
| TN0427 | M | titia | TN04 | 36.55 | -87.14 | United States | Tennessee | Gulf | titia-NAtl |
| TULIa03 | M | titia | TULI01 | 15.89 | -93.5 | Mexico | Chiapas | Pacific | titia-Pac |
| TULIa08 | F | titia | TULI01 | 15.89 | -93.5 | Mexico | Chiapas | Pacific | titia-Pac |
| TULIa09 | F | titia | TULI01 | 15.89 | -93.5 | Mexico | Chiapas | Pacific | titia-Pac |
| TXRSa02 | M | titia | TXRS01 | 16.65 | -93.16 | Mexico | Chiapas | Gulf | titia-SAtl |
| TXRSa06 | M | titia | TXRS01 | 16.65 | -93.16 | Mexico | Chiapas | Gulf | titia-SAtl |
| TXRSb01 | M | titia | TXRS02 | 16.65 | -93.16 | Mexico | Chiapas | Gulf | titia-SAtl |
| TXRSb03 | M | titia | TXRS02 | 16.65 | -93.16 | Mexico | Chiapas | Gulf | titia-SAtl |
| TXRSb05 | M | titia | TXRS02 | 16.65 | -93.16 | Mexico | Chiapas | Gulf | titia-SAtl |
| TXRSb07 | M | titia | TXRS02 | 16.65 | -93.16 | Mexico | Chiapas | Gulf | titia-SAtl |
| WCKCa01 | M | titia | WCKC1 | 31.05 | -90.17 | United States | Mississippi | Gulf | titia-NAtl |
| WCKCa02 | M | titia | WCKC1 | 31.05 | -90.17 | United States | Mississippi | Gulf | titia-NAtl |
| WCKCa03 | M | titia | WCKC1 | 31.05 | -90.17 | United States | Mississippi | Gulf | titia-NAtl |
| ZANAa04 | M | titia | ZANA01 | 16.45 | -94.34 | Mexico | Oaxaca | Pacific | titia-Pac |
| ZANAa06 | M | titia | ZANA01 | 16.45 | -94.34 | Mexico | Oaxaca | Pacific | titia-Pac |
| 05LT11 | M | americana | 05lt | 20.73 | -103.62 | Mexico | Jalisco | Pacific | americana-S |
| 05PU01 | M | americana | 05pu | 19.5 | -104.67 | Mexico | Jalisco | Pacific | calverti |
| 05SI11 | F | americana | 05si | 19.56 | -104.79 | Mexico | Jalisco | Pacific | americana-S |
| 05SI24 | F | americana | 05si | 19.56 | -104.79 | Mexico | Jalisco | Pacific | calverti |
| 10AP36 | f | americana | AP | 19.47 | -96.48 | Mexico | Veracruz | | calverti |
| EAFB014142 | M | americana | EAFB01 | 30.54 | -86.86 | United States | Florida | Gulf | americana-N |

| Sample name | Sex | Species | Site ID | Lat | Long | Country | Province | Ocean drainage | Lineage assignment |
|---|---|---|---|---|---|---|---|---|---|
| EAFB014144 | M | americana | EAFB01 | 30.54 | -86.86 | United States | Florida | Gulf | americana-N |
| EAFB014146 | M | americana | EAFB01 | 30.54 | -86.86 | United States | Florida | Gulf | americana-N |
| EAFB014149 | F | americana | EAFB01 | 30.54 | -86.86 | United States | Florida | Gulf | americana-N |
| EC0101 | m | americana | EC01 | 23.1403 | -99.1158 | Mexico | Tamaulipas | | americana-S |
| GL23892 | M | americana | GL2 | 30.07 | -99.24 | United States | Texas | Gulf | americana-N |
| GL23898 | F | americana | GL2 | 30.07 | -99.24 | United States | Texas | Gulf | americana-N |
| HXRCaAM01 | M | americana | HXRC01 | 15.84 | -96.33 | Mexico | Oaxaca | Pacific | calverti |
| HXRCcAM01 | M | americana | HXRC03 | 15.86 | -96.31 | Mexico | Oaxaca | Pacific | americana-S |
| HXRCcAM02 | F | americana | HXRC03 | 15.86 | -96.31 | Mexico | Oaxaca | Pacific | americana-S |
| HXRCcAM03 | F | americana | HXRC03 | 15.86 | -96.31 | Mexico | Oaxaca | Pacific | americana-S |
| IA0101 | M | americana | IA01 | 41.29 | -94.15 | United States | Iowa | Gulf | americana-N |
| IA0103 | F | americana | IA01 | 41.29 | -94.15 | United States | Iowa | Gulf | americana-N |
| IA0506 | M | americana | IA05 | 42.3 | -92.02 | United States | Iowa | Gulf | americana-N |
| IA0607 | F | americana | IA05 | 42.3 | -92.02 | United States | Iowa | Gulf | americana-N |
| IA0608 | M | americana | IA05 | 42.3 | -92.02 | United States | Iowa | Gulf | americana-N |
| IL0203 | F | americana | IL02 | 41.64 | -88.07 | United States | Illinois | Gulf | americana-N |
| IL0204 | M | americana | IL02 | 41.64 | -88.07 | United States | Illinois | Gulf | americana-N |
| IL0301 | F | americana | IL03 | 42.46 | -89.24 | United States | Illinois | Gulf | americana-N |
| IL0302 | M | americana | IL03 | 42.46 | -89.24 | United States | Illinois | Gulf | americana-N |
| MEBA014006 | M | americana | MEBA01 | 29.72 | -99.08 | United States | Texas | Gulf | americana-N |
| MI0407 | M | americana | MI04 | 42.38 | -83.92 | United States | Michigan | Atlantic | americana-N |
| MI0409 | F | americana | MI04 | 42.38 | -83.92 | United States | Michigan | Atlantic | americana-N |
| MI0504 | M | americana | MI05 | 43.46 | -84.08 | United States | Michigan | Atlantic | americana-N |

| Sample name | Sex | Species | Site ID | Lat | Long | Country | Province | Ocean drainage | Lineage assignment |
|---|---|---|---|---|---|---|---|---|---|
| MI0505 | M | americana | MI05 | 43.46 | -84.08 | United States | Michigan | Atlantic | americana-N |
| MO0110 | M | americana | MO01 | 37.44 | -90.96 | United States | Missouri | Gulf | americana-N |
| MO0111 | F | americana | MO01 | 37.44 | -90.96 | United States | Missouri | Gulf | americana-N |
| MO0112 | M | americana | MO01 | 37.44 | -90.96 | United States | Missouri | Gulf | americana-N |
| MO0213 | F | americana | MO02 | 36.95 | -90.99 | United States | Missouri | Gulf | americana-N |
| MO0214 | M | americana | MO02 | 36.95 | -90.99 | United States | Missouri | Gulf | americana-N |
| MS0222 | M | americana | MS01 | 32.52 | -90.74 | United States | Mississippi | Gulf | americana-N |
| NMSCaAM03 | F | americana | NMSC01 | 15.77 | -93.3 | Mexico | Chiapas | Pacific | calverti |
| OK0102 | F | americana | OK01 | 35.24 | -98.55 | United States | Oklahoma | Gulf | americana-N |
| OK0104 | F | americana | OK01 | 35.24 | -98.55 | United States | Oklahoma | Gulf | americana-N |
| OK0105 | M | americana | OK01 | 35.24 | -98.55 | United States | Oklahoma | Gulf | americana-N |
| RG0101 | m | americana | RG01 | 18.41 | -95.17 | Mexico | Veracruz | | calverti |
| RG0103 | f | americana | RG01 | 18.41 | -95.17 | Mexico | Veracruz | | calverti |
| SJ01.20 | M | americana | SJ01_US | 30.21 | -95.4 | United States | Texas | Gulf | americana-N |
| STDMaAM01 | M | americana | STDM01 | 15.82 | -96.69 | Mexico | Oaxaca | Pacific | americana-S |
| STDMaAM02 | M | americana | STDM01 | 15.82 | -96.69 | Mexico | Oaxaca | Pacific | americana-S |
| TN0119 | M | americana | TN01 | 35.03 | -89.35 | United States | Tennessee | Gulf | americana-N |
| TN0120 | M | americana | TN01 | 35.03 | -89.35 | United States | Tennessee | Gulf | americana-N |
| TN0413 | F | americana | TN04 | 36.55 | -87.14 | United States | Tennessee | Gulf | americana-N |
| TN0414 | M | americana | TN04 | 36.55 | -87.14 | United States | Tennessee | Gulf | americana-N |
| TULIaAM01 | M | americana | TULI01 | 15.89 | -93.5 | Mexico | Chiapas | Pacific | calverti |
| TULIaAM02 | F | americana | TULI01 | 15.89 | -93.5 | Mexico | Chiapas | Pacific | calverti |
| TXRSaAM01 | F | americana | TXRS01 | 16.65 | -93.16 | Mexico | Chiapas | Gulf | americana-S |
| TXRSaAM04 | M | americana | TXRS01 | 16.65 | -93.16 | Mexico | Chiapas | Gulf | americana-S |
| TXRSbAM02 | M | americana | TXRS02 | 16.65 | -93.16 | Mexico | Chiapas | Gulf | americana-S |

| Sample name | Sex | Species | Site ID | Lat | Long | Country | Province | Ocean drainage | Lineage assignment |
|---|---|---|---|---|---|---|---|---|---|
| TXRSbAM03 | M | americana | TXRS02 | 16.65 | -93.16 | Mexico | Chiapas | Gulf | americana-S |
| TXRSbAM04 | F | americana | TXRS02 | 16.65 | -93.16 | Mexico | Chiapas | Gulf | calverti |
| WCKC14173 | M | americana | WCKC1 | 31.05 | -90.17 | United States | Mississippi | Gulf | americana-N |
| WCKC14176 | M | americana | WCKC1 | 31.05 | -90.17 | United States | Mississippi | Gulf | americana-N |

Supplementary Table 4: The number of votes received for each demographic scenario from the random forest classifier when applied to the observed data. For schematic diagrams of each demographic scenario see Supplementary Figure 2.

| **Observed data** | | **Model votes** | | | | | | | | | | | | | | |
|---|---|---|---|---|---|---|---|---|---|---|---|---|---|---|---|---|
| **Species** | **Draft genome** | **1** | **2** | **3** | **4** | **5** | **6** | **7** | **8** | **9** | **10** | **11** | **12** | **13** | **Selected model** | **Post probability** |
| *H. titia* | *H. titia* | 0 | 0 | 0 | 0 | 143 | 0 | 0 | 0 | 0 | 0 | 0 | 0 | 857 | 13 | 0.83 |
| *H. titia* | *H. americana* | 0 | 0 | 0 | 0 | 438 | 1 | 2 | 0 | 0 | 0 | 0 | 0 | 559 | 13 | 0.58 |
| *H. americana / calverti* | *H. titia* | 0 | 0 | 0 | 0 | 556 | 0 | 124 | 12 | 0 | 1 | 1 | 0 | 441 | 5 | 0.74 |
| *H. americana / calverti* | *H. americana* | 0 | 0 | 0 | 0 | 225 | 0 | 0 | 0 | 0 | 0 | 0 | 0 | 775 | 13 | 0.87 |

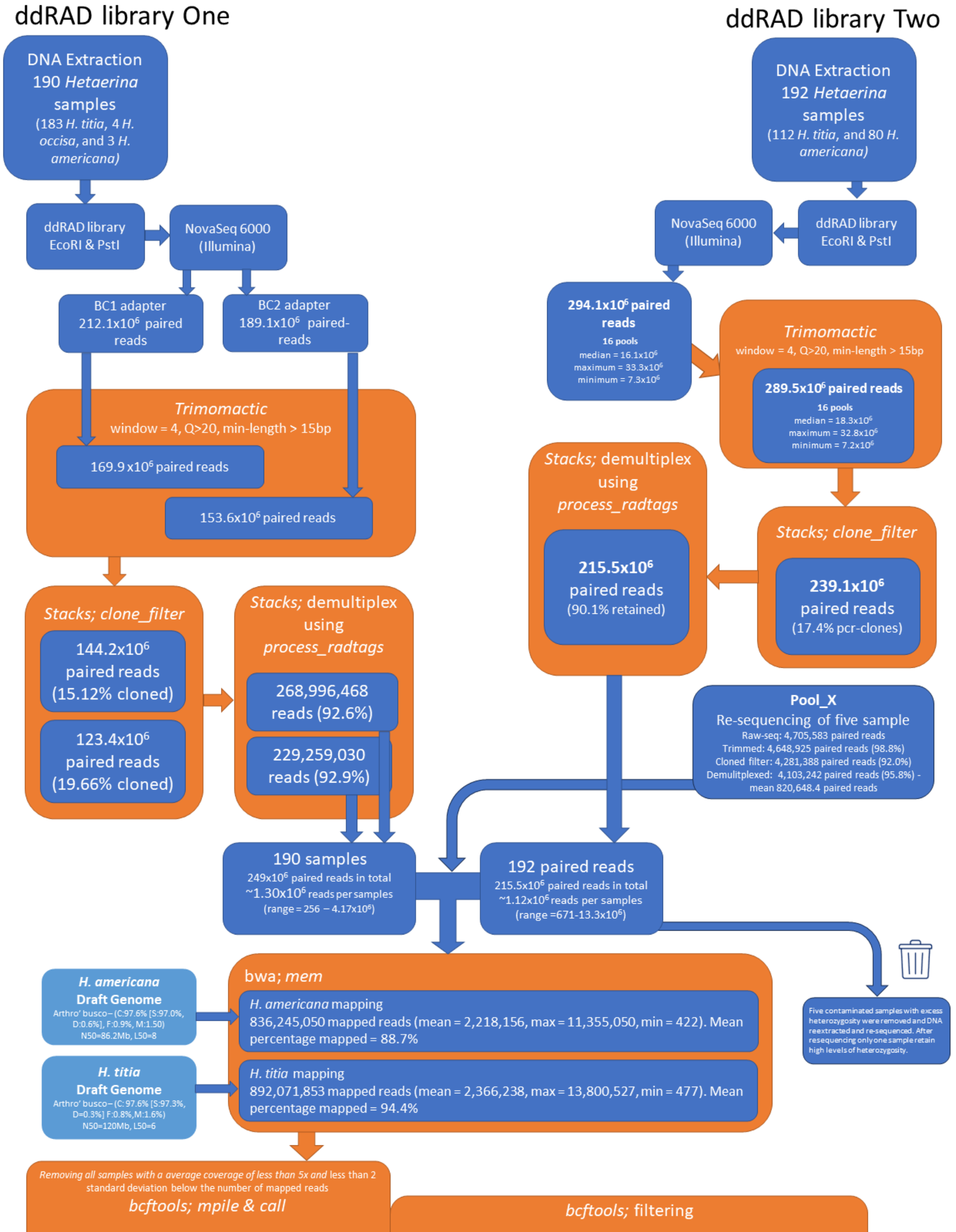


| Species | Draft genome | Number of samples | All variant calls | All single base calls | DP10-200, Q>20 | 80% Samples | MAF>0.05 | Random 1-1000 |
|---|---|---|---|---|---|---|---|---|
| **all** | *H. americana* | 364 | 134810308 | 134707986 | 2233411 | 80344 | 34506 | 1156 |
| **all** | *H. titia* | 363 | 161441042 | 161305711 | 2871466 | 81701 | 36376 | 1199 |
| ***H. americana/ calverti*** | *H. americana* | 75 | 73615352 | 73549147 | 4842870 | 1042635 | 151033 | 6997 |
| ***H. americana/ calverti*** | *H. titia* | 75 | 44189192 | 44164151 | 2258691 | 495629 | 80935 | 4103 |
| ***H. titia*** | *H. americana* | 286 | 81630947 | 81586704 | 1709783 | 260765 | 73471 | 3674 |
| ***H. titia*** | *H. titia* | 285 | 136427605 | 136309637 | 2997868 | 494012 | 121745 | 6369 |

Supplementary Figure 1: Flow diagram of bioinformatics pipeline for ddRAD libraries. ddRAD one conducted at the NERC environmental omics facility (NEOF) using the ddRAD protocol from DaCosta and Sorenson (2014). Library Two was conducted at Durham University following the methodology from Franchini *et al.* (2017). For increased clarity, the final bottom table is also presented in Supplementary Table 1.

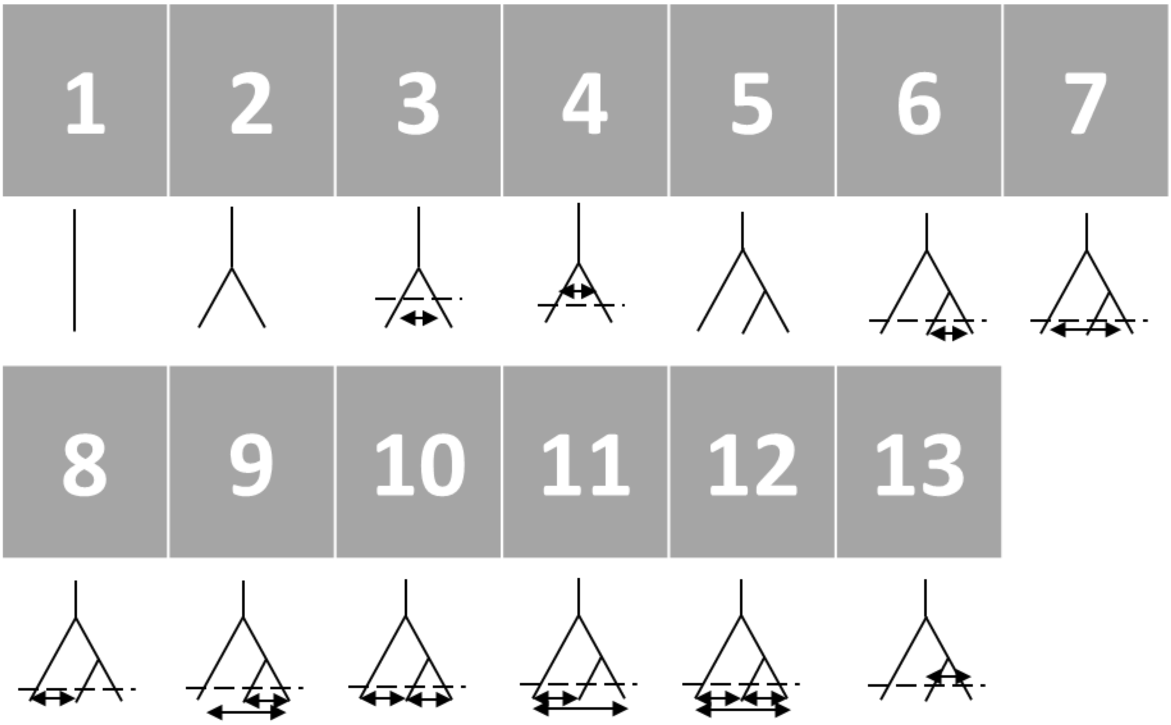


Supplementary Figure 2: Schematic diagrams of the thirteen demographic scenarios used in delimitR, We tested between all valid combination of the following demographic scenarios; one, two, or three potential lineages that arose either by pure isolation, isolation with gene flow, or isolation with secondary contact, and one to three migration bands between lineages. Arrows represent which lineages migration was included between in each demographic scenario. The dashed lines indicate a distinction between migration that occurred during (isolation with gene flow) or after (secondary contact) divergence, depending on whether the migration arrows are drawn above or below the dashed line respectively.

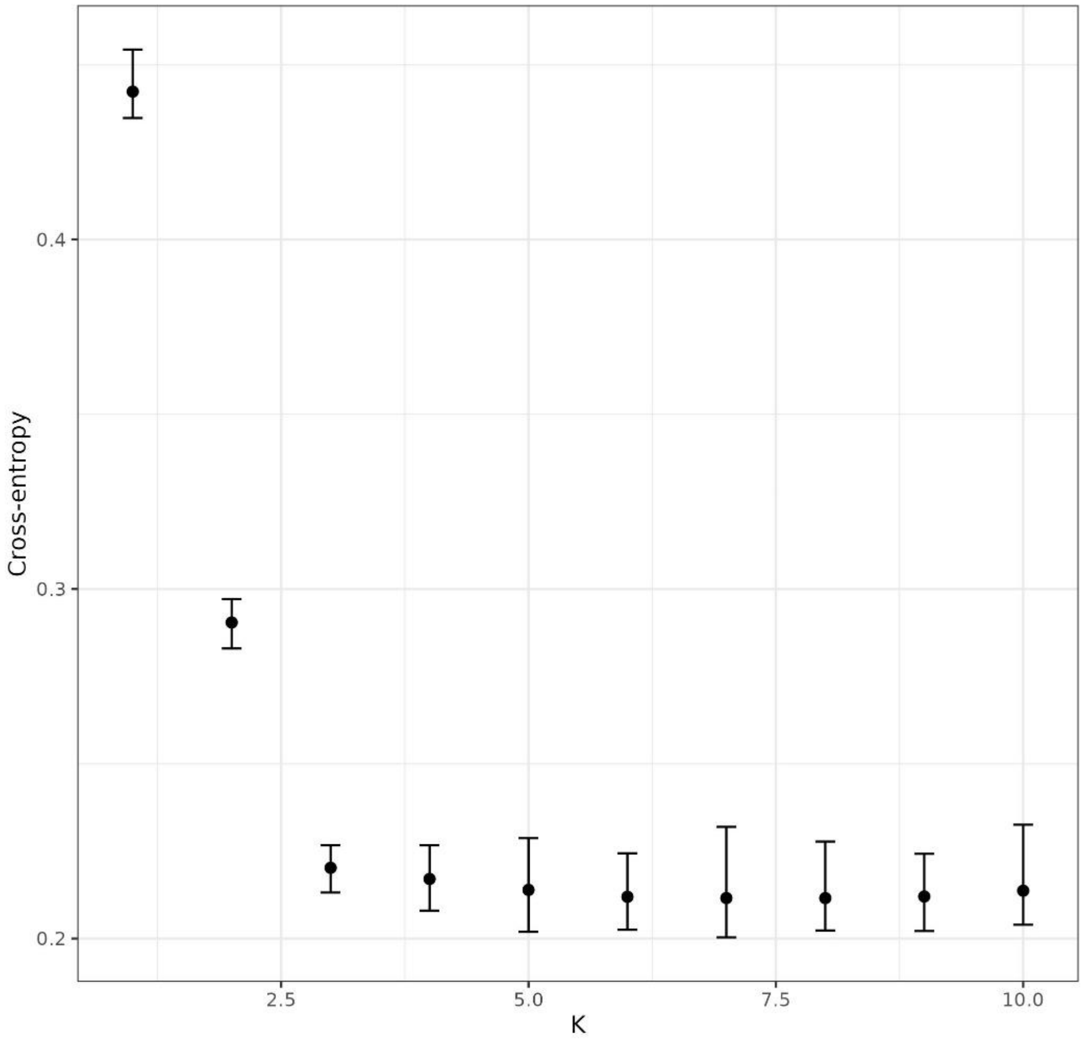


Supplementary Figure 3: The cross-entropy values from K = 1-10 and with error bars showing the maximum and minimum values from 100 repetitions of snmf analysis. Produced using the SNP library of *Hetaerina titia* samples mapped to the *H. titia* genome HetTit1.0 (Patterson et al., 2023)

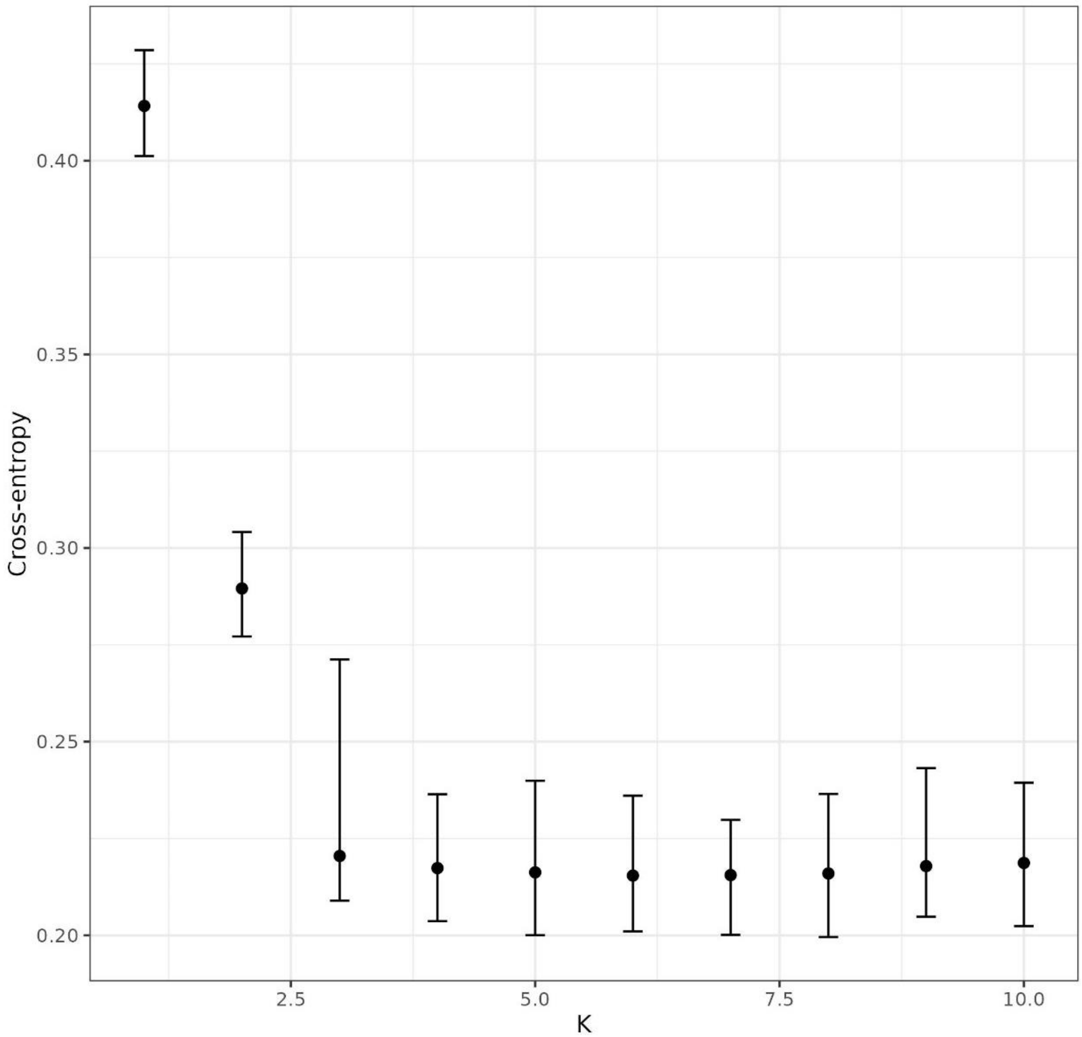


Supplementary Figure 4: The cross-entropy values from K = 1-10 and with error bars showing the maximum and minimum values from 100 repetitions of snmf analysis. Produced using the SNP library of *Hetaerina titia* samples mapped to the *H. americana* genome HetAmer1.0 (Grether et al., 2023)

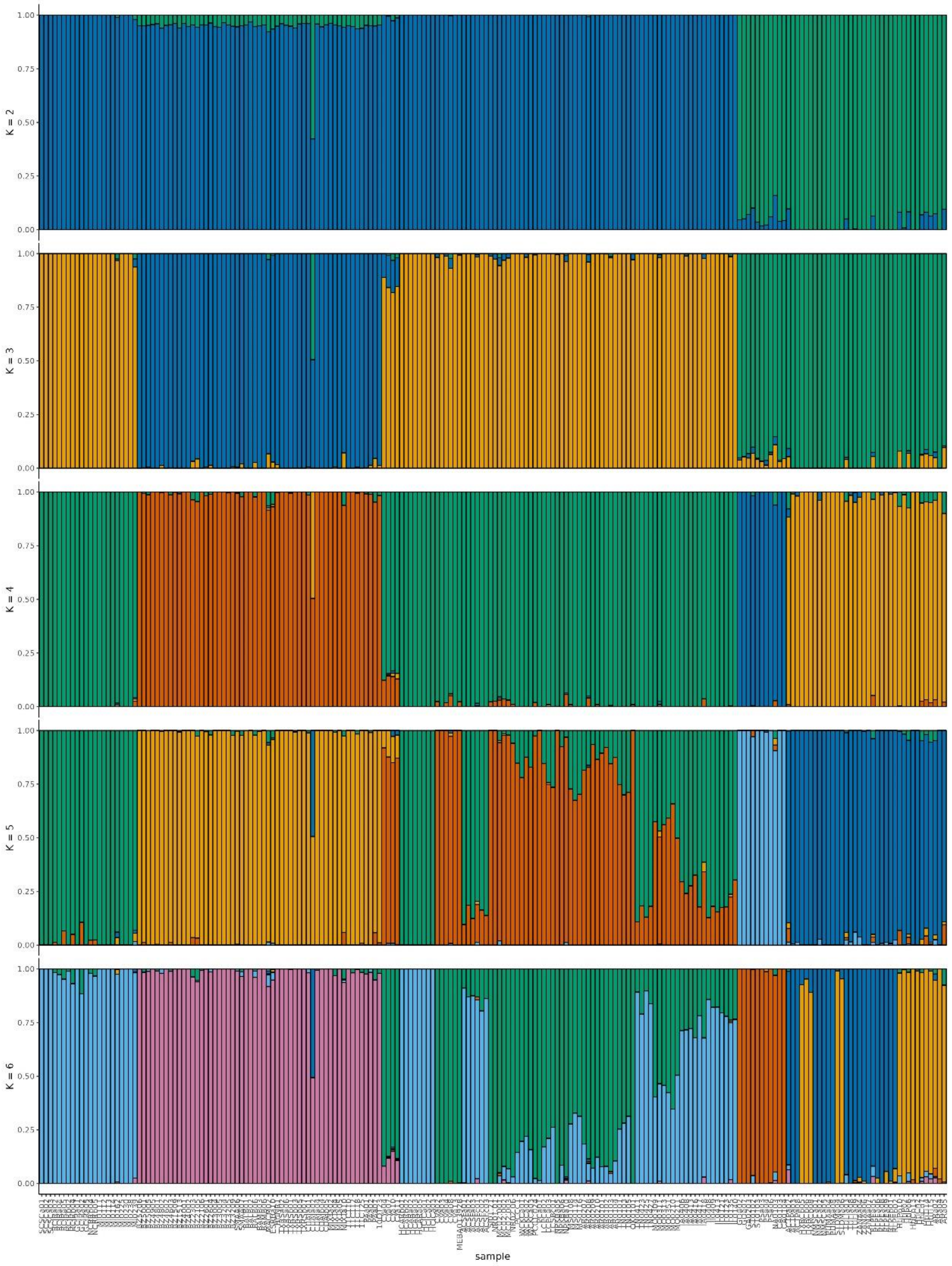


Supplementary Figure 5: Estimate of ancestry analysis for each individual using sNMF using K = 2-6 for all *Hetaerina titia* samples using the *Hetaerina titia* draft genome HetTit1.0 from (Patterson et al., 2023) Each value of K was run for 100 times with an alpha value of 100. Samples are ordered by drainage, then country, and then latitude.

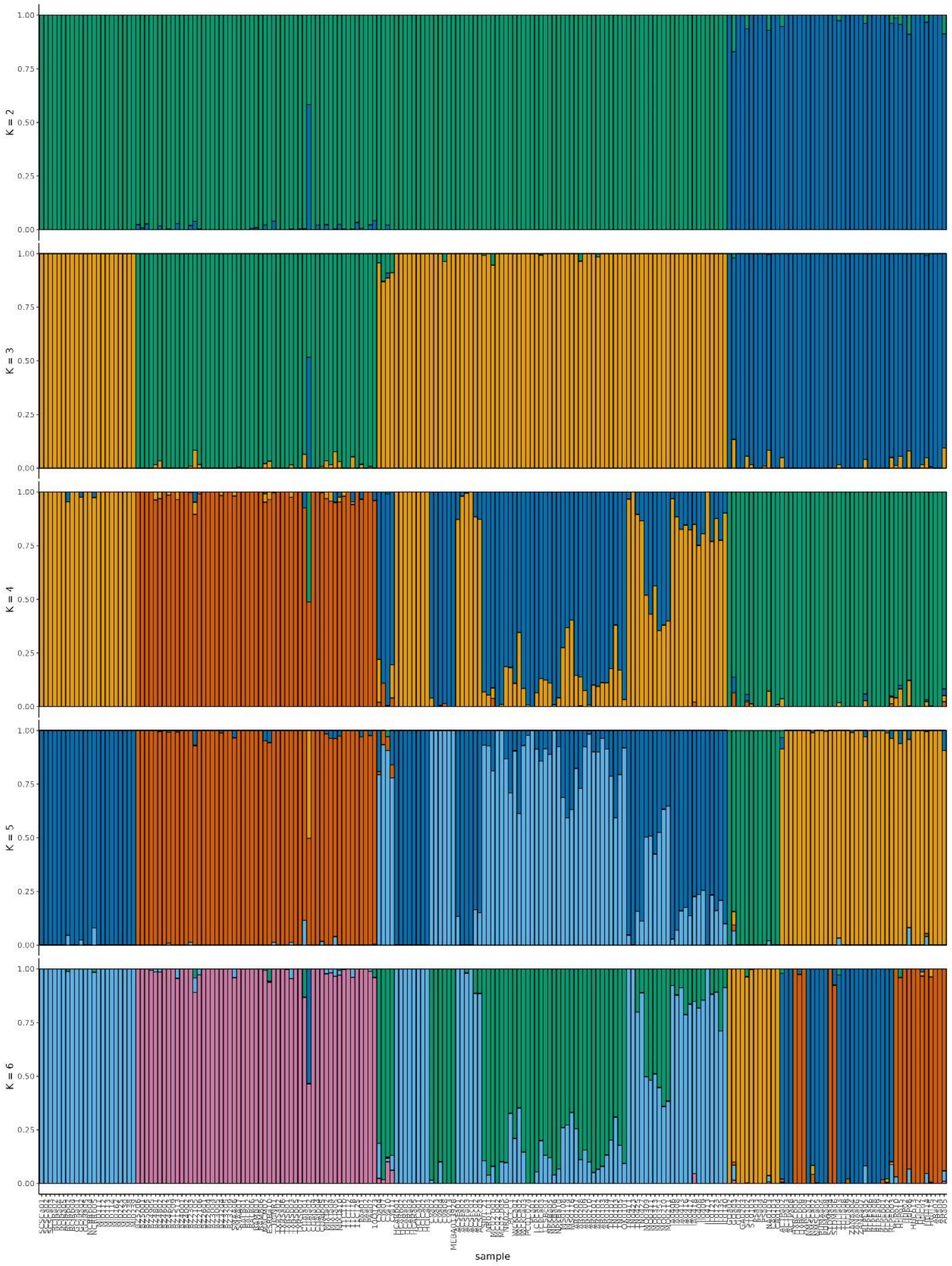


Supplementary Figure 6 :Estimate of ancestry analysis for each individual using sNMF using K = 2-6 for all *Hetaerina titia* samples using the *Hetaerina americana* draft genome HetAmer1.0 from (Grether et al., 2023) Each value of K was run for 100 times with an alpha value of 100.

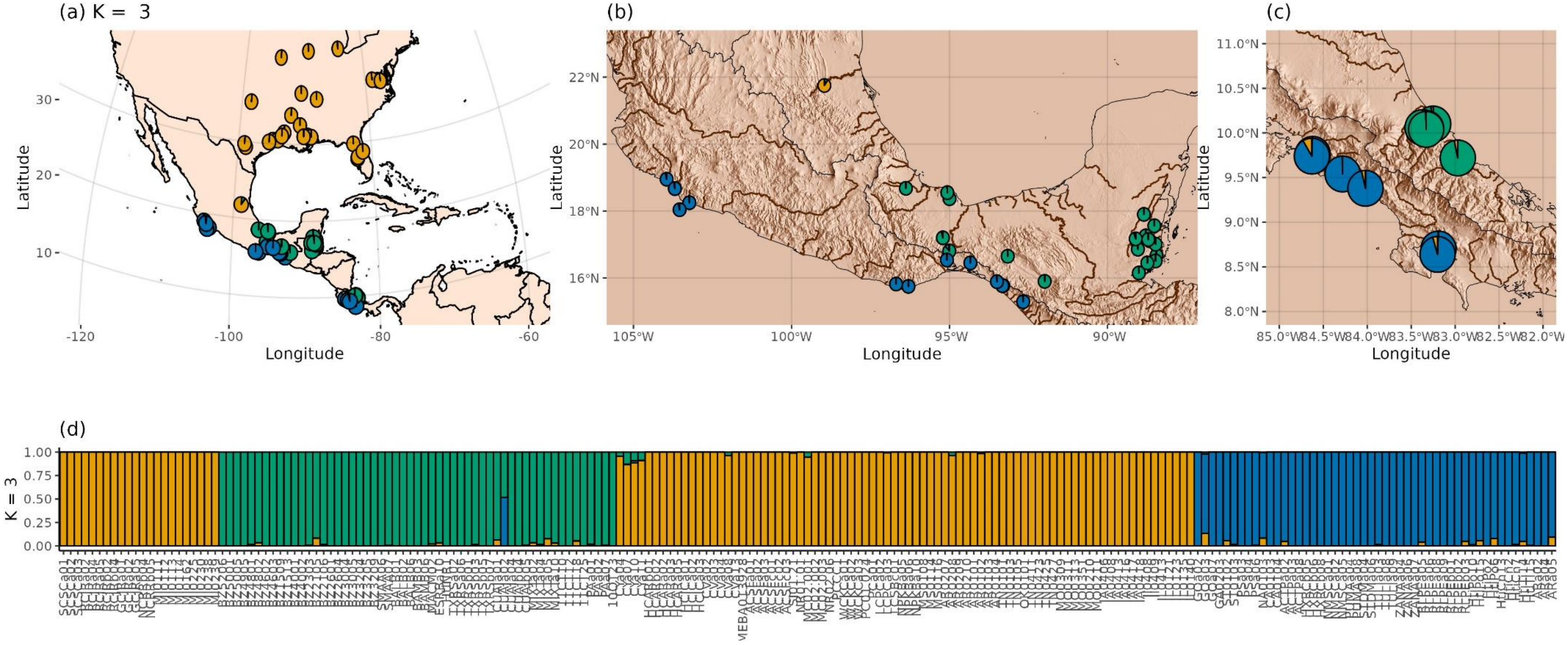


Supplementary Figure 7: Ancestry estimates using three ancestral populations for *Hetaerina titia*. SNPs were generated by mapping ddRAD reads to the draft genome of *H. americana.* This is in comparison to the Figure 1 in the main text which used the reads mapped to the draft genome of *H. titia*. LEA was run for 20 repetitions and an alpha value of 100. (a) The mean estimate of ancestry proportion for all samples within each sample site of *Hetaerina titia* across North and Central America, (b) Isthmus of Tehuantepec and Belize, and (c) Costa Rica. (d) Estimate of ancestry analysis for each individual. Samples are ordered by drainage, then country, and then latitude. Rivers and drainage basins from Hydrosheds. Topography data from the R package elevatr.

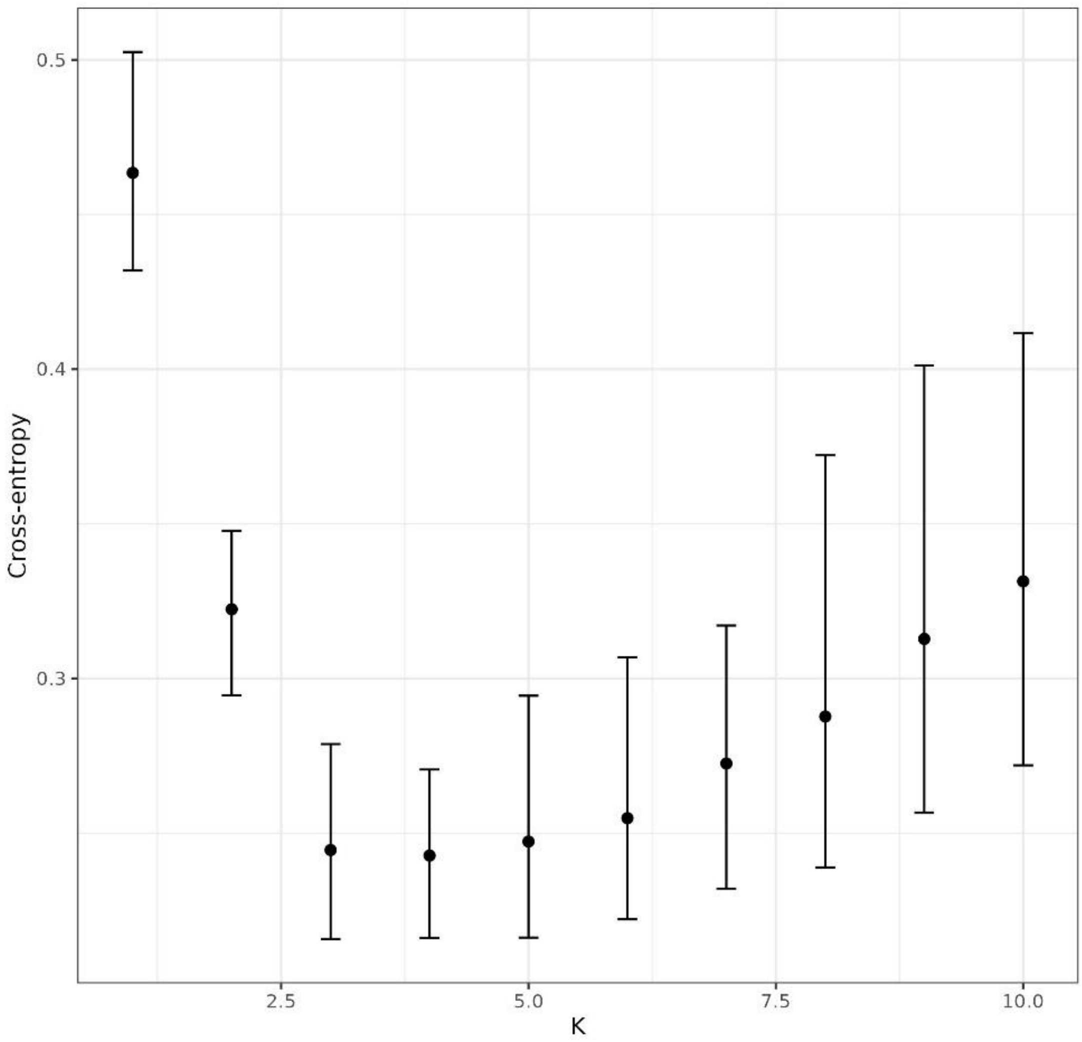


Supplementary Figure 8: The cross-entropy values from K = 1-10 and with error bars showing the maximum and minimum values from 100 repetitions of snmf analysis. Produced using the SNP library of *Hetaerina americana* sensu lato samples mapped to the *H. titia* genome HetTit1.0 (Patterson et al., 2023)

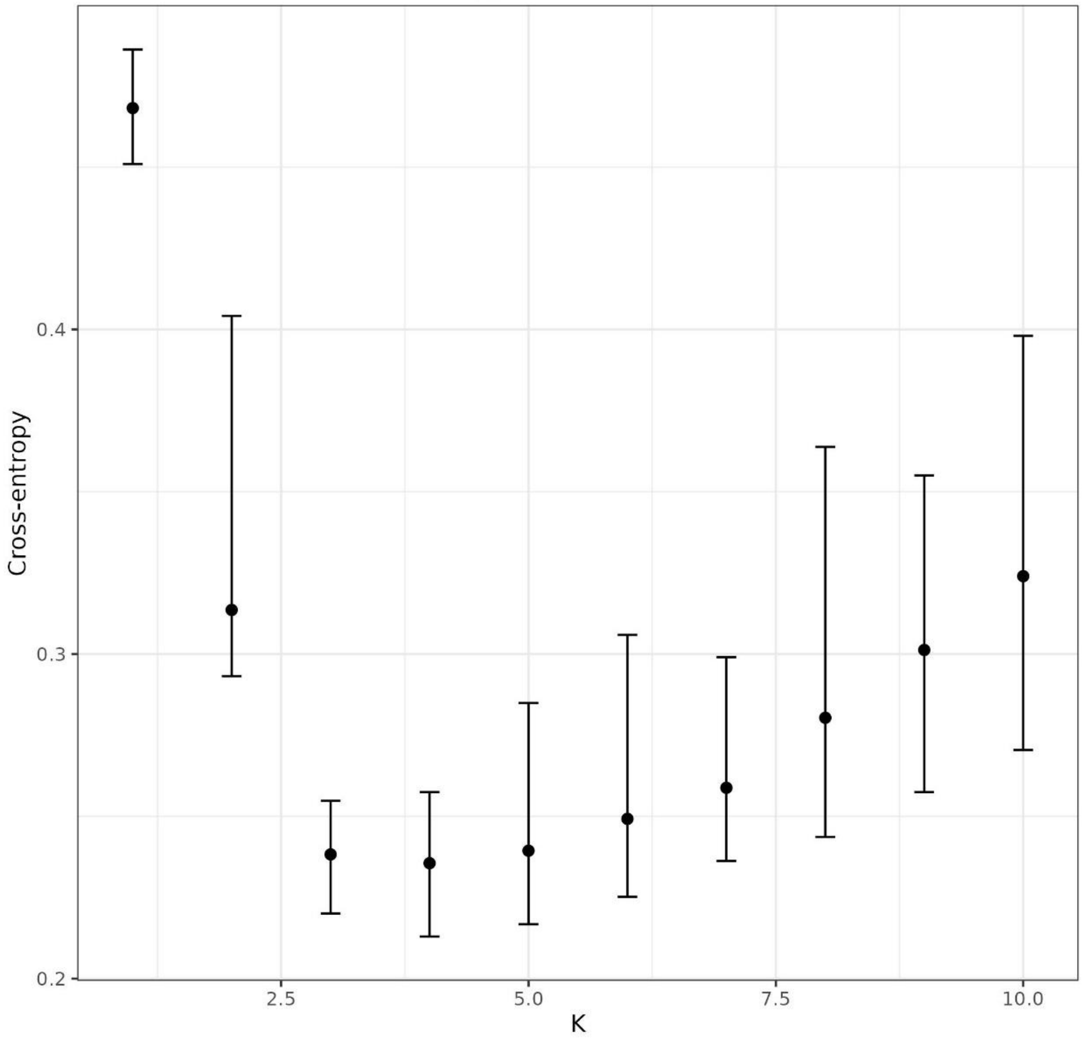

Supplementary Figure 9: The cross-entropy values from K = 1-10 and with error bars showing the maximum and minimum values from 100 repetitions of snmf analysis. Produced using the SNP library of *Hetaerina americana* sensu lato samples mapped to the *H. americana* genome HetAmer1.0 (Grether et al., 2023).

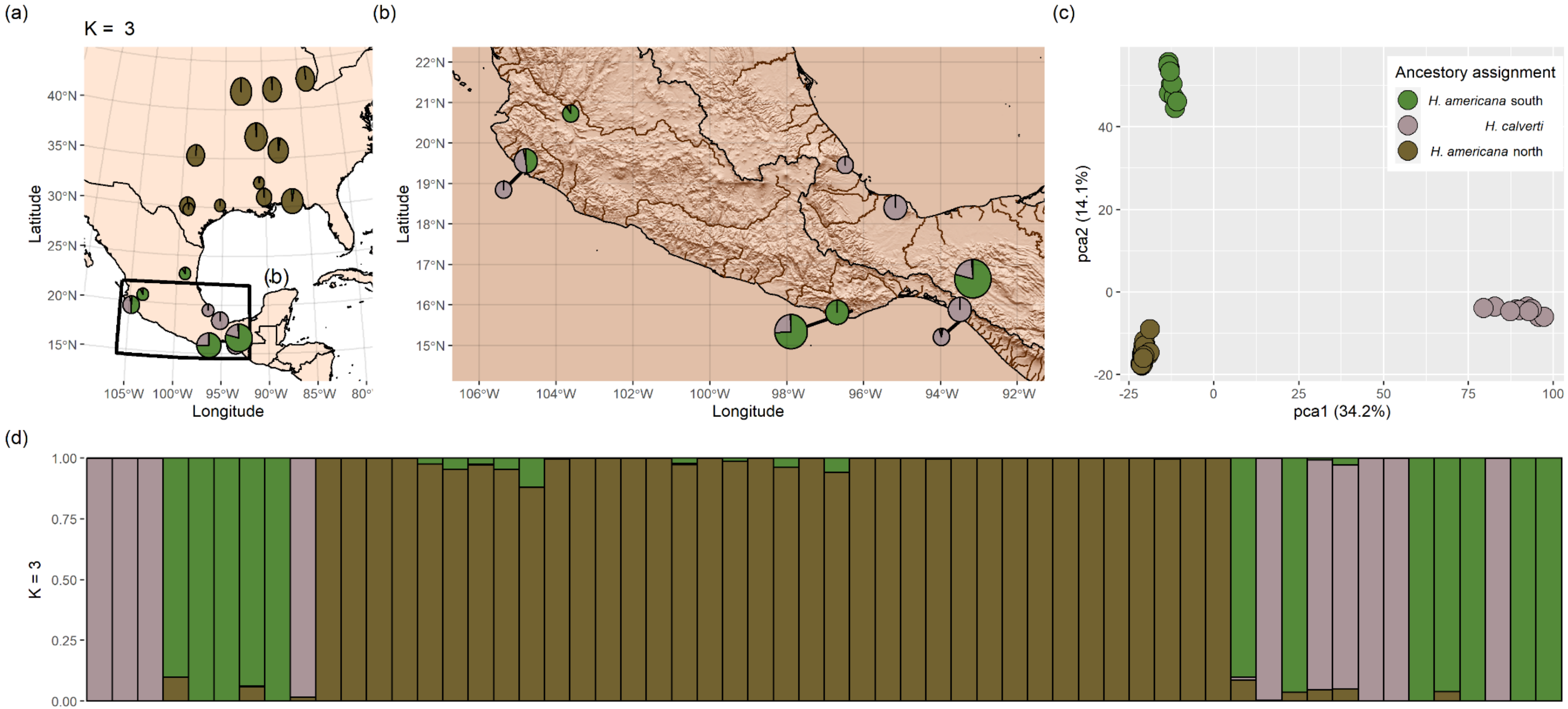


Supplementary Figure 10: Ancestry estimates using three ancestral populations for 58 *Hetaerina americana* (green and gold) and *H. calverti* (light grey) with a dataset of 5,259 unlinked biallelic autosomal SNPs, 20 repetitions and an alpha value of 100. SNPs where generated from reads mapped to the draft genome of *H. titia* **(a, b)** The mean estimate of ancestry proportion for all samples within each sample site of *H. americana/calverti*. **(c)** Principal component analysis of the same dataset with sample colour indicating the highest assigned ancestry population from sNMF for each individual. **(d)** Estimate of ancestry for each sample. Samples are ordered by drainage, then country, and then latitude. Rivers and drainage basins from Hydrosheds. Topography data downloaded using R package *elevatr*.

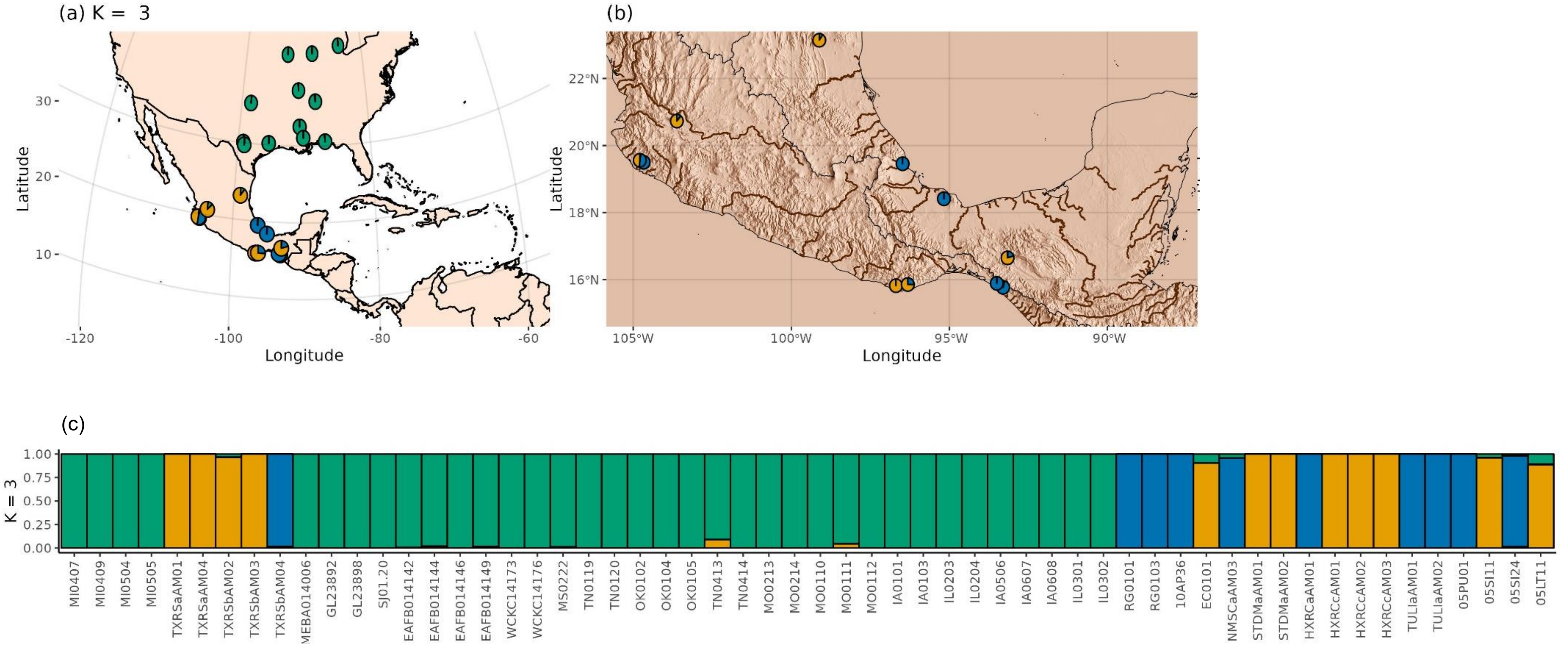


Supplementary Figure 11: Ancestry estimates using three ancestral populations for *Hetaerina americana* sensu lato. SNPs were generated by mapping ddRAD reads to the draft genome of *H. americana*. LEA was run for 20 repetitions and an alpha value of 100. (a) The mean estimate of ancestry proportion for all samples within each sample site of *Hetaerina americana* sensu lato across North and Central America, and (b) Isthmus of Tehuantepec and Belize. (c) Estimate of ancestry analysis for each individual. Samples are ordered by drainage, then country, and then latitude. Rivers and drainage basins from Hydrosheds. Topography data from the R package elevatr

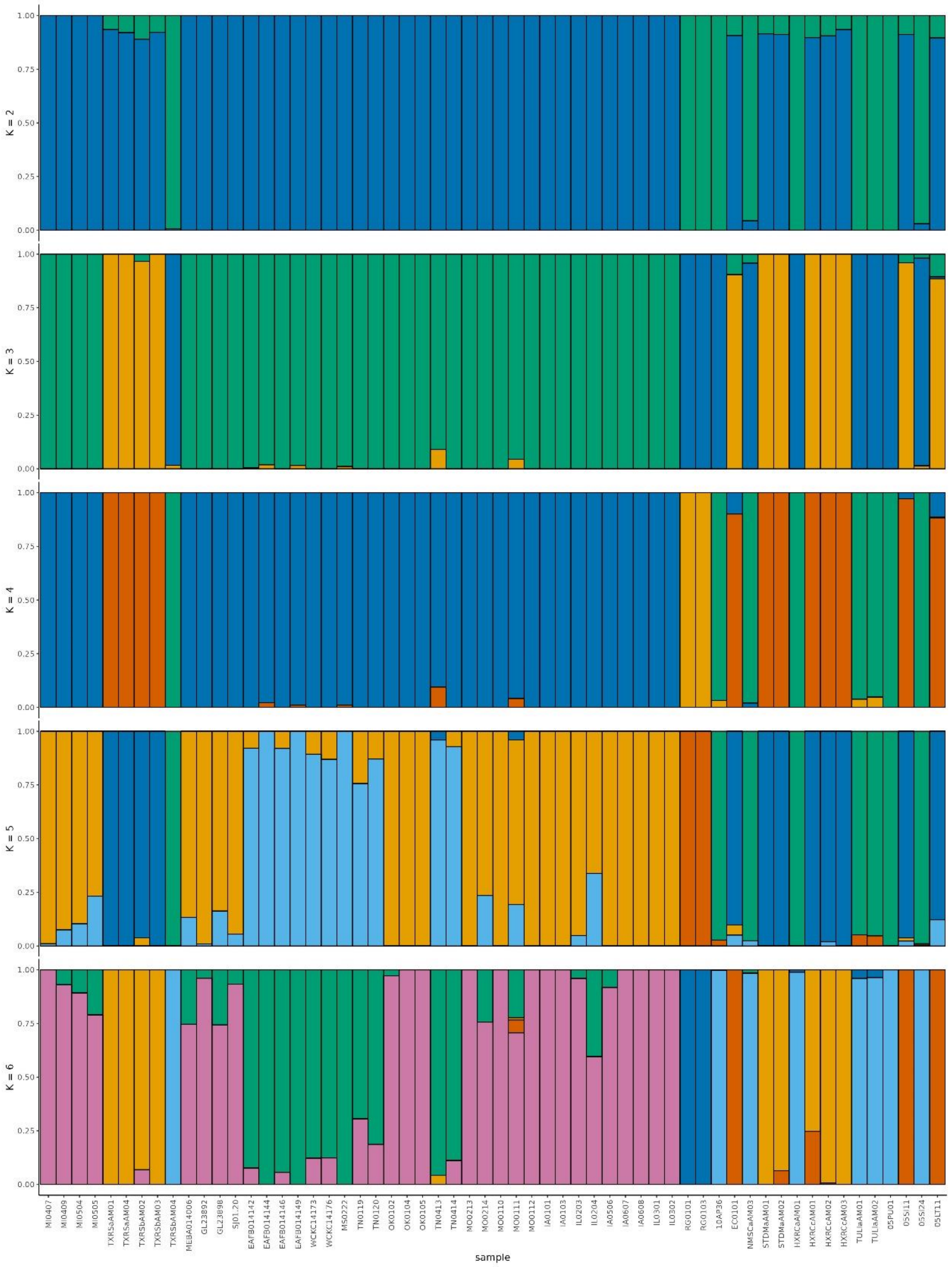


Supplementary Figure 12: Estimate of ancestry analysis using sNMF using K = 2-6 for all *Hetaerina americana* sensu lato samples using the *Hetaerina titia* draft genome HetTit1.0 from (Patterson et al., 2023). Each value of K was run for 100 times with an alpha value of 100.

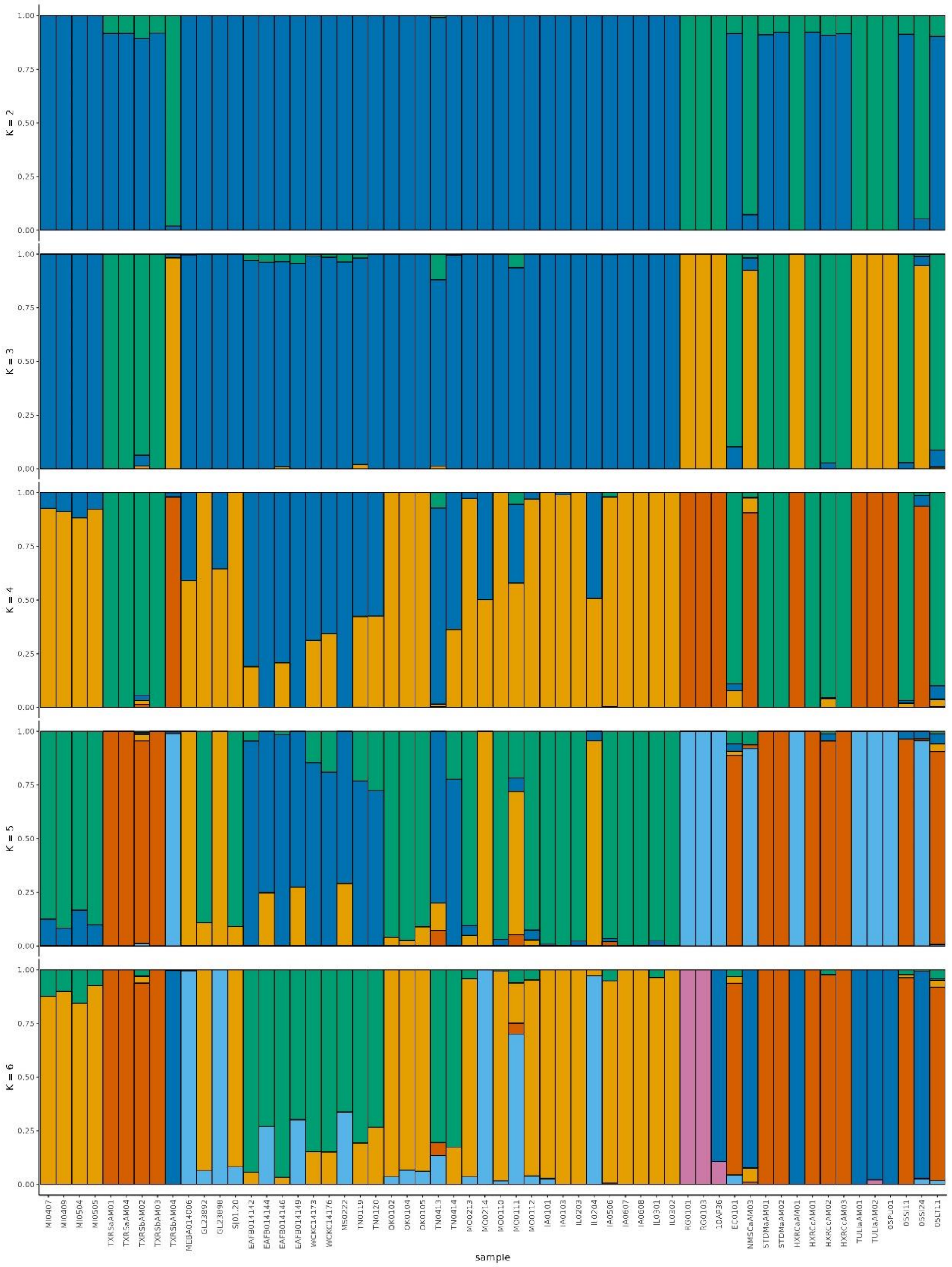


Supplementary Figure 13: Estimate of ancestry analysis using sNMF using K = 2-6 for all *Hetaerina americana* sensu lato samples using the *Hetaerina americana* draft genome HetAmer1.0 from (Grether et al., 2023) Each value of K was run for 100 times with an alpha value of 100.

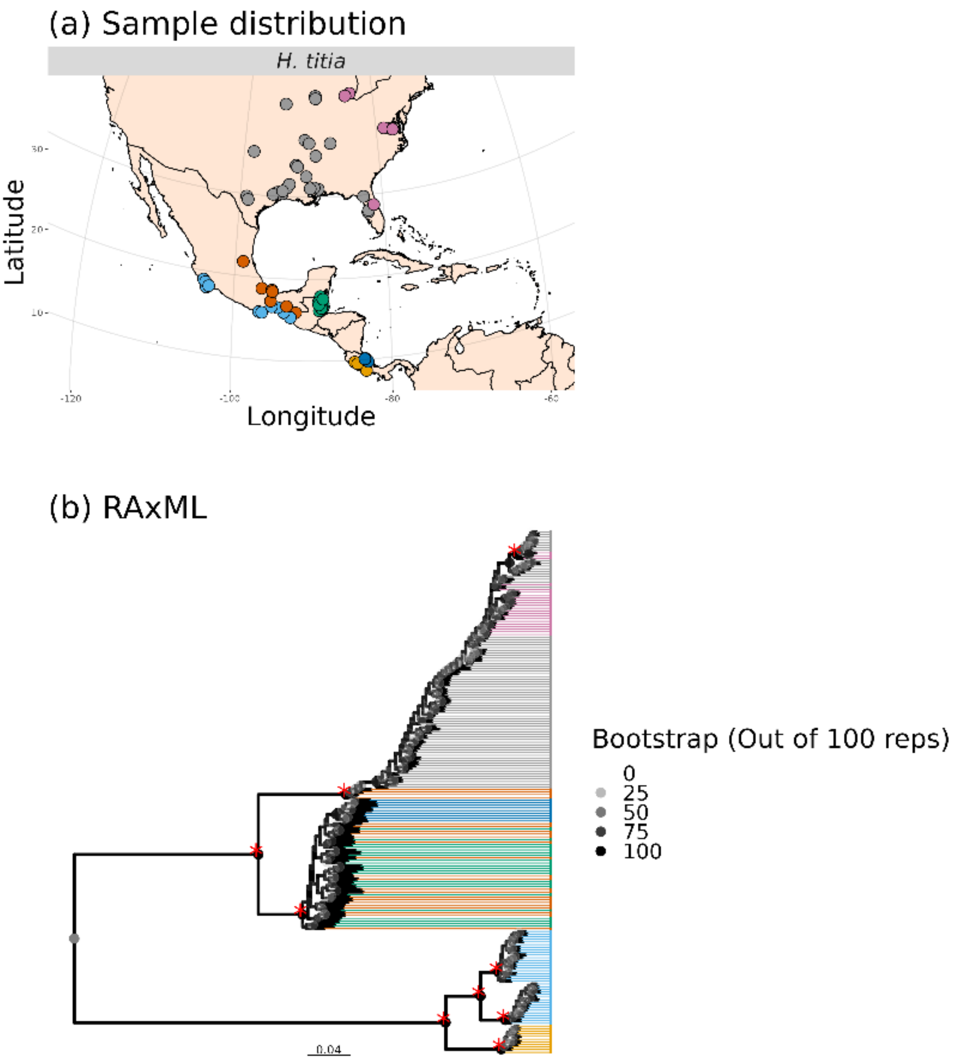


Supplementary Figure 14: The maximum likelihood tree for *Hetaerina titia* calculated by RAxML using a SNP library mapped onto the genome of *H. titia*. The tree is rooted around the midpoint which corresponds the root identified by the multi-species tree. The nodes are coloured by the number of bootstrap support values (out of 100). Nodes with greater than 95% support are marked with a red “*”. The tree tips are coloured according to the species, the country, and river drainage of each sample shown in the inset map. Scale bar indicates number of substitutions per SNP site.

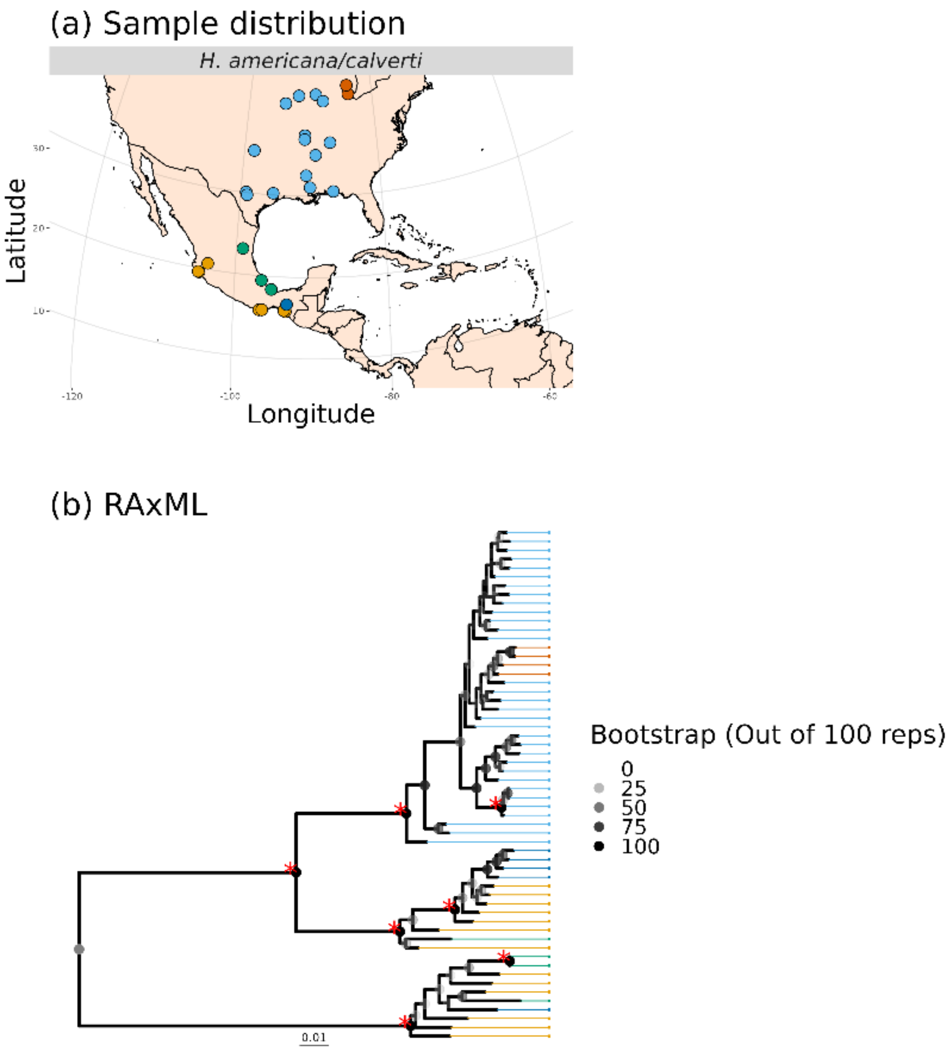


Supplementary Figure 15: The maximum likelihood tree for *Hetaerina americana and Hetaerina calverti* calculated by RAxML using a SNP library mapped onto the genome of *H. titia*. The tree is rooted around the midpoint which corresponds the root identified by the multi-species tree. The nodes are coloured by the number of bootstrap support values (out of 100). Nodes with greater than 95% support are marked with a red "*". The tree tips are coloured according to the species, the country, and river drainage of each sample shown in the inset map. The colour of each tip and geographical position of each sample does not separate *Hetaerina americana* and *Hetaerina calverti.* Scale bar indicates number of substitutions per SNP site.

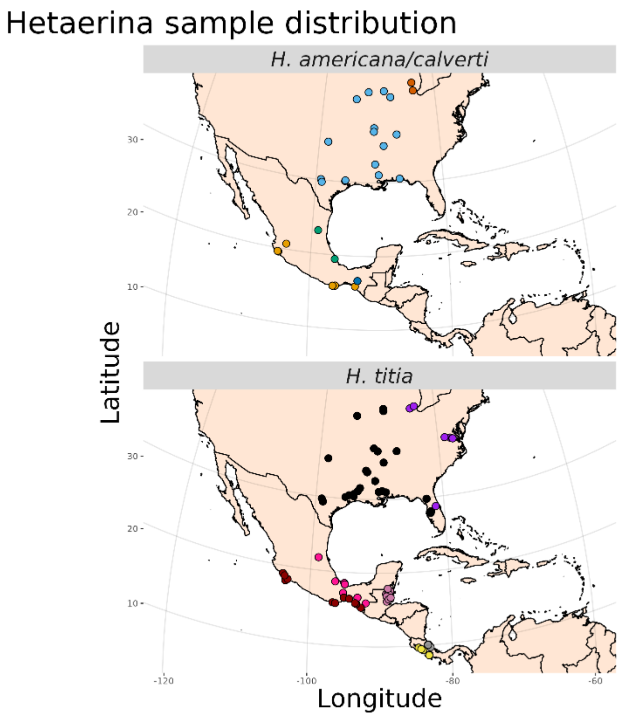


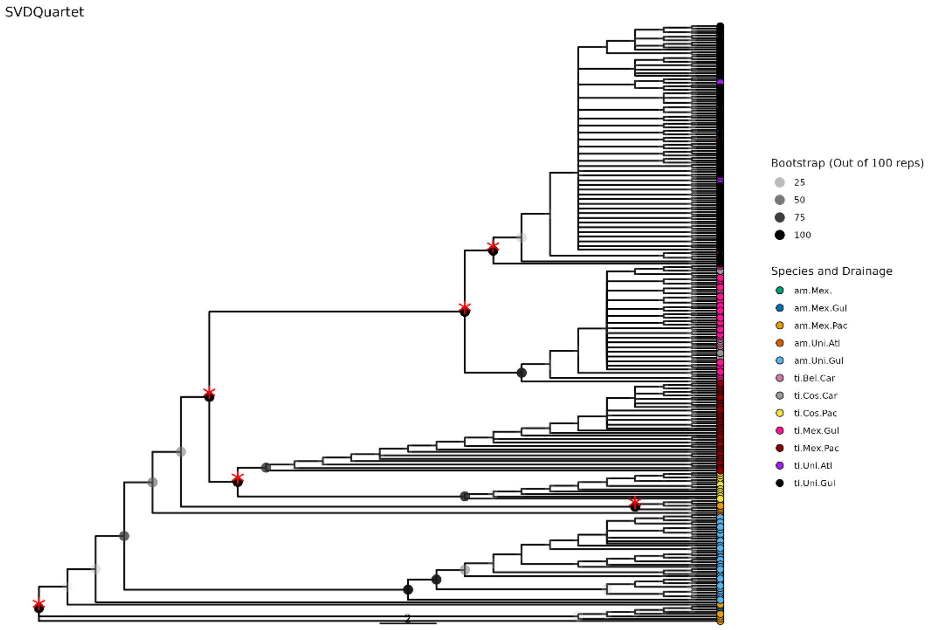


Supplementary Figure 16: SVDquartet analysis of *H. tita, H. americana* and *H. calverti* mapped to the draft genome of H. titia. Node with greater than 95% bootstrap support indicated with a read "*".

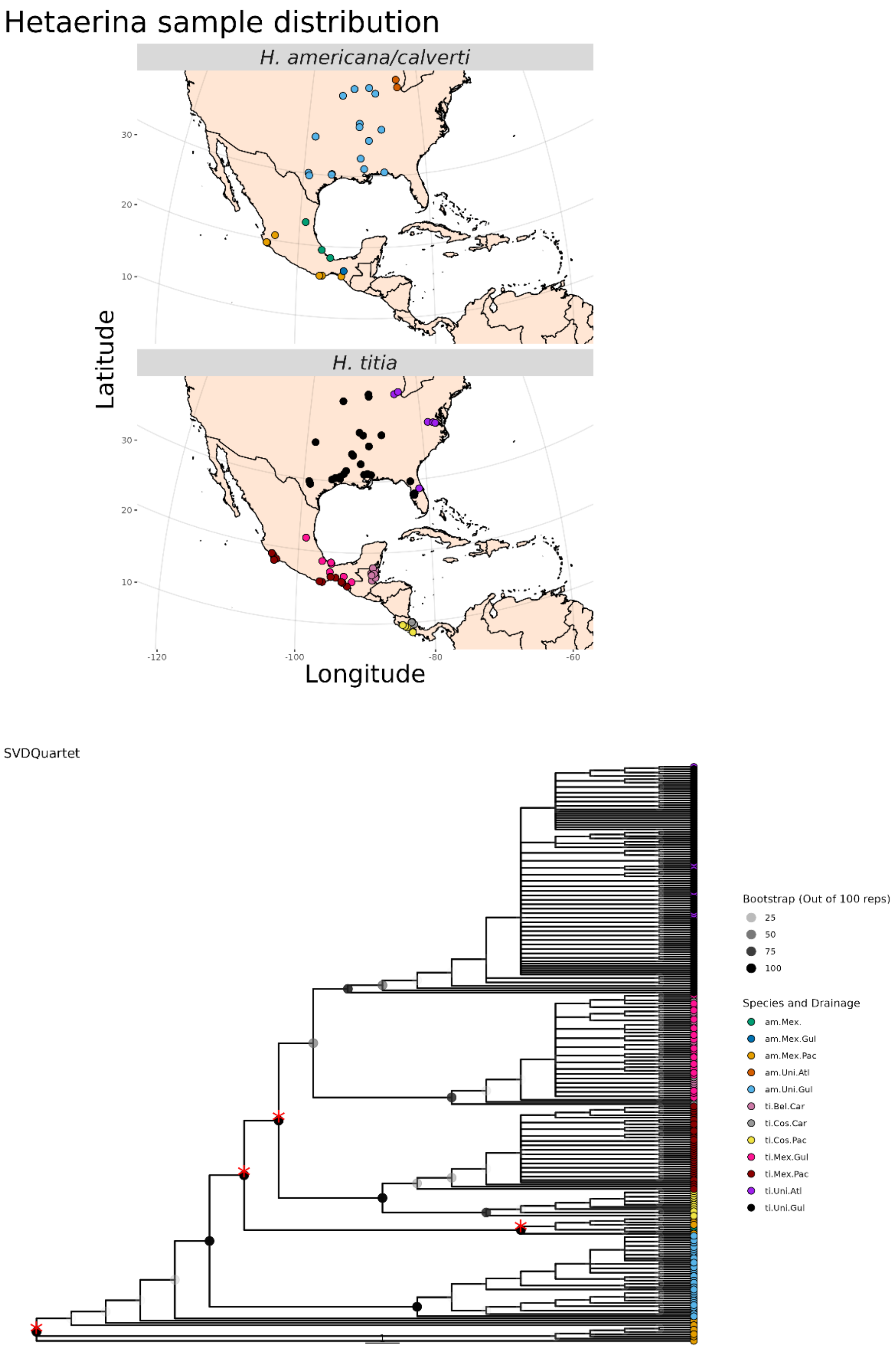


Supplementary Figure 17: SVDquartet analysis of *H. tita, H. americana* and *H. calverti* mapped to the draft genome of *H. americana.* Node with greater than 95% bootstrap support indicated with a read "*".

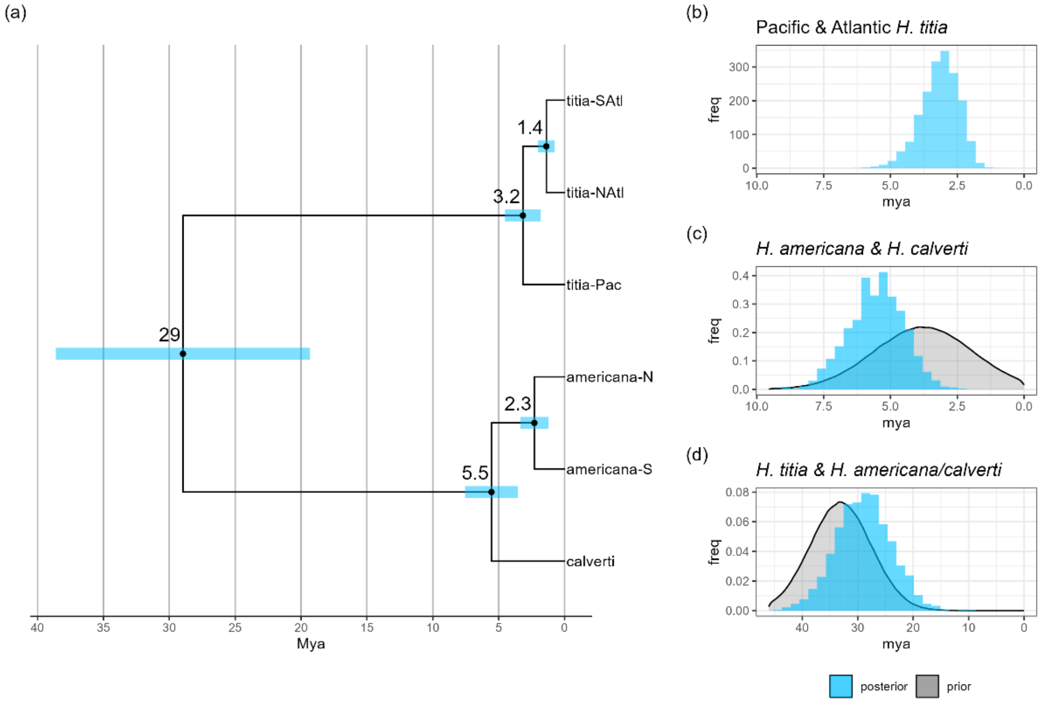


Supplementary Figure 18: Estimates of divergence dates between populations of *Hetaerina titia* (titia-NAtl, titia-SAlt, and titia-Pac), *Hetaerina americana* (americana-N americana-S), and *Hetaerina calverti* (calverti) calculated using SNAPP analysis in beast. Input data was 540 SNPs with 4 samples per tip, mapped to the genome of *Hetaerina americana*. Node labels indicate the mean estimated divergence date with 95% highest posterior density in blue. All branches had a posterior distribution of 1.

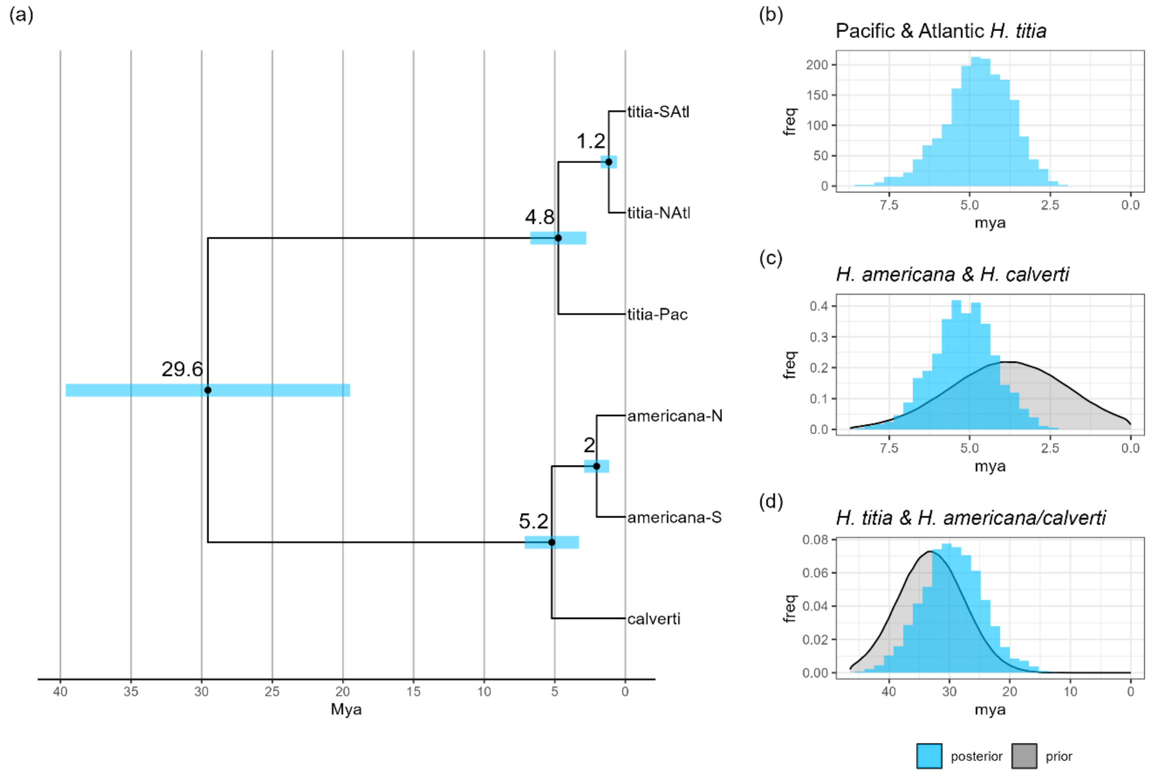


Supplementary Figure 19: Estimates of divergence dates between populations of *Hetaerina titia* (titia-NAtl, titia-SAlt, and titia-Pac), *Hetaerina americana* (americana-N americana-S), and *Hetaerina calverti* (calverti) calculated using SNAPP analysis in beast. Input data was 552 SNPs with four samples per tip, mapped to the genome of *Hetaerina titia*. Node labels indicate the mean estimated divergence date with 95% highest posterior density in blue. All branches had a posterior distribution of 1.

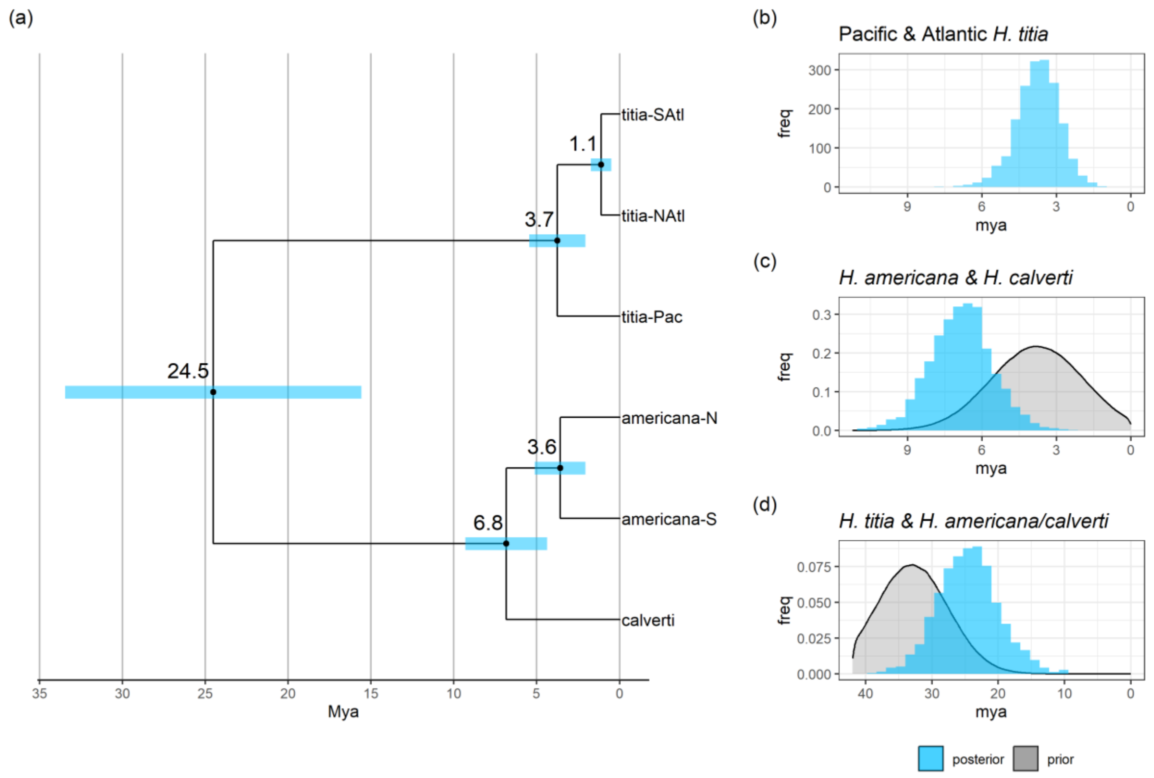


Supplementary Figure 20: Estimates of divergence dates between populations of *Hetaerina titia* (titia-NAtl, titia-SAlt, and titia-Pac), *Hetaerina americana* (americana-N americana-S), and *Hetaerina calverti* (calverti) calculated using SNAPP analysis in beast. Input data was 519 SNPs with three samples per tip, mapped *de novo* using ipyrad. Node labels indicate the mean estimated divergence date with 95% highest posterior density in blue. All branches had a posterior distribution of 1.

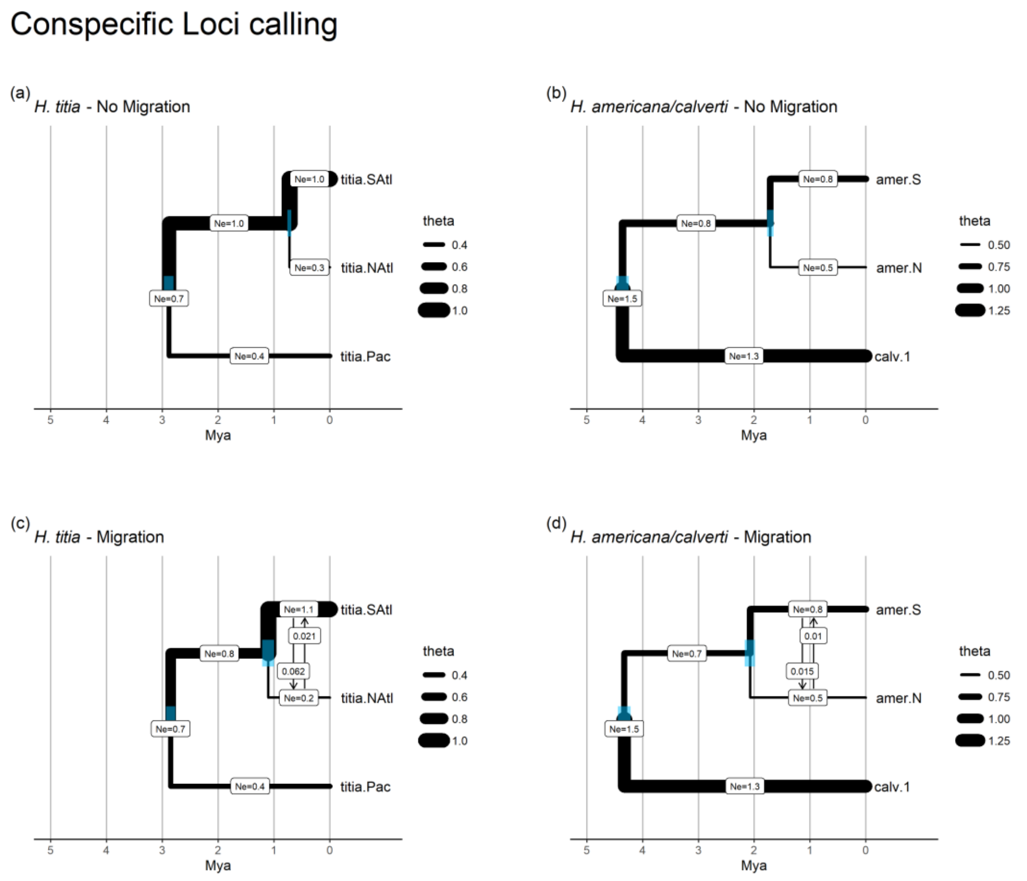


Supplementary Figure 21: The estimated divergence times (million years ago) and effective population size (theta – N*e* in millions of individuals) from G-PhoCS analysis of *H. titia* and *H. americana.* Migration rate is shown as the number of individuals per generation. All models ran for 1,000,000 interactions with 10% burn in. Blue bars show 95% highest posterior density for each divergence date **(a)** Model estimates for *H. titia* with no migration bands. **(b)** Model estimates for *H. americana* and *H. calverti* with no migration bands. **(c)** Model estimates for *H. titia* demography with migration bands between Northern and Southern Atlantic *H. titia* **(d)** Model estimates for *H. americana* and *H. calverti* with migration bands between North and Southern *H. americana*. G-PhoCS runs presented here are conducted on the conspecific draft genome. i.e., *H. titia* loci mapped to the *H. titia* draft genome and the *H. americana* loci mapped to the *H. americana* draft genome.

### Heterospecific Loci calling

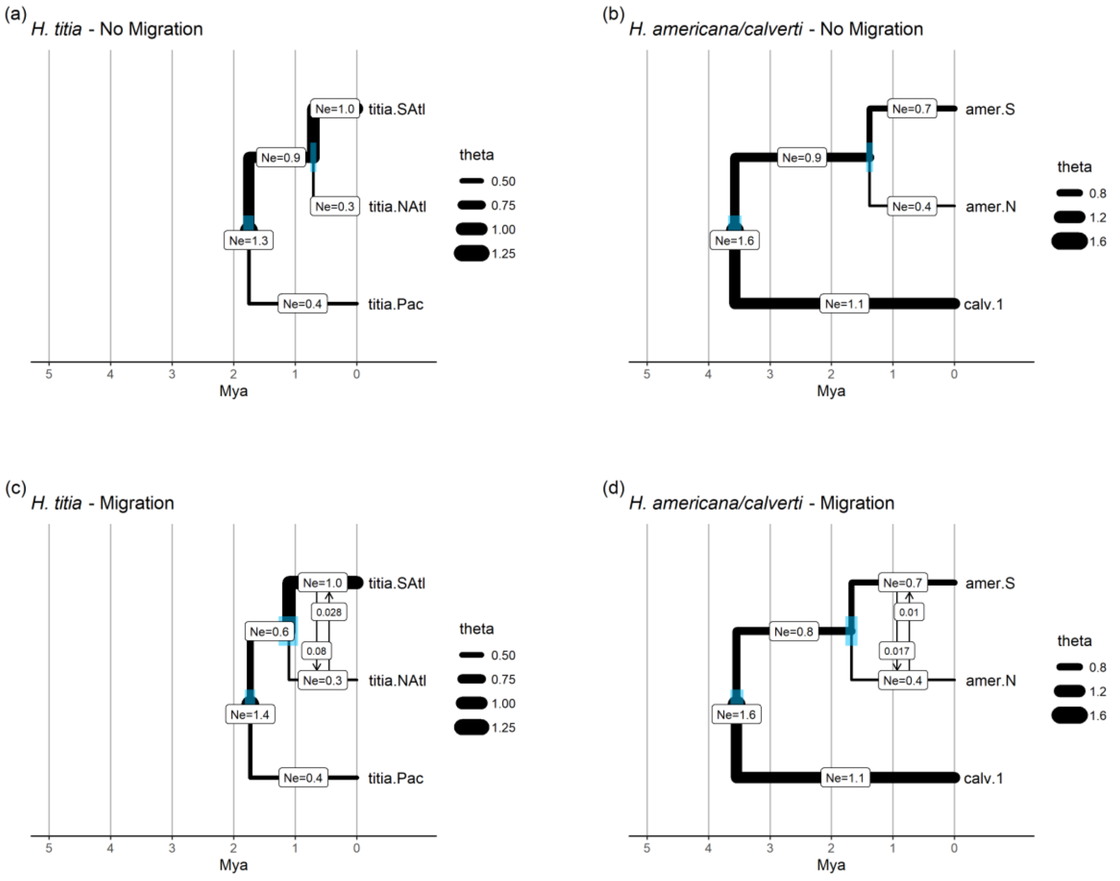


Supplementary Figure 22: The estimated divergence times (million years ago) and effective population size (theta – N*e* in millions of individuals) from G-PhoCS analysis of *H. titia* and *H. americana.* Migration rate is shown as the number of individuals per generation. All models ran for 1,000,000 interactions with 10% burn in. Blue bars show 95% highest posterior density for each divergence date **(a)** Model estimates for *H. titia* with no migration bands. **(b)** Model estimates for *H. americana* and *H. calverti* with no migration bands. **(c)** Model estimates for *H. titia* demography with migration bands between Northern and Southern Atlantic *H. titia* **(d)** Model estimates for *H. americana* and *H. calverti* with migration bands between North and Southern *H. americana*. G-PhoCS runs presented here are conducted on the heterospecific draft genome. i.e., *H. titia* loci mapped to the *H. americana* draft genome and the *H. americana* loci mapped to the *H. titia* draft genome.

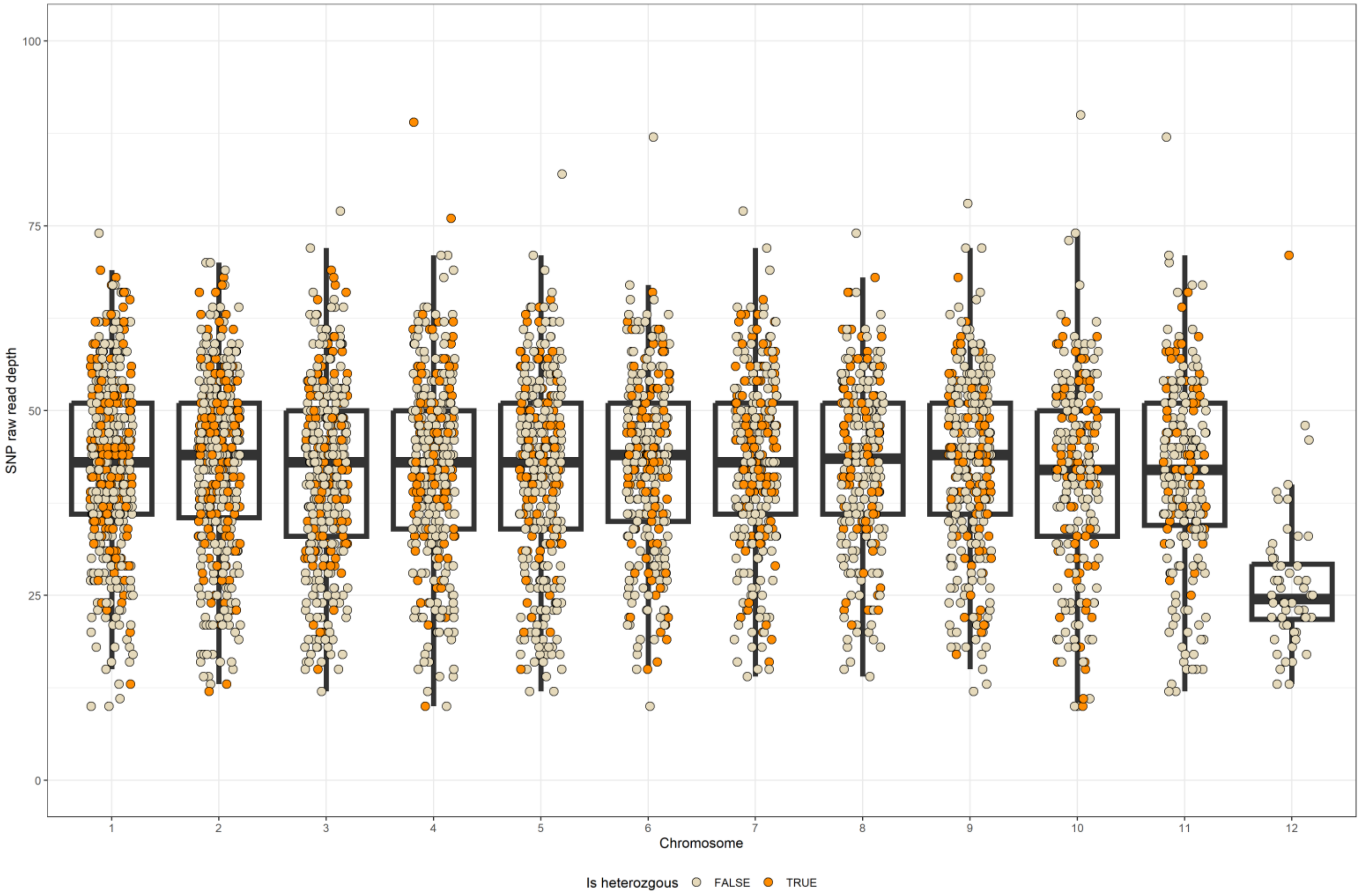


Supplementary Figure 23: All genotyped SNPs from individual CUAJa02 from site CUAJ01, Cuajinicuil, Oaxaca (16° 47'24.00"N, 95°0'36.00"W). Each point marks the raw read number used for SNP calling at that specific site, which chromosome the SNP was mapped to and whether site was heterozygous. Chromosome 12 is the X chromosome. The y axis is restricted to a read coverage of 0 to 100 excluding a small number of high coverage autosomal SNPs.

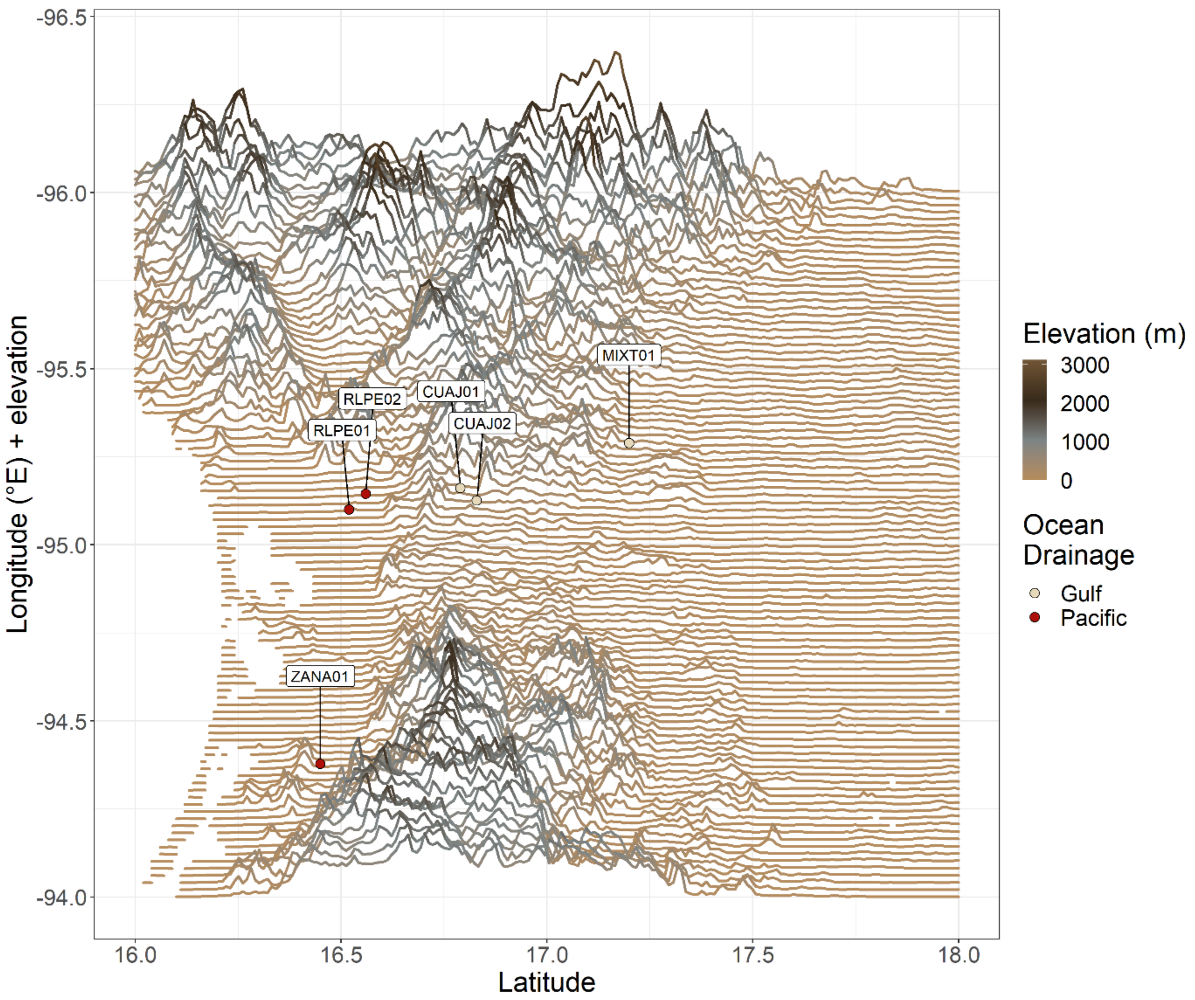


Supplementary Figure 24: Elevational transects running East to West for the Isthmus of Tehuantepec. Each transect is plotted as the longitude of the transect plus the elevation, at each latitude, using the equation (Longitude – (Elevation/max_elevation) x 0.4). Sample sites are marked and transformed to be in line with the elevational transects. The $F_1$ hybrid was collected in an Atlantic river basin at site CUAJ01.
